# Patterns of diverse gene functions in genomic neighborhoods predict gene function and phenotype

**DOI:** 10.1101/582577

**Authors:** Matej Mihelčić, Tomislav Šmuc, Fran Supek

## Abstract

Genes with similar roles in the cell are known to cluster on chromosomes, thus benefiting from coordinated regulation. This allows gene function to be inferred by transferring annotations from genomic neighbors, following the guilt-by-association principle. We performed a systematic search for co-occurrence of >1000 gene functions in genomic neighborhoods across 1669 prokaryotic, 49 fungal and 80 metazoan genomes, revealing prevalent patterns that cannot be explained by clustering of functionally similar genes. It is a very common occurrence that pairs of dissimilar gene functions – corresponding to semantically distant Gene Ontology terms – are significantly co-located on chromosomes. These neighborhood associations are often as conserved across genomes as the known associations between similar functions, suggesting selective benefits from clustering of certain diverse functions, which may conceivably play complementary roles in the cell. We propose a simple encoding of chromosomal gene order, the neighborhood function profiles (NFP), which draws on diverse gene clustering patterns to predict gene function and phenotype. NFPs yield a 26-46% increase in predictive power over state-of-the-art approaches that propagate function across neighborhoods, thus providing hundreds of novel, high-confidence gene function inferences per genome. Furthermore, we demonstrate that the effect of structural variation on gene function distribution across chromosomes may be used to predict phenotype of individuals from their genome sequence.

## Introduction

The role of many genes remains unknown. Even in well-investigated model organisms, a quarter or more of the genes have poorly characterized function. With the advance of genome sequencing techniques, the vast amounts of accumulated data provide an opportunity to infer gene function using computational methods. While such methods occur in many varieties, one widespread approach is to examine the gene neighborhoods that occur across genomes. Then, following the principle of guilt-by-association, a function of a gene is inferred by transferring it from its neighbors in the genome^1,2^. Gene neighborhoods that are conserved across multiple genomes provide additional confidence in each inference, often yielding highly accurate predictive modes of gene function^3,4,5,6,7^. One important biological mechanism that underlies similarity of gene function in neighboring genes is that they are often regulated by common factors and therefore co-expressed (reviewed in^8,9^). The prime example is the prokaryotic operon, where a single mRNA harboring multiple protein-coding regions is transcribed from a promoter, ensuring that expression of such functionally related proteins is coordinated. However, this concept extends more broadly -- neighboring genes that are not part of the same operon are also often co-regulated and share function. Moreover, in eukaryotic organisms, which generally lack operons, gene regulation is organized regionally and this pattern can be conserved across evolutionary time^10,11,12,13,14^ Consistently, gene function is non-randomly distributed also across eukaryotic chromosomes^15,16^, even though the neighborhood signal is overall more subtle than in prokaryotes, important exceptions notwithstanding (reviewed in^17^).

The current computational methods that use conserved gene neighborhoods to predict gene function rely on the principle that similar functions cluster together on the chromosomes. While there is abundant evidence that this is the case, we were motivated by known individual examples of clustering of genes with apparently unrelated functions. For instance, a certain metabolic gene (FAD synthase) was reported to hitchhike with clusters of protein translation-related genes, and certain RNA modification/degradation genes were found to hitchhike with signal transduction genes^18,19^. We asked if these examples represent a broader trend that could be tapped into to improve methodologies that predict gene/protein function and phenotypes from genomic sequence in an automated manner. In other words, we searched for pairs of gene functions that are highly dissimilar, according to the structure of the Gene Ontology, yet that systematically cluster in genomic neighborhoods. If found, such clustering patterns would be predictive of gene function in a manner which is not accessible to previous automated approaches, which propagate a particular gene function across genomic neighborhoods. Furthermore if this type of clustering of diverse functions were widespread i.e. if it affected many genes, it would provide a basis for a broadly useful, general methodology to infer gene function that relaxes the requirement for homogeneity of function across neighborhoods.

Here, we examined the functional composition of genomic neighborhoods across 1669 prokaryotic, 49 fungal and 80 metazoan genomes (for more details see Supplementary material 1, section S1), finding that indeed it is a very common occurrence that certain pairs of unrelated functions cluster together. Almost all gene functions are significantly enriched in the genomic neighborhood of one or more semantically distant gene functions. Using this signal to infer function from neighbors results in 3.5-fold increase in predictive power (estimated by the information accretion criterion) over a naïve guilt-by-association approach, and an 1.4-fold increase over a state-of-the-art network approach. In addition to predicting gene function, accounting for the complementarity patterns in genomic neighborhoods enhances the ability to predict phenotype from structural variation in genomic sequences of individuals. Our work highlights a widespread pattern in how gene function is organized across chromosomal domains. This has implications for understanding genome evolution and brings immediate practical benefits for methods to predict gene function and phenotype.

## Results

It is known that functionally related genes reside in the same genomic neighborhood more often than expected at chance ^3,4,5,7,15,16,17^. We employed a simple framework to systematically quantify the extent of such co-occurrence in neighborhoods across many genomes, separately for individual gene functions. In brief, we used COG (clusters of orthologous groups) and NOG (non-supervised orthologous groups) gene families, to each of which we assigned a set of gene functions, herein represented by Gene Ontology (GO) terms. Of note, here we refer to ‘function’ in a general sense, encompassing all three sub-ontologies of the GO: biological process (BP), molecular function (MF) and cellular component (CC). These assignment of function to gene families were based on the known functions of the constituent genes of each COG/NOG (henceforth jointly referred to as COG) (Methods). The mapping of COG gene families to GO terms allowed us to determine whether a certain GO term *x* is enriched in the genomic neighborhood of any gene assigned to this same function *x*, normalized to the prevalence of *x* outside such neighborhoods, examined across 1669 prokaryotic genomes. This yields an odds ratio (OR_x_), which describes the effect size of the enrichment of each function GO *x* in its own neighborhood of size *k*. Given that operons tend to be short^20^, we present the data for prokaryotes with *k*=2 (we also provide systematic analyses covering larger neighborhood sizes in Supplementary document 1, Figure S27, relevant in the light of work suggesting that bacterial functional neighborhoods may extend up to 20kb^21,22^). The neighborhood is defined here as two genes to each side of a central gene, corresponding to a total size of five genes -- one central and four flanking (irrespective of orientation). We were able to examine a total of 1048 GO terms that occured in at least 5 COG gene families. Out of these GO terms, 81.5% had neighborhoods significantly enriched in that same GO term (OR_x_>1 at FDR<20%; Z-test for significance of log odds ratio; Methods) and in 78.8% was the enrichment significant and also higher than twofold (OR_x_ ≥ 2). In the usually larger eukaryotic genomes, we considered neighborhoods of *k*=5 genes to each side of a central gene, and examined the 2617 GO terms (fungi) and 2336 terms (metazoa) that occurred in at least 3 gene families (for more detailed analyses of different neighbourhood size see Supplementary document 1, Figure S27). Across eukaryotes, 35.1% of the analyzed GO terms are significantly enriched in their own neighborhoods across 49 fungal genomes, and 99.1% of GO terms across 80 metazoan genomes. 17.7% of functions are significant and at least two-fold enriched in their own neighbourhood in Fungi, and this is the case for 99.1% of the gene functions in Metazoa. These enrichments and significance calls are upheld by comparing them to enrichments computed on randomized data (see Methods and Supplementary material 1, Table S3). The above data are consistent with the known clustering of genes with similar functions across genomes^23,15,24^ and demonstrate that our approach can be used to detect function enrichment in genomic neighborhoods.

Next, we applied the same method to exhaustively test for enrichment of pairs of diverse GO functions in genomic neighborhoods. In particular, we measured the enrichment (as odds ratio, OR_xy_) of the genes annotated with a GO term *y* that are close to genes with GO term *x*, again in a neighborhood of *k*=2 genes to each side thereof. Indeed we found that 2.9×10^5^ of 1.1×10^6^ total examined pairs of GO terms in prokaryotes are significantly enriched (FDR<20%, Z-test on log OR), which makes all of the tested GO terms significantly enriched in the neighborhood of at least one non-self GO term (again, the significance calls are largely supported in a randomization test, see Methods and Supplementary material 1, Table S3). Mirroring the trend in prokaryotes, 1.0×10^6^ out of total 6.8×10^6^ considered function pairs are significantly enriched in fungi and similarly, in metazoa, 1.6×10^6^ out of 5.4×10^6^ pairs of gene functions are significant.

By design, in the GO there are many terms which describe very similar concepts and it is conceivable that a gene family might be assigned to either a GO term *x* or a very similar GO term *y* due to noise in the annotation process. This has the potential to inflate the number of non-self GO terms we observe mutually enriched in neighborhoods. We have therefore filtered out related GO term pairs by using measure of semantic similarity (Resnik similarity, RS, see Methods), which is defined using the structure of the Gene Ontology graph. Of note, because semantic similarity is only defined within each GO sub-ontology but not across sub-ontologies, we present data for the ‘biological process’ GO terms. By conservatively requiring RS<1 (prokaryotes), we effectively restrict to GO term pairs in the different branches of the ‘biological process’ GO graph, removing 60% of the pairs in our prokaryotic data (details in Methods). Even in this remaining set there exist 0.5×10^5^ out of 1.8×10^5^ significantly enriched pairs of semantically distant functions. Moreover, in prokaryotes 100% of the BP GO terms are significantly enriched (at FDR=10%) in the neighborhood of at least one other semantically distant BP GO term (Figure 1). For eukaryotes we required RS<2 (see Methods for justification), removing 87% and 79% of all GO term pairs in the BP sub-ontology in Fungi and Metazoa, respectively. Still, 0.3×10^6^ out of 1.8×10^6^ distant BP GO term pairs in Fungi are significantly enriched in genomic neighborhoods, and 0.4×10^6^ out of 1.3×10^6^ in Metazoa. These high proportions mean that almost all gene functions are significantly enriched in the neighborhood of at least one other semantically distant gene function (96.4% in Fungi, 100% Metazoa), when considering BP GO terms.

**Figure 1.**
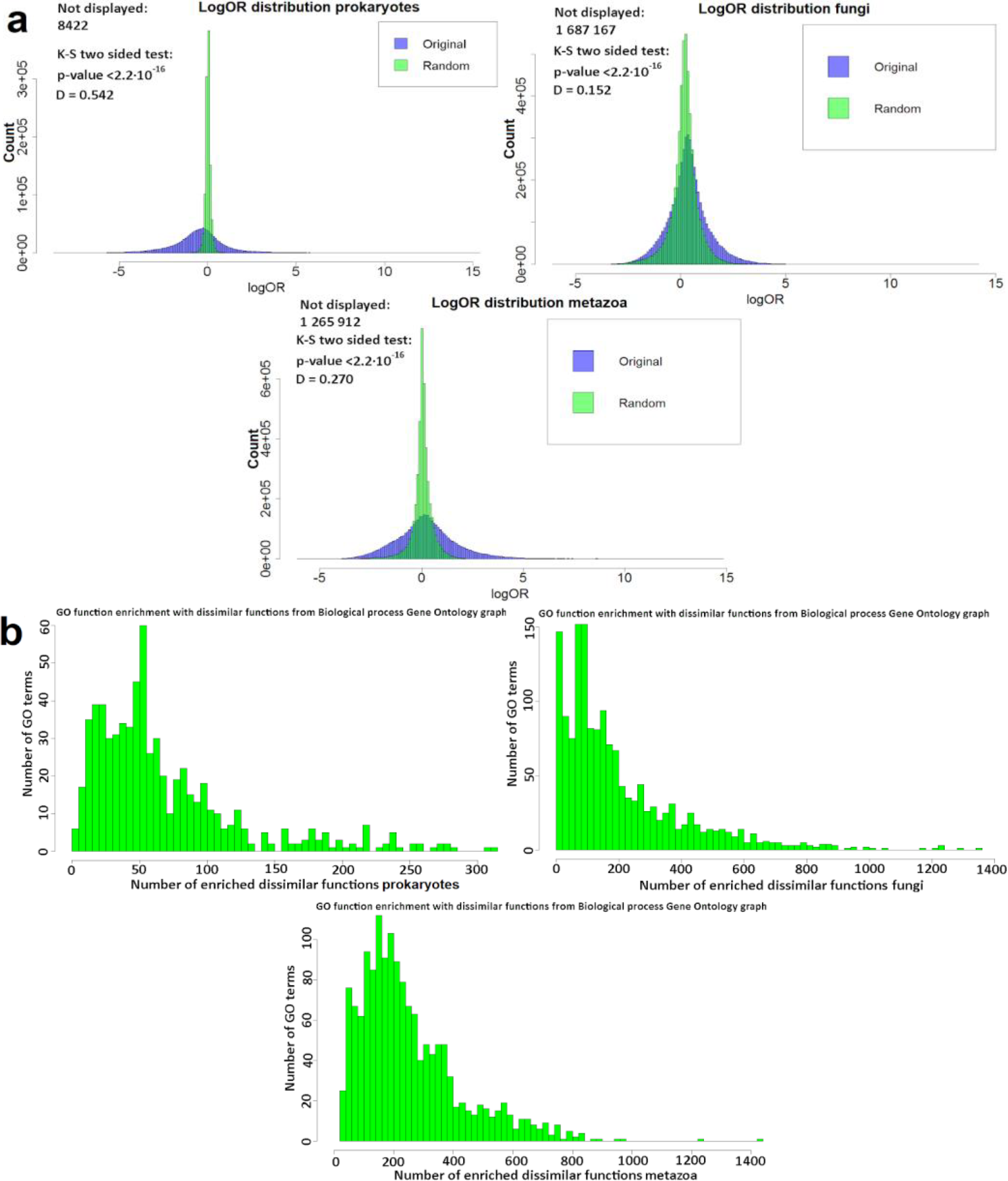
Systematic enrichment of diverse gene functions is widespread in genomic neighborhoods. (**a**) Distribution of neighborhood enrichment scores (log odds ratio, OR) for all pairs of GO functions on original and randomized genomes of prokaryotes, fungi and metazoa. See also Supplementary 1, Figures S8, S11, S12 and Supplementary 1, Table S3. Pairs with OR = 0 are not shown on graphs (see Methods; these pairs result in artifactually high or low log OR values after continuity correction). (**b**) Number of GO terms that are semantically distant, but significantly enriched in genomic neighborhoods (FDR≤10%) of each GO term, summarized in histograms for prokaryotes, fungi and metazoa. GO term pairs with Resnik similarity<1 (for prokaryotes) and RS<2 (for eukaryotes) from the ‘biological process’ GO sub-ontology are tallied in the figure.

The effect sizes of such enrichments may be substantial: 98.8%, 94% and 100% of the GO terms have a significant enrichment in the neighborhood that is also at least two-fold in magnitude with at least one other semantically distant GO term in prokaryotic, fungal and metazoan data, respectively. These statistics were also supported by comparing against a baseline distribution obtained by randomizing gene coordinates (Supplementary material 1, Table S3). GO term enrichments in neighborhoods of four example gene functions are shown in Figure 2, illustrating how there exist pairs of dissimilar functions that have neighborhood enrichments comparable to or higher than the enrichment of a gene function in its own neighborhood (histograms including statistical significance calls are shown in Supplementary 1, Figure S7).

**Figure 2.**
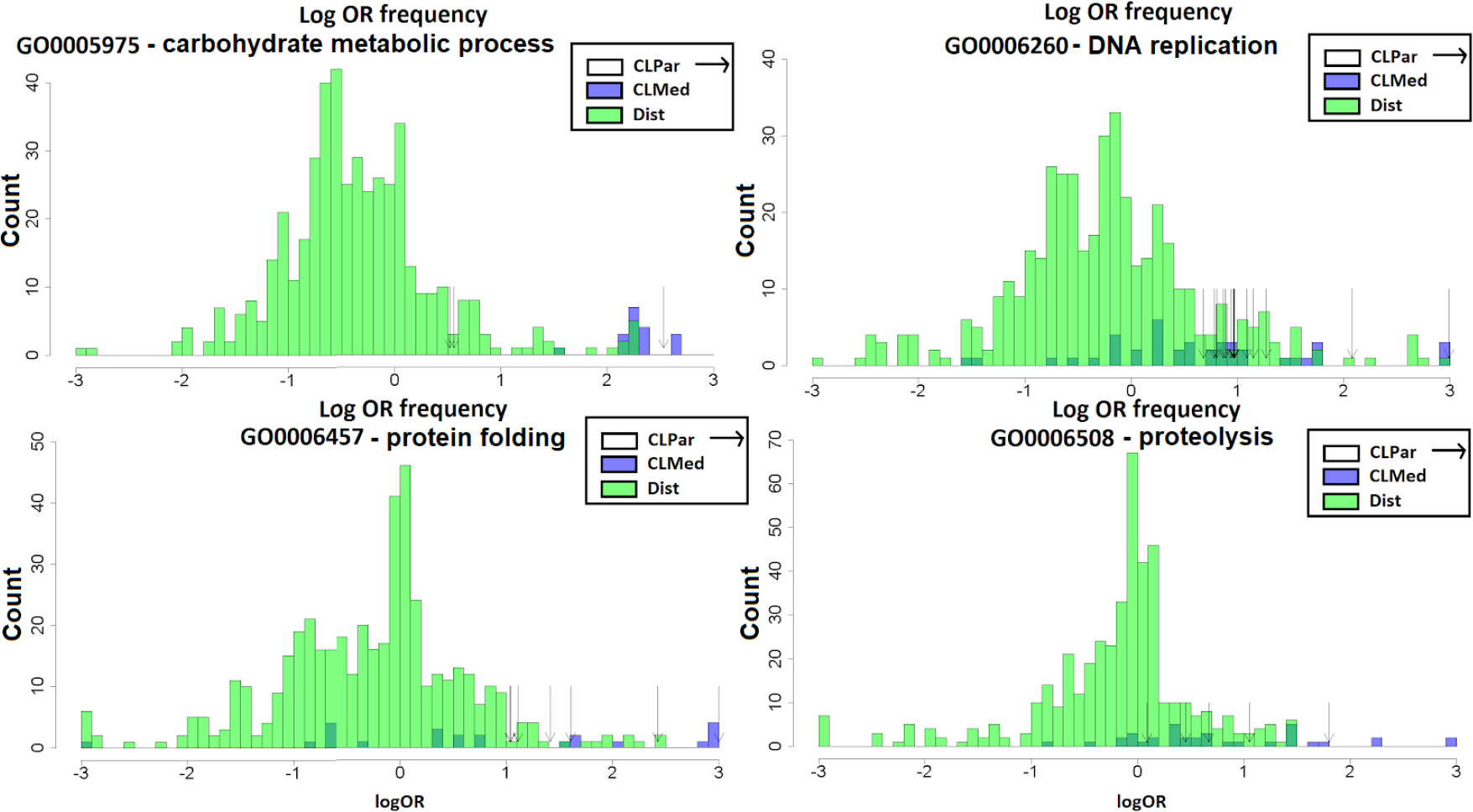
Semantically distant GO terms can be as strongly enriched in gene neighborhoods as the semantically close GO terms. For four example GO terms of the ‘Biological process’ ontology, histograms show numbers of GO terms at a certain log odds ratio (log OR) of the enrichment in gene neighborhood (for prokaryotic genomes). The GO terms in neighborhoods of a central GO function are broken down into three groups: the “CLPar” group (the central function itself + all its parent functions in the GO graph), “CLMed” group (functions with Resnik semantic similarity > 2 with the central function) and “Dist” group (Resnik≤2 with the central function). Instances of GO terms in the dissimilar “Dist” group and in the non-self “CLMed” group can be observed that have enrichments as high or higher as the self-enrichments (the “CLPar” functions).

For instance, there is a global trend in gene neighborhood organization where the GO term “*carbohydrate metabolism*” is 2.12-fold enriched in the neighborhood of the semantically unrelated GO term “*carbohydrate transport*” (of note, the magnitude of this enrichment is similar to that of “*carbohydrate metabolism*” in its own neighborhood – 2.53-fold). This co-occurrence is reminiscent of the textbook example of the *lac* operon in *Escherichia coli*, where the *lacY* gene encodes lactose permease, which shuttles lactose into the cell, while the neighboring *lacZ* gene encodes the enzyme β-galactosidase, which breaks down lactose into monosaccharides. Our analysis suggests that the proximity of genes encoding carbohydrate transporters and carbohydrate metabolizing enzymes is a systematic trend. This can provide supporting evidence for inferring the presence of putative transporters next to known metabolic enzymes, and also *vice versa*.

Some other examples include significant enrichment of both “*DNA topological change*” (OR_xy_=1.42) and also of “*nucleoside bisphosphate biosynthesis*” (OR_xy_=1.41) in the neighborhood of gene families annotated with ‘‘*response to DNA damage stimulus*”, which suggests coordinated regulation between DNA repair-related activities. Furthermore, “*lipid biosynthesis*” genes are often in the neighborhood of “*localization within membrane*” genes (2.43) and “*carbohydrate biosynthesis*” is in the neighborhood of “*cell envelope organization*” (2.10). These associations suggest coordinated activation of processes that generate building blocks of cellular structures, and subsequent processes that incorporate the building blocks into these structures. We list these and other examples of enriched GO term pairs for prokaryotic neighborhoods in Table 1, while the exhaustive list is in Supplementary Table S5.

**Table 1.**
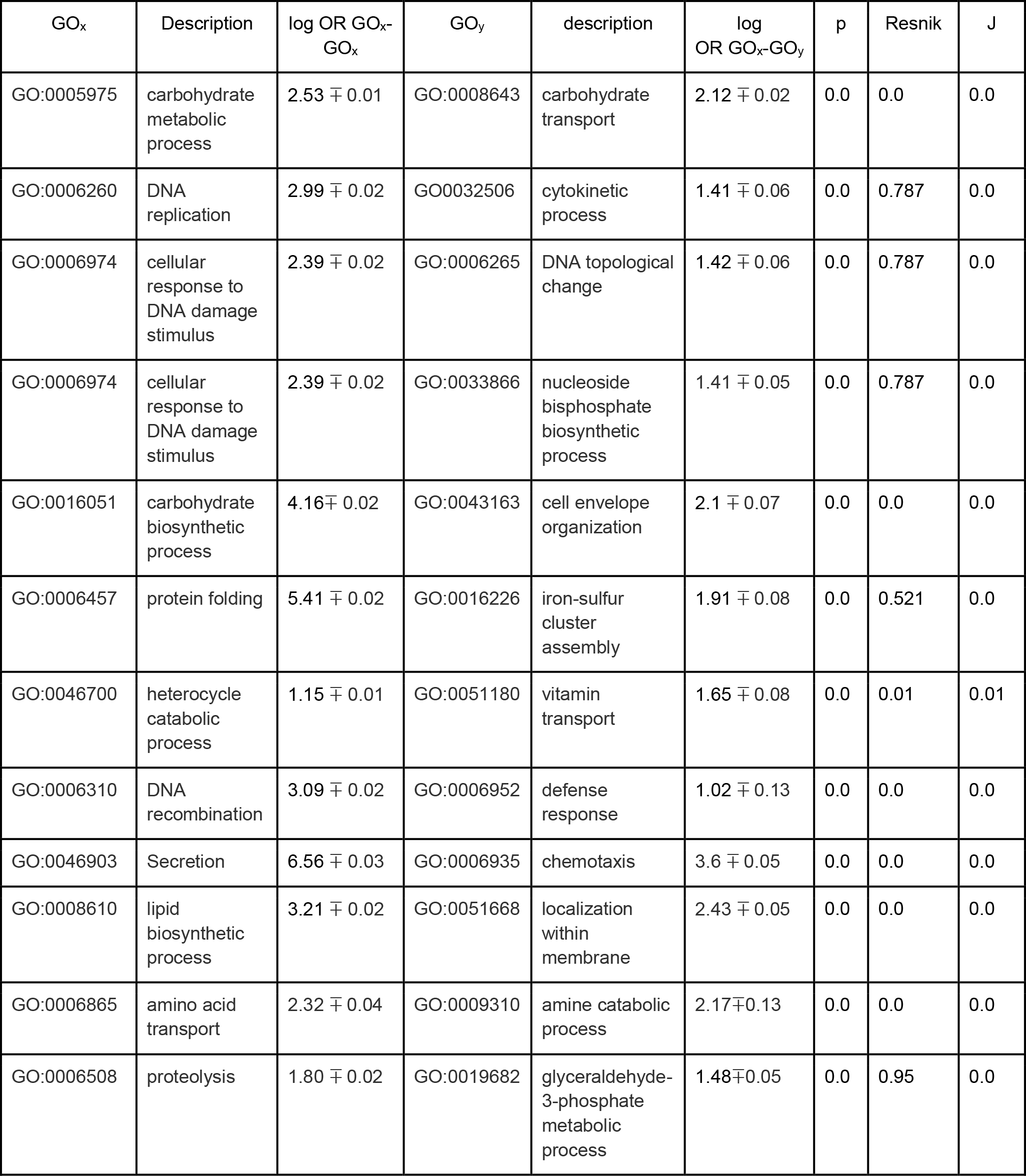
Examples of dissimilar gene function pairs enriched in genomic neighborhoods. Data shown for “biological process” GO graph of the prokaryotic genomes. GO_x_-GO_x_ denotes enrichment of a GO term *s* in its own neighborhood, while GO_x_-GO_y_ denotes enrichment of the other GO term *y* in the neighborhood of GO term *x*. Enrichments are given as log odds ratio (log OR) +/- corresponding 95% confidence interval and the p-value for the association (by Z-test on log OR, one-tailed). “Resnik” denotes Resnik semantic similarity; this is an unbounded score where any value <2 corresponds to very distant terms that reside in separate branches of the GO graph (meaning the closest common ancestor has information content <2). “J” is the Jaccard index that quantifies co-occurrence of gene functions in COG gene families from our prokaryotic data set, and can vary from 0 to 1.

While all GO term pairs listed above either have low semantic similarity or are from different GO sub-ontologies, it is possible that GO terms which are distant in the GO graph may overlap in the set of genes assigned to them^24^. This has the potential to inflate the observed enrichments, which might be due to the same genes artifactually creating two apparently distinct functional neighborhoods. However after quantifying the overlap of genes assigned to the pairs of GO terms using Jaccard index, we find this not to be a common issue (Table 1; Supplementary Material 5). Furthermore, a randomization of gene positions within each genome supports that the observed enrichment of extreme OR values between distinct GO terms is nonrandom. Importantly, this holds true to a similar extent for the semantically very close GO term pairs and the semantically distant pairs (panels ‘CLPar’ and ‘Dist’ in Figure S8). Overall, our data suggests that clustering of semantically distant gene functions is very common in genomic neighborhoods, occurring to a similar extent as the well-known clustering of similar gene functions in genomes.

### A general method to infer gene function based on neighborhood patterns

Having demonstrated that genomic neighborhoods are significantly enriched in certain combinations of diverse gene functions, we asked if such neighborhood patterns can be used to derive a general method that predicts gene function. To evaluate this, we compared two established methods that propagate a gene function to neighboring genes, with a novel classifier that can draw on neighborhood co-occurrence of diverse gene functions to predict GO terms for COG gene families. The first method is a simple k-nearest neighbors (kNN) classifier that transfers known functions to a COG gene family from the *k* neighboring gene families with smallest average logarithmized distances across many genomes (Methods). The second classifier can transfer gene functions to neighbors additionally via indirect links: a gene network is constructed from neighborhoods and the gene function assignments diffuse across the links (using the GeneMania method^25^, here referred to as the Gaussian Field Label Propagation (GFP) classifier). In addition to these known approaches, the third, novel classifier can draw on both the enrichment of similar functions in neighborhoods and additionally the enrichment of semantically distant gene functions. For example, this method should be able to infer that a gene family deals with “*carbohydrate metabolism*” based on its neighbors being annotated with “*carbohydrate metabolism*” and additionally based on its neighbors dealing with “*carbohydrate transport*”, a semantically distant function in the Gene Ontology graph. To this end, we employed a Random Forest classifier on a data set constructed such that examples are COG gene families, features are the normalized counts of every GO term in the neighborhood of that COG (across all genomes), and the target variable is the set of known GO term labels of the COG gene family; see Methods for a more formal definition. In other words, from such a representation, the function of a gene can in principle be inferred from the presence of any function -- or a combination thereof -- in the genomic neighborhood of the gene, as long as such a pattern occurs commonly enough to be recognized by the algorithm. We named this approach the “neighborhood function profile” (NFP) classifier.

We first evaluated the accuracy of all methods in a cross-validation test (using the out-of-bag error statistic provided by Random Forest; see Methods) using the 3475 prokaryotic COG gene families and 1048 corresponding GO terms, 15741 fungal COGs and 2617 corresponding GO terms, and 9185 metazoan COGs and 2336 corresponding GO terms. The average area under precision-recall curves (AUPRC) for GO terms in the prokaryotic dataset is 0.153 for the 10-NN classifier and, expectedly, a much improved 0.199 score for the network-based (GFP) classifier. Both methods operate by transferring an annotation across gene neighborhoods, while the latter also has the ability of using indirect links to improve accuracy. However the NFP-based classifier which can draw on diverse neighborhood patterns substantially improves this with a 0.266 average area under P-R curves in prokaryotes, a 34% increase (*p* < 10^−10^ for the increase over the next best method, one-tailed Wilcoxon test on AUPRC scores across GO categories; distributions of scores in Figure 3, see also Supplementary 1, Figures S16, S17 and S18). Similarly so, in the two groups of eukaryotic organisms, the AUPRC scores of the novel NFP method were significantly improved: for Fungi, the 0.0460 (for the 10-NN) increased to 0.0545 (for the NFP; *p* < 10^−15^ for the increase, Wilcoxon test) and for Metazoa, the increase is 0.0228 (for 10-NN) to 0.0267 (for NFP, *p* < 10^−15^ for the increase; Figure 3). The network diffusion approach applied to gene neighborhood data in eukaryotes did not on average bring benefits over the simpler 10-NN (Supplementary 1, Figures S19–S24), therefore the latter is used as a baseline.

**Figure 3.**
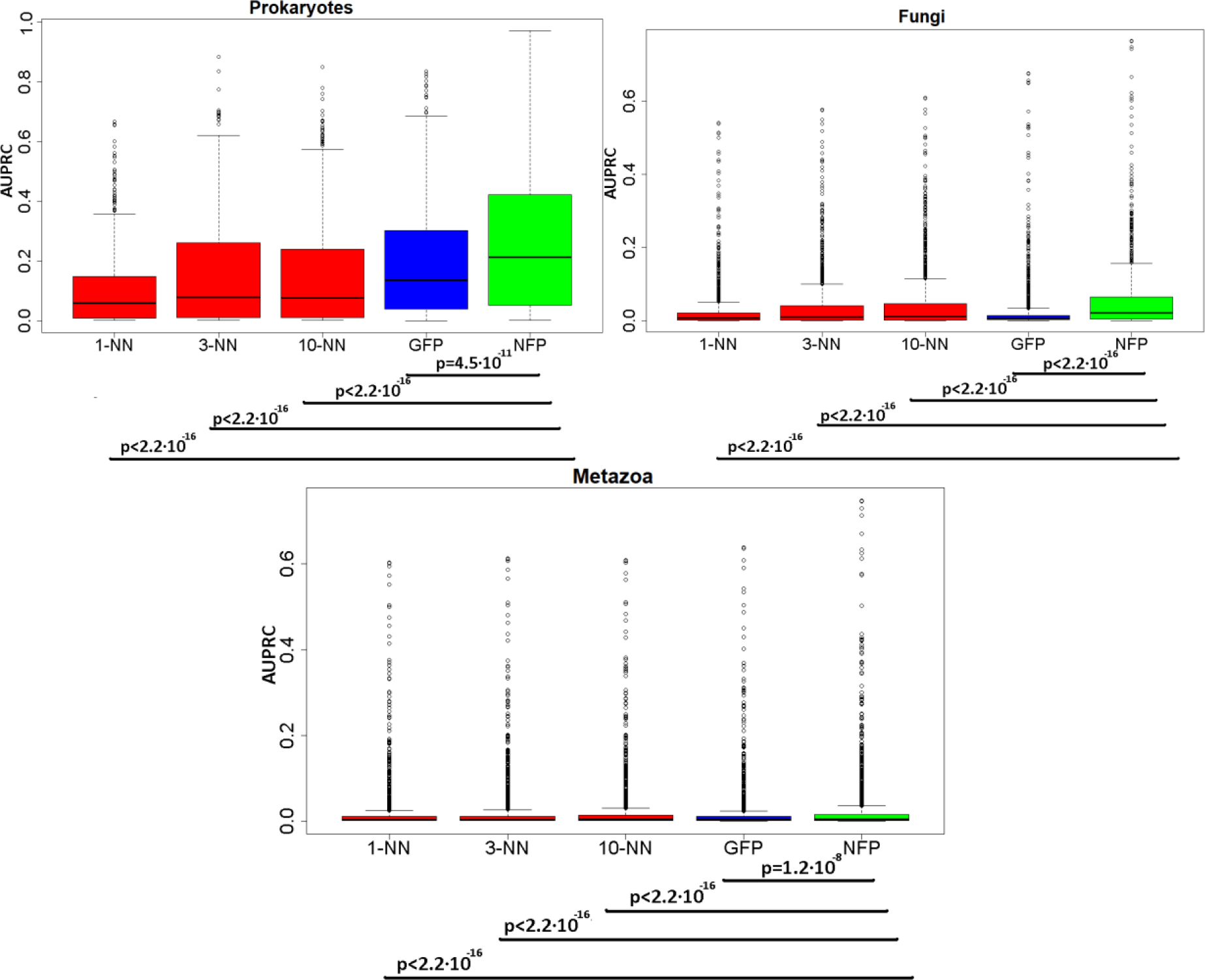
The gene function profile of genomic neighborhoods provides a more accurate method to infer gene function. The distribution of area under the precision-recall curve (AUPRC) scores, measured in cross-validation, across all examined gene functions in prokaryotes (top), fungi (middle) and metazoa (bottom). The methods compared are the nearest neighbor (NN) classifiers (1-NN, 3-NN, 10-NN), a network-based approach (Gaussian Field Label Propagation, GFP) and finally the novel Neighbourhood Function Profile (NFP) method. See Supplementary 1, Figures S17, S20 and S23 for the area under ROC curve (AUC) scores. P-values are from a one-tailed Wilcoxon signed-rank test.

### Predictive power of the Neighborhood Function Profile (NFP) classifier

A more accurate classifier would be expected to provide a higher number of predictions at a certain confidence threshold. We quantified this increase provided by the predictive models based on NFP. In particular, we tallied the number of predictions (COG-GO term associations) made at a precision threshold of 50% (equivalent to 50% FDR) and additionally at a more stringent 80% (20% FDR) for the three classifiers; Table 2. This reveals remarkable, several-fold increases in the amount of predictions afforded by the NFP in prokaryotes, fungi and metazoa, when considering the more general gene functions (information content (IC) between 2 and 4). In the highly specific gene functions (IC>4), there is a substantial increase in new annotations provided by NFP for prokaryotes and fungi. In metazoa however, the gain afforded by NFP in the IC>4 group of functions is more modest. We further examined the diversity of predictions: in particular, we asked if the new predictions afforded by NFP are largely added to the gene families that were already assigned predictions by the known methods, or they instead afford coverage of new gene families. Our data suggested that the latter is the case (Supplementary 1, Tables S5, S6 and S7), because the number of COG gene families receiving at least one prediction increases in NFP compared to the baseline classifiers.

**Table 2.** Number of predictions (associations between a gene function and a COG gene family) obtained using different gene function prediction methods based on neighborhoods. 10-NN, ten nearest neighbors; GFP, Gaussian Field Label Propagation (network-based approach); NFP, neighborhood function profile. IC, information content of GO term, where lower IC signifies more general functions. Bold numbers show the best method for a given combination of dataset, stringency and set of functions. The exhaustive list of annotations obtained by the NFP can be seen in Supplementary material 2 and novel predictions in Supplementary material 3.

| Dataset | Method | Number of predictions<br>(functions of any IC) | | Number of predictions of<br>general functions with<br>$2 < IC \leq 4$ | | Number of predictions of<br>specific functions with<br>$IC > 4$ | |
| --- | --- | --- | --- | --- | --- | --- | --- |
|  |  | Precision<br>0.5 | Precision<br>0.8 | Precision<br>0.5 | Precision<br>0.8 | Precision<br>0.5 | Precision<br>0.8 |
| Prokaryotes | 10-NN | 31,759 | 4,828 | 3,642 | 1,093 | 5,664 | 1,942 |
|  | GFP | 61,418 | 15,094 | 11,740 | 3,089 | 13,194 | 4,763 |
|  | NFP | **88,579** | **25,635** | **26,448** | **7,247** | **16,804** | **6,228** |
| Fungi | 10-NN | 65,370 | 448 | 17 | 0 | 1,284 | 448 |
|  | GFP | 140,255 | 20,020 | 3 | 2 | 0 | 0 |
|  | NFP | **178,687** | **20,204** | **3,592** | **355** | **7,668** | **1,579** |
| Metazoa | 10-NN | 66,403 | 327 | 912 | 74 | 936 | **253** |
|  | GFP | 89,992 | 464 | 2,552 | 11 | 390 | 94 |
|  | NFP | **102,631** | **3,057** | **9,546** | **769** | **988** | 217 |

Our gene neighborhood classifiers, as implemented, provide function predictions at the level of COG gene families (exhaustive list given in Supplementary material 2 and 3.). We also provide data showing how this reflects in the number of genes receiving predictions in certain model organisms (Supplementary 1, Tables S7 and S8). For instance, in *Escherichia coli*, at precision 0.5, the number of novel predictions (gene-function pairs) is 7572 for the network approach, and increases to 10559 provided by the novel NFP classifier; similarly so in the pathogen *Staphylococcus aureus*, increasing from 4314 to 5386 by use of NFP; all counts given for gene functions with IC>4. Eukaryotes, consistent with more modest AUPRC scores (see above), provide overall fewer predictions but increases by NFP are still quite evident. The number of novel annotations at Pr=0.5 substantially increased from 89 (10-NN) to 552 (NFP) for *Saccharomyces cerevisiae* and 87 to 282 for *Schizosaccharomyces pombe*, in the fungal predictor (Supplementary 1, Table S8). For metazoans, the increases were striking for the more general gene functions with IC 2-4, with more than twofold higher number of predictions afforded by NFP at precision=0.5 for mouse, human or *Drosophila melanogaster*, compared to the next best method based on propagating functions across the neighborhoods. The NFP gains were more modest for highly specific gene functions with IC>4 in Metazoa (Supplementary 1, Table S8). One possible explanation was the lower coverage with known functions (average 18 GO terms per gene in Metazoa *versus* 26 in Fungi in the databases we used; see Methods). This might prevent NFP from discovering complex association patterns between gene functions in neighborhoods, while the simpler kNN classifier is less affected.

We examined another measure of the utility of the predictive models, based on the information accretion (IA) criterion^26^. In brief, IA weighs the predictions such as to give higher scores to higher information content (rarer) GO terms; see Methods. By this method, in prokaryotes kNN predicts 2.69 bits/gene novel information, the GFP network approach 6.89 bits/gene, while the NFP increases this to 9.44 bits/gene; all given at precision=0.5 (bits/gene at precision 0.5 and 0.8 thresholds are in Table 3, for visualization of proportions see Supplementary 1, Figures S30 and S31). Therefore, we suggest that NFP brings a 37% increase in coverage with predicted gene functions over a state-of-the-art genomic neighborhood method. This result is mirrored in two groups of eukaryotes we tested: in fungi, the increase was 0.70 (kNN), 1.37 (network) and 2.00 (NFP) bits/gene, while in Metazoa it was 0.70 (kNN), 0.99 (network) increasing to 1.25 bits/gene in the NFP method (Table 3, Supplementary 1, Figures S32, S33, S34 and S35). In other words, in eukaryotes, the new NFP method increases the predictive power by 46% (Fungi) and 26% (Metazoa) over a state-of-the art network approach for propagating gene function across neighborhoods.

**Table 3.** Amount of predicted information on gene function, measured using the ‘information accretion’ methodology and expressed as bits per gene. 10-NN, ten nearest neighbors; GFP, Gaussian Field Label Propagation (network-based approach); NFP, neighborhood function profile.

| Dataset | Stringency (Precision score threshold) | Method | Known annotations (bits/gene) | Recovered known annotations (bits/gene) | Newly predicted annotations (bits/gene) |
| --- | --- | --- | --- | --- | --- |
| Prokaryotes | 0.5 | 10-NN | 23.19 | 4.31 | 2.69 |
|  |  | GFP |  | 7.71 | 6.89 |
|  |  | NFP |  | **10.22** | **9.44** |
|  | 0.8 | 10-NN |  | 1.16 | 0.15 |
|  |  | GFP |  | 3.09 | 0.77 |
|  |  | NFP |  | **4.78** | **1.14** |
| Fungi | 0.5 | 10-NN | 27.91 | 1.08 | 0.7 |
|  |  | GFP |  | 1.96 | 1.37 |
|  |  | NFP |  | **2.59** | **2** |
|  | 0.8 | 10-NN |  | 0.01 | 0.001 |
|  |  | GFP |  | 0.39 | 0.087 |
|  |  | NFP |  | **0.45** | **0.093** |
| Metazoa | 0.5 | 10-NN | 19.05 | 1.01 | 0.7 |
|  |  | GFP |  | 1.34 | 0.99 |
|  |  | NFP |  | **1.56** | **1.25** |
|  | 0.8 | 10-NN |  | 0.01 | 0.002 |
|  |  | GFP |  | 0.03 | 0.003 |
|  |  | NFP |  | **0.08** | **0.014** |

### NFP accuracy is due to semantically distant functions

The above data indicate that a classifier based on the NFP -- an exhaustive description of the composition of gene functions in a genomic neighborhood -- provides high accuracy and yields many additional function predictions. Next, we performed tests to ascertain if this increase in accuracy of the NFP is indeed specifically due to the semantically distant GO terms enriched in gene neighborhoods. To this end, we examined 11 example gene functions, which were shown to have other semantically dissimilar functions enriched in their neighborhoods (by our OR_xy_ measure, see above). As expected, the NFP classifier strongly outperforms the baseline kNN and the GFP classifiers for these functions (Supplementary 1, Figure S28). Then, we created partial NFPs, which contained only the semantically distant functions, compared to the function being predicted (“Dist”, having RS < 2), contrasting this to the partial NFP which contains only the function being predicted itself and its ancestors that are semantically close (“CLPar”, parents in the GO graph having *RS* ≥ 4 plus the GO function itself). Random Forest classifiers were trained on the partial NFPs and cross-validation accuracy compared (Figure 4). For 9 out of 11 functions, a higher accuracy was obtained from neighborhood features describing semantically distant functions (“Dist/Par”, mean AUPRC = 0.047 across gene functions) than from features describing of the target function and its close parents (“CLPar”, mean AUPRC = 0.035), which is an implementation of the guilt-by-association principle using the same classifier and the same data representation; for 8 out of 11 the increase was significant at p<0.0002; Wilcoxon one-sided paired test. For all 11 functions, using the full set of neighborhood features (including the close, intermediate-distance, and distant functions), significantly outperforms the default model using only close features, suggesting a benefit to predictive accuracy also from including the intermediate-distance functions. This analysis demonstrates that even when using the same statistical methodology (Random Forest) and same type of data representation (NFP), the presence of semantically distant functions in genomic neighborhoods is often highly predictive of a gene having a certain function. For more details on the experimental setup see Supplementary document 1 (methodology section).

**Figure 4.**
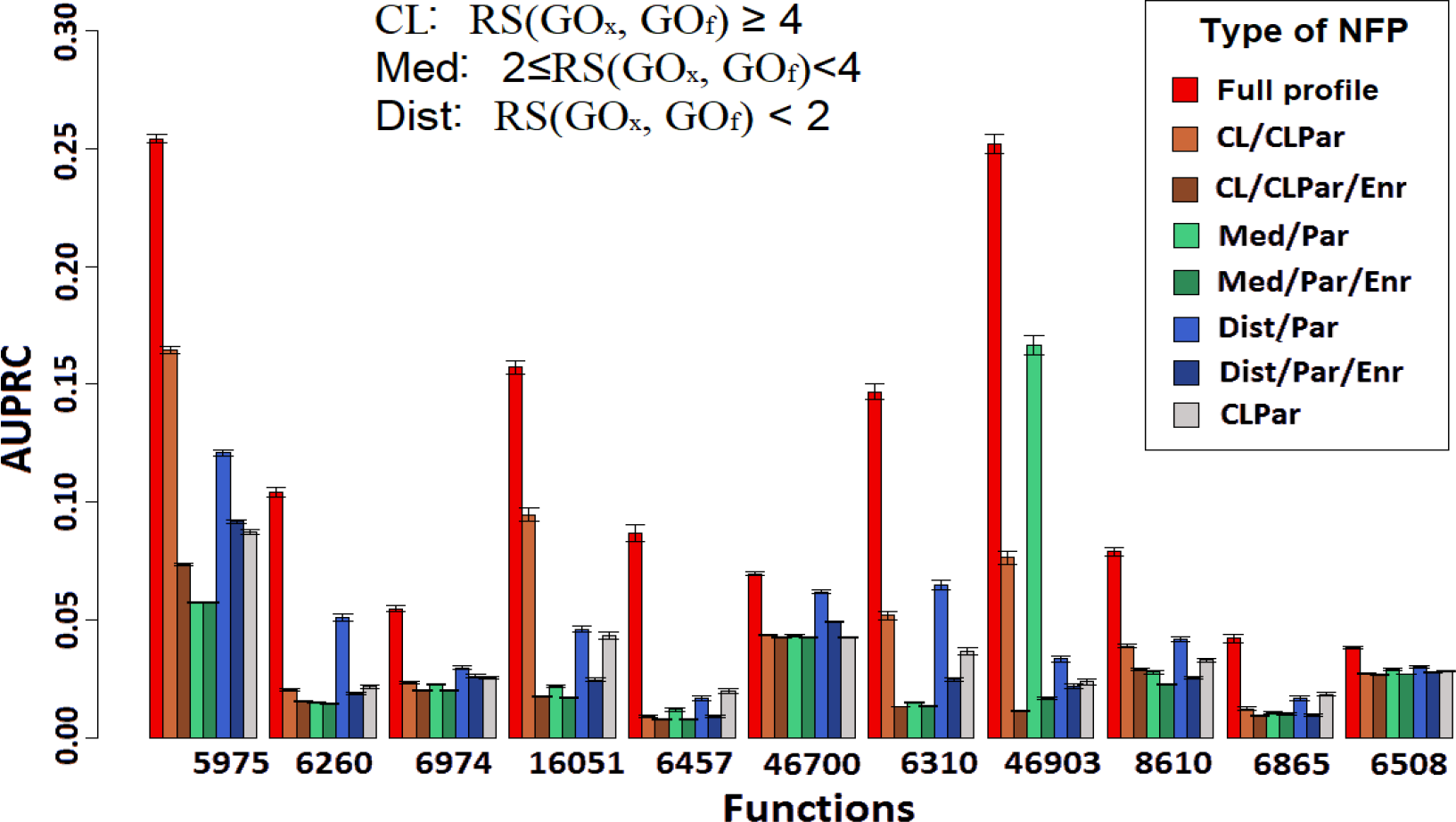
Semantically distant functions in gene neighborhoods are important for accurate inference of gene function. Bars show accuracy (as AUPRC score, measured in crossvalidation) for predicting the eleven representative gene functions, using various types of neighborhood function profiles (NFP) that are listed in the legend. The “Full profile” are the full NFP of the ‘biological process’ GO graph, while the “CL/CLPar”, “Med/Par” and “Dist/Par” represent the partial NFP consisting only of close, medium-distance and distant functions, respectively (the “/Par” denotes that parent GO terms of the target functions were removed). The “CLPar” partial profiles contain only the selected function and its semantically close parents, meaning that “CLPar” is an implementation of the standard approaches that transfer similar functions across neighborhoods. In many cases, the close (but non-self), medium-distance and distant functions are more predictive than CLPar, and the complete profile (BPP) is most predictive. Serving as a control, the removal of the significantly enriched functions (labeled as “/Enr” in the legend) from the partial NFP strongly reduces accuracy, either for the close functions (CL), the medium-distance (Med) or the distant functions (Dist). Bars are average AUPRC scores of 200 runs of cross-validation of the Random Forest classifier, whereas error bars show standard deviation across the 200 runs.

Generalizing this principle, we further included GO terms from all three sub-ontologies (Biological Process [BP], Molecular Function [MF] and Cellular Component [CC]) into global functional profiles of gene neighborhoods, which therefore includes a variety of semantically unrelated GO terms. While testing this, we took provisions to exclude the pairs of GO terms across different ontologies that are commonly mapped to same genes, using a very stringent threshold of Jaccard similarity <0.1. Including all sub-ontologies further helped improve accuracy of predicting GO terms (for details see Supplementary Document 1, Section S2.10 and Supplementary 1, Figure S38). Even thus, we found that the average AUPRC score increased from 0.296 to 0.298 for BP (when including non-overlapping MF and CC terms that occur in neighborhoods), 0.191 to 0.213 for MF (when including BP and CC) and 0.400 to 0.461 for CC (including BP and MF). This further supporting the general notion that a variety of unrelated gene functions tend to be organized into common genomic neighborhoods.

### Validation on external datasets

Our gene neighborhood-based NFP predictive models used cross-validation to obtain overall estimates of accuracy, and additionally to estimate the FDRs for thousands of individual predictions made in bacteria and eukaryotes (Supplementary material 2 and 3). To further support these estimates, we analyzed an external dataset of gene functions derived from the Critical Assessment of Function Prediction (CAFA 2) challenge data^27^ (https://biofunctionprediction.org/cafa/) (Supplementary 1, Section S2.9). Indeed, also on the external validation set, the NFP approach (average AUPRC: 0.207) again outperforms the 10-NN method (AUPRC: 0.130); the difference is significant at p<2.2×10^−16^ by Wilcoxon test, one-tailed). The average AUPRC of 0.207 on the external validation set is broadly consistent with the average AUPRC of 0.266 on crossvalidation. The AUPRC scores for the individual GO categories were significantly correlated between crossvalidation and the external set (Supplementary 1, Figure S36).

External validation of eukaryotic predictive models is consistent with the above. The accuracy on external data is higher for NFP (average AUPRC: 0.065 in Fungi, 0.025 in Metazoa), compared to the (next best) 10-NN approach (average AUPRC: 0.055 in Fungi, 0.0195 in Metazoa); the differences are significant at p<5.3×10^−16^ in Fungi and Metazoa. These scores for the NFP predictions on the external set are largely similar to those originally obtained in crossvalidation for the two groups (external 0.065 and 0.025 for Fungi and Metazoa, *versus* crossvalidation 0.0545 and 0.0267 respectively; Supplementary 1, Figure S37).

In summary, external validation confirmed that NFP outperforms previous approaches that propagate gene functions across neighborhoods, and additionally lent credibility to the original cross-validation estimates of accuracy. The set of predictions we supply as Supplementary material 2 and 3 may be used to prioritize further validation work. The FDR score provided for each individual prediction allows the users to make informed decisions on choosing the predictions to validate in further experiments.

### Gene neighborhood composition predicts phenotype of individuals

Predicting phenotype from the genome sequence of an individual is a central goal in modern genetics. While single-nucleotide variants and indels are commonly considered in such analyses ^28,29^, structural variants also have considerable potential to affect gene regulation and may therefore bear on the phenotype. Encouraged by the high accuracy of the NFP classifiers in predicting gene function based on neighborhood composition, we therefore asked if a related method could be used to infer phenotype from gene order observed in genomes of individuals in a population. We focused on prokaryotes, for two reasons. First, the NFP classifiers were more accurate for prokaryotes, which is likely at least in part due to a much larger set of genomes currently available. Second, structural variants are known to be abundant between microbial strains and affect the major part of the genome therein. Moreover, our recent work has shown that across prokaryotic species, many phenotypes are strongly associated with certain gene neighborhoods^30^. This motivated us to examine to what extent this holds true also for individuals (strains) of one species and to what extent are the associations with neighborhoods predictive. We have therefore examined a previous data set of 696 naturally-occurring *E. coli* strains that have been systematically tested for 151 phenotypes (Methods), such as ability to metabolize certain substrates or resistance to a variety of toxins and antibiotics^29^. In the original work, occurrence of deleterious variants, such as nonsense variants or frameshifting indels in certain genes was associated with specific phenotypic outcomes.

Here, as a baseline we use conditional scores (CS) supplied in Galardini *et al.*, which are based on an estimate of gene disruption in a particular strain combined with the phenotypes that are known to result from loss-of-function mutations for each gene^29^. Upon computing the AUC and AUPRC predictive performance measures for each phenotypic trait (here encoded as a binary outcome; see Methods) based on the CS, we obtained the median AUC of 0.672 (0.591 - 0.736; Q1-Q3) across the 151 phenotypes (Figure 5a).

**Figure 5.**
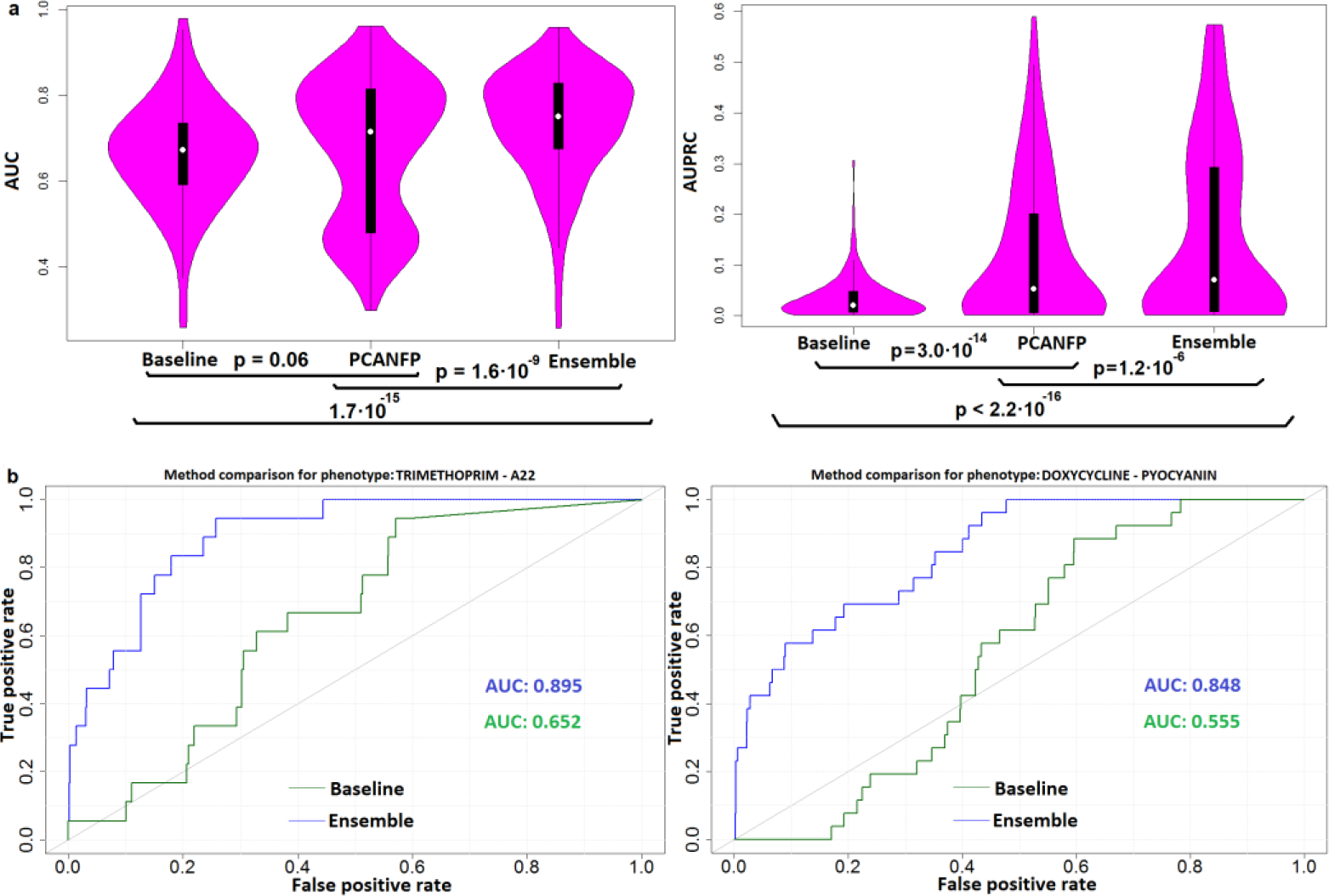
Predicting phenotypes of individuals from the effects of structural variants on the composition of gene neighborhoods. (**a**) Accuracy of predictive models’ AUC scores (top) and AUPRC scores (bottom) across 151 *Escherichia coli* phenotypes, estimated in crossvalidation. The baseline classifier predicts phenotype from the scores based on gene disruption by small variants. The PCA-NFP classifier predicts from neighborhood function profiles, which are a representation of structural variants in the genomes. The Ensemble classifier is a combination of both sources of data (see Methods). **(b)** The cross-validation receiver operating characteristic (ROC) curve achieved by a baseline method based on small genetic variants (green) and the ensemble method (blue) that also includes structural variants, shown for two example phenotypes. Additional examples are in Supplementary 1, Figure S43.

Next, we created a NFP dataset from this genomic data, where examples are *E. coli* strains, while features are frequency counts of each GO term in the neighbourhood of each COG. A principal components (PC) analysis was applied to reduce this data set to 228 PCs that provide a compact representation of the functional composition of gene neighborhoods across many gene families and which were used to train a Random Forest classifier. This yielded NFP models with broadly improved predictive accuracy, resulting in out-of-bag AUC scores of 0.7150 (0.4800 - 0.8145; median, Q1-Q3) across the phenotypes. In specific, 42 out of 151 phenotypes had a significant increase in accuracy (FDR=20%, DeLong test) over the baseline classifier that draws on deleterious point mutations / indels. In contrast, only 9 of 151 phenotypes had reduced accuracy in the NFP. Overall this suggests that composition of gene neighborhoods is substantially associated with phenotype.

The baseline classifier consults deleterious mutations but not structural variants (which manifest in changed gene neighborhoods). Furthermore, it is expected that, broadly, the effects of the point mutations/indels on the one hand and the structural variants on the other hand will often be qualitatively different -- the former commonly abolish or modify protein function, while the latter commonly affect gene regulation. We therefore hypothesized that the two types of variants need to be considered jointly to predict phenotype of individuals more accurately. This was tested by constructing an ensemble classifier (see Supplementary Document 1, Section S2.11, and Supplementary 1, Figures S41 and S42 for details) that results in further significant increases (AUC 0.761; Q1-Q3: 0.673 - 0.848) over the baseline and also over the NFP classifier (*p*_baseline_=1.7 ⋅ 10^−15^, *p*_NFP_ = 1.6 ⋅ 10^−9^, one-sided Wilcoxon signed-rank test). There was a significant increase in accuracy (FDR<20%, DeLong test) over the baseline model that draws only on deleterious gene variants, when predicting 62 out of 151 phenotypes, while only 10 phenotypes showed a significant decrease in accuracy over the baseline, signalling an increase in predictive ability for many different phenotypes.

We highlight some examples. The phenotype “Trimethoprim.A22”, which describes growth inhibition by a combination of two antibiotics, can be predicted by the baseline method (drawing on deleterious mutations) such that, at a precision=0.5, only 6% of the strains exhibiting the growth phenotype are able to be recovered by the model (recall=0.06; estimates from precision-recall curves in crossvalidation; Figure 5b). In contrast, the ensemble method which combines the mutations and the structural variants, can recover 28% of the strains exhibiting the growth phenotype (recall=0.28), which is a four-fold increase. Furthermore, the phenotype “Doxycycline. Pyocyanin” which describes sensitivity to a combined treatment by an antibiotic and a reactive oxygen species-generating toxin, does not yield any predictions (recall=0) at a precision threshold of 50% when drawing on mutations only, but recovers 27% of the strains known to exhibit growth phenotypes (Figure 5b) when considering also the gene neighborhoods encoded via their function profile. This data for other phenotypes is listed in Supplementary Material 4. This demonstrates that structural variation in the genome of individuals can be used to predict many phenotypes by drawing on the NFP representation of gene ordering along the chromosomes.

## Discussion

Our work characterizes the distribution of gene function across genomic neighborhoods in hundreds of genomes. Expectedly, we detected the well-known phenomenon where genes with similar function cluster together in eukaryotic and prokaryotic genomes. However, the same analysis revealed that another type of genomic pattern is very common -- the clustering of certain pairs of gene functions that appear unrelated, measured either by their proximity in the Gene Ontology graph (via the Resnik similarity) or the overlap in genes assigned to the functions (via the Jaccard coefficient). The prevalence of such clustering is very high: while 92.6% of all examined gene functions in prokaryotes are significantly enriched in their own neighborhood or in the neighborhood of a related function, 100% of all functions are so in a neighborhood of at least one unrelated function (at a stringent threshold of FDR=1%; see Methods for definitions). Similarly so, also in the eukaryotic groups we have examined, a higher number of gene functions have an unrelated function significantly enriched in their neighborhood than have a related function enriched (Methods for details). In other words, while neighborhood organization following the similarity principle is certainly prevalent in genome neighborhoods, other patterns appear similarly or more widespread – they are the rule rather than the exception. Given that the effect sizes of these between-function neighborhood enrichments often approaches the within-function enrichment, it is conceivable that this widespread pattern is too a result of selective forces shaping genome organization, although the underlying evolutionary mechanisms remain to be elucidated. Irrespective of the mechanism that created it, this pattern is sufficiently strong that it can yield accurate predictive models to infer gene function and also the phenotype of individuals.

If indeed the clustering between unrelated gene functions brings a selective benefit to the organism, one possible interpretation is that the clustered pairs of functions play complementary roles in the physiology of the organism. This raises a concern that such a pair of complementary functions, whose genes are commonly inter-linked by functional associations (here, inferred from neighborhoods), might be instead better merged into a single large functional group, which would better reflect biological reality. In other words, are these complementary function pairs seen by evolution simply as a single function? An argument against this is that such pairs of gene functions are not commonly annotated to the same gene families (low Jaccard index of example GO terms in Table 1; exhaustive list in Supplementary material 5), even if they do commonly occur in the neighborhood of each other. More generally, all significantly enriched pairs of GO terms tend to overall have very low overlap, with 92% of these pairs having Jaccard index <0.1 in genes assigned to them (Supplementary 1, Figure S15).

Turning to the example of the E. coli *lac* operon and the corresponding functions “*carbohydrate transport*” and “*carbohydrate metabolism*”: it is evident that these molecular functions must be distinct due to a different molecular basis for transmembrane transport and for enzymatic cleavage. It is also clear from genomic data that the functions are distinct because they occur independently i.e. they do not co-occur in the same gene families (0 co-occurrences out of 3475 examined prokaryotic COGs; 2.63 co-occurrences of the two functions expected at random, given 199/3475 COGs annotated with “*carbohydrate transport*” and 46/3475 annotated with “*carbohydrate metabolism*”). Given the knowledge of the *lac* operon, it is plausible that many other similar neighborhoods exist that incorporate both transport and metabolism of compounds, reaping benefit from coregulated expression of such complementary gene functions. The NFP approach for predicting gene function and phenotype is able to leverage such systematic co-occurrences, and propagate such dissimilar but co-occurring gene functions across neighborhoods in a rigorous manner.

Such pairs of putatively complementary gene functions might be thought of as child functions of a single hypothetical parent function, which currently does not exist in the Gene Ontology graph -- if it did, then the semantic similarity statistic would mark this pair of functions as closely related. This suggests a possible manner of enhancing of the GO graph based on this data, which would involve creating additional parent nodes that bridge the semantically distant, but biologically related GO terms. Of note, there were previous suggestions to derive alternate GO graphs by drawing on co-occurrence of function annotations in the same genes^24^, which is distinct from what we report here. The complementary functions we propose do not co-occur in the same genes, but are instead associated with each other (in the current analysis, *via* conserved gene neighborhoods, but it is conceivable that functional interactions inferred from other large-scale data might yield similar results). Past work^31^ has proposed that some GO terms may be considered ‘classes’, whose member genes are densely interconnected by functional associations inferred from large-scale data, and other GO terms are ‘categories’, whose members are not linked by functional associations, and which therefore represent artificial concepts. Here, we see widespread evidence for a third type of pattern in the GO graph of gene functions, wherein pairs of distant GO terms are linked by prevalent functional associations bridging the two GO terms. This suggests that such pairs (and possibly larger groups) of GO terms that are significantly interlinked provide a biologically meaningful manner for organizing the catalogue of gene function, with practical implications for automated inference of gene function or phenotype from genomic data.

## Methods

### Methodology overview

In this work, we assess a novel gene neighbourhood representation, called “Neighbourhood functional profile” (NFP), for gene function prediction in 1669 prokaryotic organisms, 49 fungal and 80 metazoan organisms. To predict gene functions, we used Clusters of Orthologous Groups^32^ (COGs and NOGs) gene families, derived from Eggnog database^33^ (version 4.0 for prokaryotic and 5.1 for eukaryotic organisms), henceforth collectively referred to as COGs. We have assigned functions from Gene Ontology^34^ to gene families (COGs) as those occurring in at least 50% genes assigned to a given COG. The resulting datasets contain: a) 3475 COGs (entities) and 1048 GO functions (targets) obtained from the prokaryotic genomes, b) 15969 COGs and 2617 GO functions obtained from the fungal genomes, c) 9187 COGs and 2336 GO functions obtained from the metazoan genomes.

### Methodology description

For a given set of genes **G** and a set of COGs *Ω*, we define a mapping γ: ***G*** → ***P***(***Ω***) which assigns each gene to one or more corresponding COGs. Similarly, for a set of gene functions ∑, mapping *δ*: *Ω* → *P*(*∑*) mapps OGs to a corresponding set of gene functions contained within GO ontology.

For a given set of organisms Θ, a set of genes **G**, a selected gene *g*_*i*_ ∈ Θ_*l*_ a number of neighboring positions from either side of the gene k ∈ ℵ, a set of COGs *Ω* and a set of GO functions *∑*, a functional gene neighbourhood of *g*_*i*_ in the organism *Θ*_*l*_ is defined as a count of all GO functions occurring in COGs or NOGs assigned to genes in its k-neighbourhood. Formally: 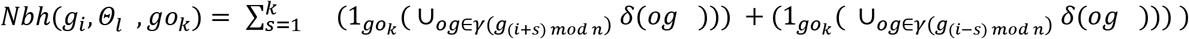, for prokaryotic organisms and 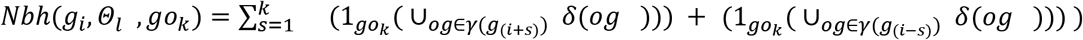, for eukaryotic organisms where *g*_*s*_ are genes contained in a k-neighbourhood of gene *g*_*i*_ and *n* denotes the total size of a genome. Thus, the result of gene neighbourhood computation for all go functions is a tuple of size |*∑*|containing corresponding occurrence frequencies of all GO functions in the k-neighbourhood of *g*_*i*_. In our data analyses, we use COGs as entities, thus each COG is associated with a vector containing |*∑*| elements, corresponding to occurrence of GO functions in its k-neighbourhood, derived from neighbourhoods of corresponding genes. The neighbourhood frequencies computed for a gene *g*_*i*_ are added to the frequency vector of all OGs such that 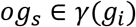. Thus 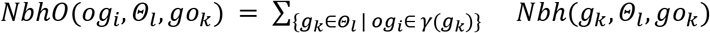 and the final features are computed as 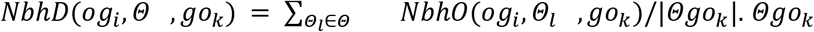 denotes a set of all organisms containing at least one COG with function *go*_*k*_. Note that genes that are not assigned to any COG do not add to function frequency count in the neighbourhoods.

We trained a Random Forest of Predictive Clustering Trees^35^ model on this features to predict gene functions. Performance of this methodology is compared to the biological (1-NN) and Gaussian Field Label Propagation model^36^, trained on the average of logarithmic distances of pairs of COGs in different organisms. For a pair of genes 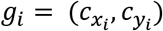 and 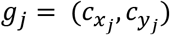, where 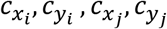 are coordinates of corresponding genes in a genome. The logarithmic distance of two genes contained in a prokaryotic organism is computed as: 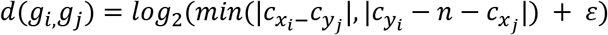, if 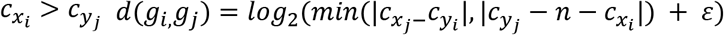, if 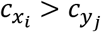

If two genes are overlapping, we define their distance to equal small constant ε. In our work, we use ε = 10^−10^. Distances in eukaryotic organisms are computed as:

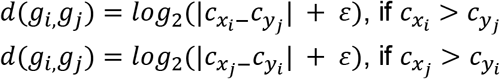

The average logarithm distance of pair of OGs is computed as: 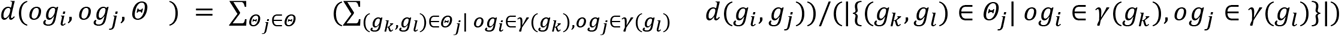

An overview of the approach is provided in Figure 7.

**Figure 7.**
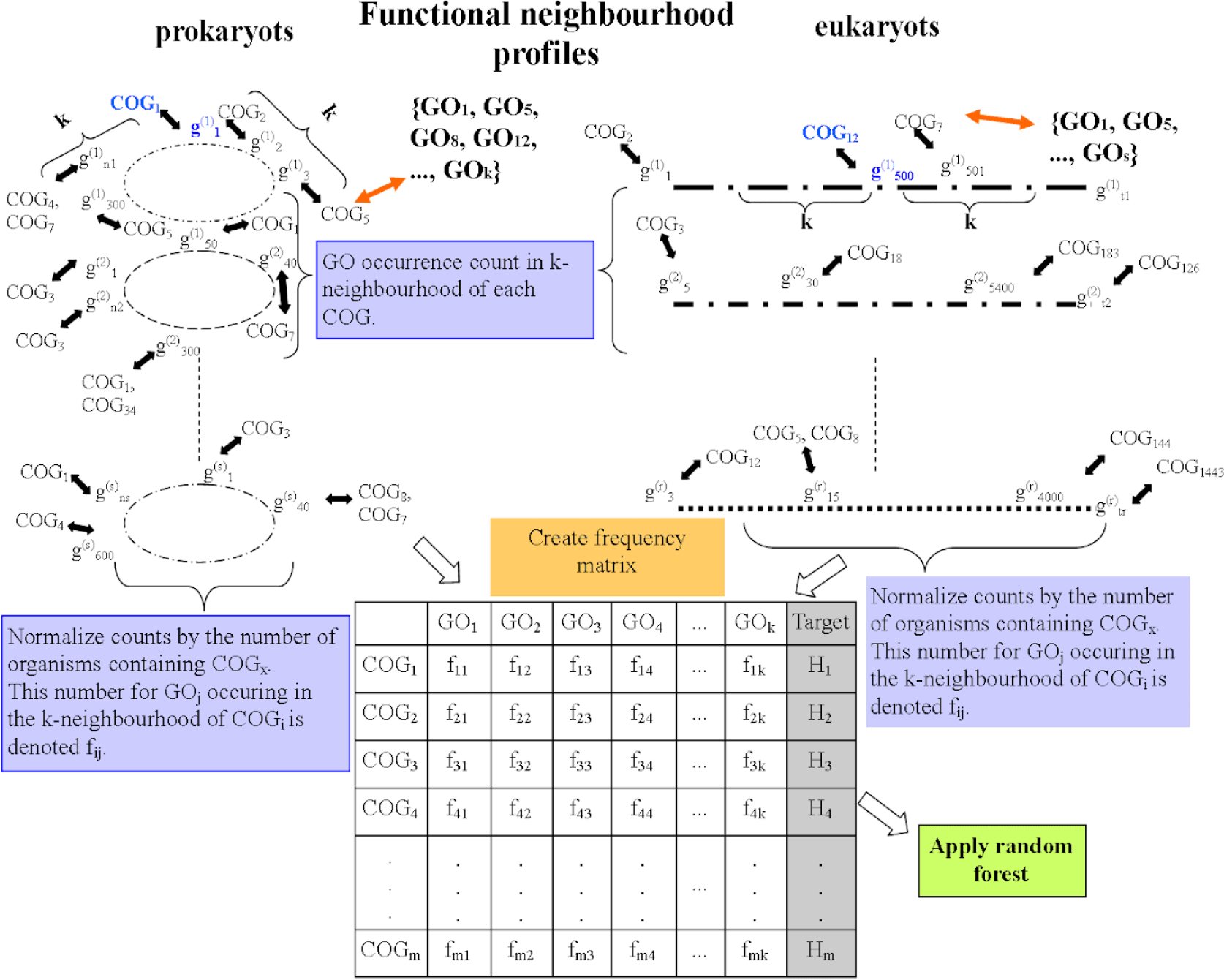
Overview of the neighborhood function profiles methodology. Location-based approaches are trained on pairwise COG/NOG distances of corresponding genes contained within genome of different prokaryotic and eukaryotic organisms (as demonstrated in a). The obtained distances are used to create a similarity table to train the k-NN model and the association network to train the Gaussian Field Label Propagation approach. Functional neighbourhoods (b) are used to create a normalized frequency matrix which is used to train the Random Forest of Predictive Clustering trees model. “COG” in the Figure is used to denote both COG and NOG. Target H_i_ denotes the sub hierarchy of GO ontology associated with COG_i_ (sub hierarchy contains information about the GO functions assigned to some COG and parent-child relations between these GO functions).

### Method evaluation and relevance

All methods are evaluated in cross-validation setting (k-NN and GFP using leave-one-out cross-validation and NFP using out-of-bag estimates using a forest of 600 trees). Out-of-bag estimate is used in NFP since it significantly reduces validation time of a random forest model and provides comparable estimate of error to cross-validation.

Since assessing the importance of enriched functions in genomic neighborhoods for gene function prediction was a central goal of our work, we evaluated our approach (that utilizes this information) by comparing it with other state-of-the-art approaches that use information about gene proximity, but lack the information about the enriched semantically distant functions. Increasing predictive performance of genomic neighborhood-based methods may significantly advance our knowledge about the many genes of unknown function, particularly since these methodologies can be easily incorporated into an ensemble model with other genomic predictors, which -- due to complementarity of predictions of different models -- yields superior performance^37^.

### Association testing of GO functions in neighborhoods

To measure the strength of association between different pairs of GO functions from our data, we first computed the contingency tables that contain the following components: a) COG contains GOx and Neighborhood contains GO_y_, b) COG contains GO_x_ and Neighbourhood does not contain GO_y_, c) COG does not contain GO_x_ and Neighbourhood contains GO_y_, d) COG does not contain GO_x_ and Neighbourhood does not contain GO_y_.

From these tables, we compute the Odds ratio 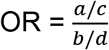 and ultimately the Log Odds ratio *log*_2_(*OR*). In addition to computing ORs and log odds ratios and testing its statistical significance using the Fisher exact test (for ORs) and z-test for*log*_2_(*OR*) > 0. We also provide empirical evidence of strength of association by computing the number (percentage) of significantly enriched pairs of functions computed on the original dataset with higher or at least two-times higher (2x or more) *log*_2_(*OR*) than the corresponding pair computed on the randomized dataset (gene locations are permuted in the genome).

For a GO term with frequency p(GO), the information content (IC) of a GO is defined as −*log*_2_(p(*GO*)). The Resnik Similarity (RS) of a pair of GO terms *GO*_*x*_, *GO*_*y*_ is defined as *RS*(*GO*_*x*_, *GO*_*y*_), where *comAnc* denotes the most informative common ancestor (the one with largest IC). We use RS<2 as a criteria for selecting pairs of distant GO functions for which we compute enrichments in Eukaryotic organisms, the reason for this choice is that GO frequencies on eukaryotic organisms are empirically computed from the data, thus upper level (children of the root) of GO ontology have Information Content higher than 1.

### Association testing of GO functions in neighborhoods

In the Discussion section of this text, for some function *GO*_*x*_ we say that *GO*_*y*_ is a “related function” if: RS(*GO*_*x*_, *GO*_*y*_)≥6 for *GO*_*x*_, *GO*_*y*_ contained in the same namespace of the GO ontology, or if J(*GO*_*x*_, *GO*_*y*_)≥0.6 for *GO*_*x*_, *GO*_*y*_ not contained in the same namespace of GO. We observe 6615 related pairs of functions with FDR<1% in prokaryotic dataset. For some function *GO*_*x*_, we say that *GO*_*y*_ is “unrelated function” if RS(*GO*_*x*_, *GO*_*y*_)<1, for *GO*_*x*_, *GO*_*y*_ contained in the same namespace of the GO ontology, or if J(*GO*_*x*_, *GO*_*y*_)<0.05 for *GO*_*x*_, *GO*_*y*_ not contained in the same namespace of GO. We observe 204202 such pairs with FDR<1% in prokaryotic dataset. These statistics are (47.5% - 8853 pairs, 97% - 878850) for Fungi and (99.9% - 36122 pairs, 100% - 1256151) for Metazoa.

The 11 gene functions analyzed individually in the manuscript are all taken from the same namespace of GO ontology, to prevent considering potentially synonymous functions from different namespaces as semantically distant.

For a given mapping 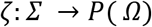, that maps a GO function to a set of COGs which contain this function, we use 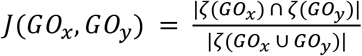 to measure the level of circularity of pairs of functions (especially these from different namespaces of GO ontology).

The information accretion of a function *GO*_*x*_ is computed as 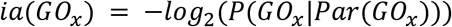, where Par denotes a set of parent nodes of *GO*_*x*_. This implies that if some model predicts GO function that does not have high probability of occurrence, given the parent nodes, it gets significantly larger accretion score.

### The average number of annotations per annotated gene

To assess the potential reason for a difference in predictive power of all models between Fungi and Metazoa dataset, we computed the average number of annotations per annotated gene. This information is important since it has direct impact on the stability and the amount of information contained in gene functional neighbourhoods.

### Dataset for predicting phenotypes

The NFP dataset used to predict phenotypes in different strains of *E. coli* was constructed so that E. coli strains form examples (entities), whereas features are COG/NOG-GO function pairs (frequency of occurrence of each GO in the neighbourhood of each COG/NOG contained in each *E. coli* strain). Thus, the whole table depicted in Figure 7 is contained in each row of our NFP dataset for phenotype prediction. This features (sparse in nature) were used to create 228 principal components using PCA algorithm.

### Target variables in analyses of phenotypes

The Phenotypic dataset contains 151 target variables (phenotypes) that denote if a defect has been detected (value 1) after application of specific combination of drugs or not (value 0).

## Author Contributions

MM performed all analyses and devised methodological improvements. TS and FS conceived and supervised the project. All authors drafted the manuscript.

## Data Availability

Data is available via the supplementary material of this publication or upon request from authors.

## Acknowledgements

We thank Jelena Repar and Ivan Erill for helpful discussions. This work was funded by the ERC Starting Grant “HYPER-INSIGHT” (757700), the Spanish MINECO grant “RegioMut” (BFU2017-89833-P) and the Croatian Science Foundation grant “Descriptive Induction” (HRZZ-9623). F.S. is funded by the ICREA Research Professor programme. F.S. acknowledges support of the Severo Ochoa Centres of Excellence programme to the IRB Barcelona. A part of this work was performed while M.M was a visiting researcher at the IRB Barcelona.

