## Supplementary Text for "Patterns of diverse gene functions in genomic neighborhoods predict gene function and phenotype"

**Supplementary material for Mihelcic, Smuc and Supek (2019)**

This document describes additional analyses related with the Neighborhood function profiles (NFP) representation of genomic data, the prevalence of semantically distant GO functions in genome neighborhoods, and the impact of this representation on machine learning for the task of predicting gene function and phenotype.

### S1. Data

#### S1.1 Genomes used in the analyses

1669 prokaryotic genomes obtained from NCBI genomes database[1], 49 fungal and 80 metazoan genomes obtained from ensemble genomes database[2]. In this study, we used fully sequenced genomes containing gene locations on the main chromosome and plasmids (on prokaryotic organisms) and gene locations on all chromosomes and non-chromosomal region in eukaryotic organisms.

#### S1.2 Information about Clusters of Orthologous groups

We used COG (clusters of orthologous groups) and NOG (non-supervised orthologous groups) gene families in prokaryots and their equivalent fuNOG and (meNOG)[3] in eukaryotic organisms. We assigned genes to which a mapping to COG or NOG was known and used COGs and NOGs to assign functions to genes. Unassigned genes can be first compared to assigned genes via blast or similar techniques (such as the eggnog mapper[4]) and can be assigned to COG/NOG. Genes not belonging to any existing COG or NOG would need to be grouped in newly defined NOGs. Our prokaryotic dataset contained 3475 COG/NOGs, fungi dataset contained 15741 fuNOGs and metazoan dataset contained 9185 meNOGs.

#### S1.3 Gene Ontology (GO)

Each COG/NOG was assigned a set of gene functions, herein represented by Gene Ontology (GO) terms[5]. We tested two scenarios: assigning a function to a COG or NOG when 50% of genes in a COG/NOG contain this function and assigning a function to a COG or NOG when 30% of genes in that COG/NOG contain this function. The main conclusions stated in the manuscript are supported using both thresholds.

### S2. Method

The depiction of location-based and function-based representations (*neighborhood function profiles*, NFP) is given in Figure S1.

---

[1] https://www.ncbi.nlm.nih.gov/genome
[2] http://ensemblgenomes.org/
[3] http://eggnogdb.embl.de/#/app/downloads
[4] http://eggnogdb.embl.de/#/app/emapper
[5] http://geneontology.org/

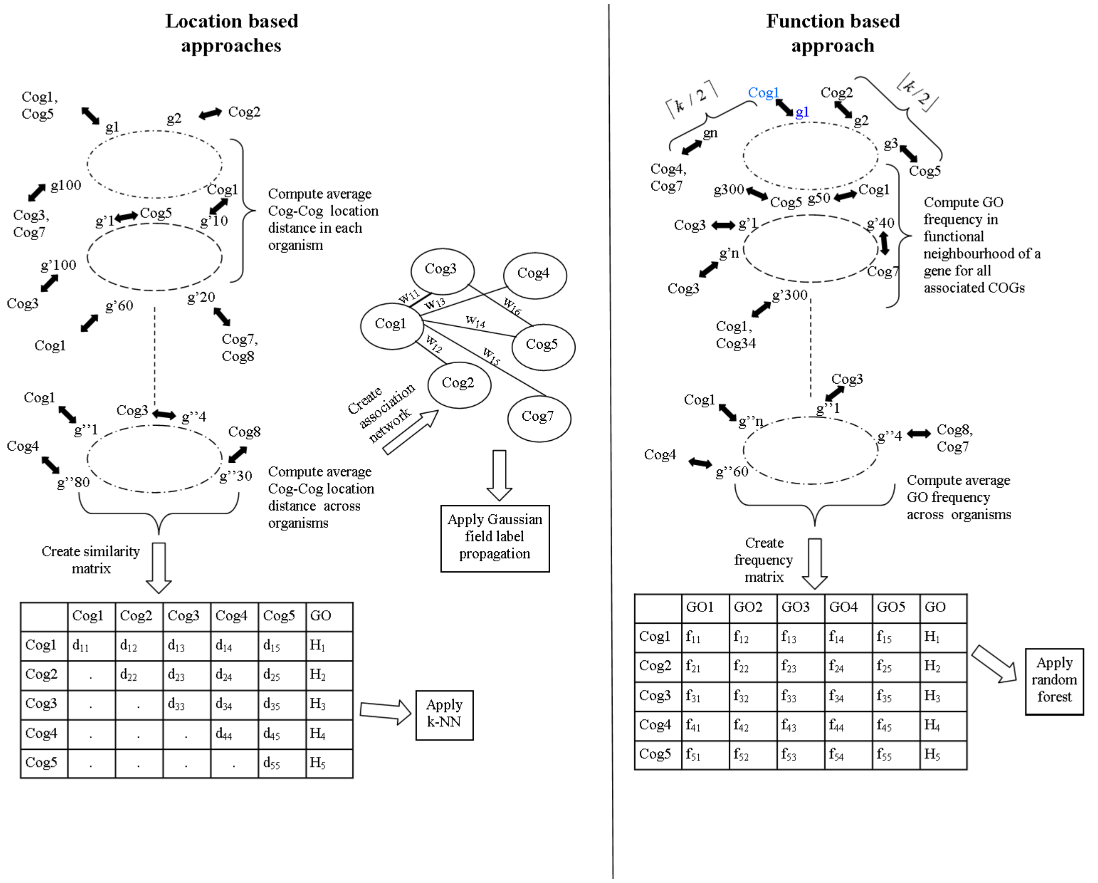

**Figure S1.** Location-based approaches are trained on pairwise OG distances of corresponding genes contained within genome of different prokaryotic (as demonstrated in this figure) and eukaryotic organisms. The obtained distances are used to create a similarity table to train the k-NN model and the association network to train the Gaussian Field Propagation (GFP) approach. Functional Neighborhoods are used to create a normalized frequency matrix which is used to train the Random Forest of Predictive Clustering trees model. In all experiments on bacterial organisms, we use a Neighborhood of 2 genes on each side of a targeted gene.

To measure the strength of association between different pairs of GO functions from our data, we first computed the contingency tables that contain the following components:

**Table S1.** Contingency tables used to assess associations between pairs of functions GOx and GOy.

| | Neighborhood contains GOy | Neighborhood does not contain GOy |
|---|---|---|
| OG contains GOx | a | b |
| OG does not contain GOx | c | d |

From these tables, we compute the Odds ratio OR = $\frac{a/c}{b/d}$ and the Log Odds Ratio $log_2(OR)$. In addition to computing ORs and log odds ratios and testing its statistical significance using the Fisher exact test (for ORs) and z-test for$log_2(OR) > 0$. We also provide empirical evidence of strength of association by computing the number (percentage) of significantly enriched pairs of functions computed on the original dataset with higher or significantly higher (2x or more) $log_2(OR)$ than the corresponding pair computed on the randomized dataset (gene locations are permuted in the genome).

For a given mapping $\zeta: \Sigma \rightarrow P(\Omega)$ that maps a GO function to a set of OGs which contain this function, we use $J(GO_x, GO_y) = \frac{|\zeta(GO_x) \cap \zeta(GO_y)|}{|\zeta(GO_x \cup GO_y)|}$ to measure the level of circularity of pairs of functions (especially these from different namespaces of GO ontology).

The experiments of assessing the influence of distant and enriched functions to the predictive performance of the NFP methodology, as described in section "Predictive power of the Neighborhood Function Profile (NFP) classifier" of the main manuscript were performed in the following manner:

a) Features were divided into several categories (detailed description available in section: "Predicting gene function from genomic neighborhoods can be greatly improved by taking semantically dissimilar functions into account ".
b) To avoid positive bias introduced with a feature selection procedure (especially in case of sets of attributes containing semantically close functions - CLPar and CL categories), we added a number of randomly generated attributes, so that the total number of attributes equals the overall number of attributes used in the experiment (|BPP|). These attributes were generated using randomly generated numbers from uniform distribution in the interval $[At_{min}, At_{max}]$.
c) The increase of predictive power of using one set of attributes over CLPar - usually used in guilt-by-association approaches, demonstrates the added benefit of enriched and semantically distant functions for gene function prediction using NFP methodology.

### S3. Experiments

#### S3.1 Log Odds Ratio analyses

We have computed Odds Ratio [Cornfield] and Logs Odds Ratio for all pairs of 1048 GO functions associated with OGs in bacterial genomes. Odds Ratios for pairs of functions $(GO_x, GO_y)$ were computed from gene functional Neighborhoods and they represent the odds of a function $GO_y$ occurring in a Neighborhood of a gene containing some function$GO_x$. The selected subset of these functions, containing only GO functions from Biological Process ontology and a prokaryotic subset[6] contains 478 different GO functions. The resulting Log Odds Ratios for all pairs of these 478 functions, clustered by log odds ratio (Figure S2) and Resnik semantic similarity [Resnik] (Figure S3) show existence of several clusters of highly associated functions.

[6] http://www.geneontology.org/page/go-subset-guide

Moreover, it can be seen from heatmap shown in Figure S3 that there exist many pairs of highly associated functions that are semantically dissimilar (those far away from the heatmaps diagonal). It can be also seen that there is a significant amount of highly associated pairs of functions (those with logOR > 1).

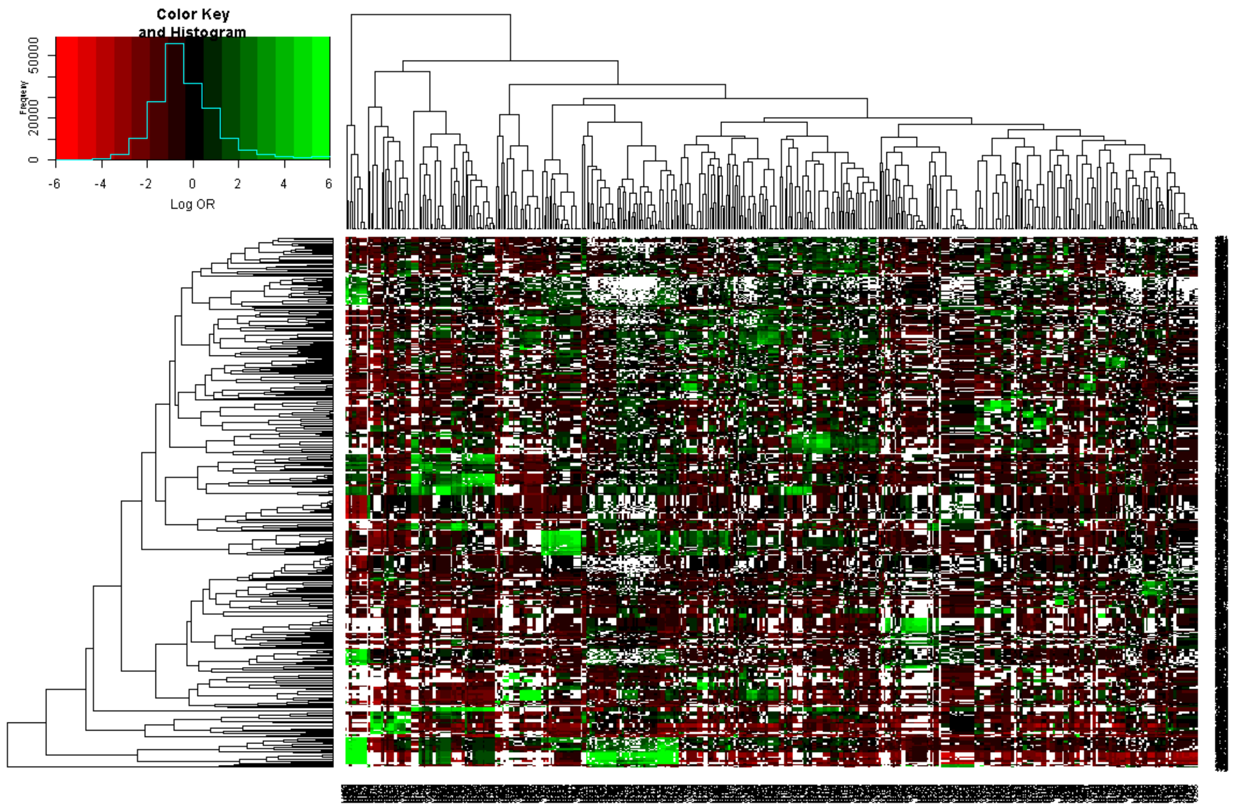

**Figure S2.** Heatmap of pairwise GO function Log Odds Ratios, derived from prokaryotic genomes. Heatmap rows and columns have been clustered by Log Odds Ratio. White color denotes insignificant associations (p-value > 0.05).

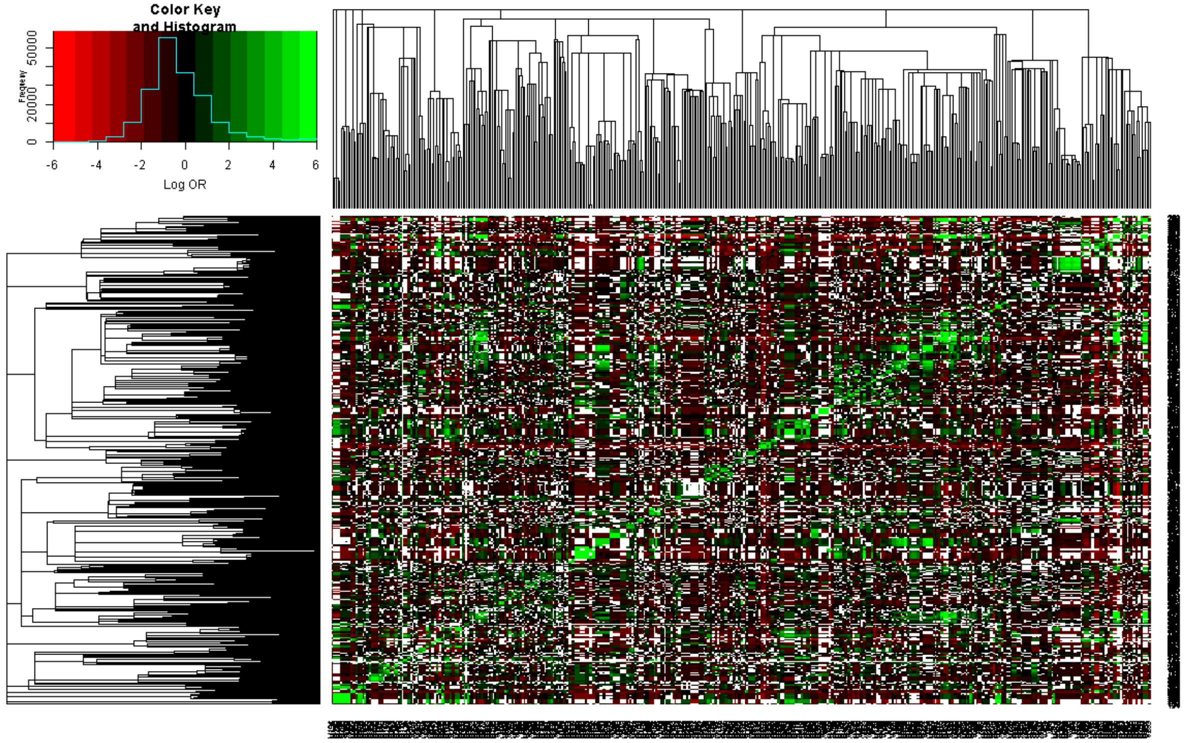

**Figure S3.** Heatmap of pairwise GO term log odds ratios (OR), quantifying enrichments in genomic neighbohoods of prokaryotic genomes. Rows and columns have been clustered by GO semantic similarity. White color denotes non-significant associations (p > 0.05).

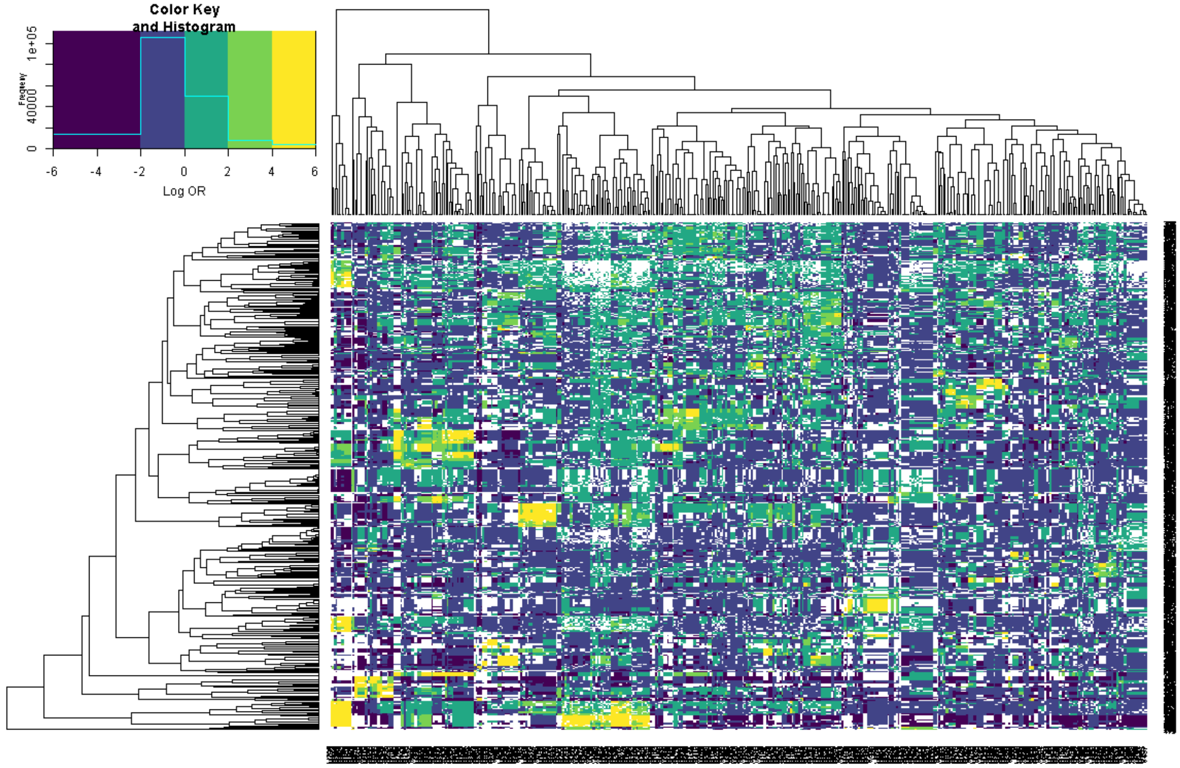

**Figure S4.** Heatmap of pairwise GO term log odds ratios (OR), quantifying enrichments in genomic neighbohoods of prokaryotic genomes. Rows and columns have been clustered by log OR profiles. White color denotes non-significant associations (p > 0.05).

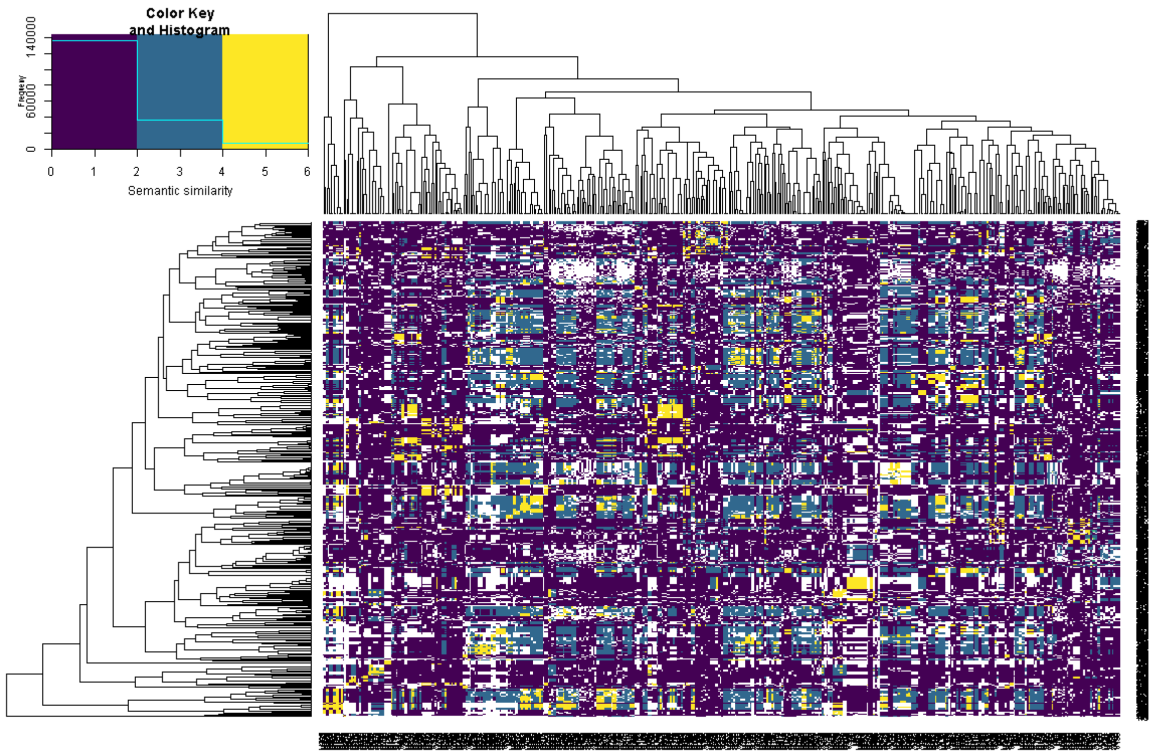

**Figure S5.** Heatmap of pairwise GO term semantic similarites, for GO terms arranged as in previous figure (clustered by log OR profiles in prokaryotic genomes). White color denotes non-significant associations (p > 0.05).

We can see from Figures S4 and S5 that clusters of semantically close functions (denoted in yellow in Figure S5) are mostly significantly enlarged when looking at Log Odds Ratios in Figure S4. This means that there exist a high number of pairs, of semantically dissimilar functions, that are highly enriched in functional Neighborhoods.

Similar thing can be seen from Figures S2 and S3, where many clusters occurring in Figure S3 are significantly smaller than these from Figure S3, which shows that semantically dissimilar functions must be a part of clusters containing highly enriched pairs of functions (these with logOR>0).

In the continuation, we present several GO functions that have highly enriched dissimilar function in their functional Neighborhood (for all selected pairs see Supplementary document 5). For each pair of enriched, dissimilar functions, we report: a) the Log Odds Ratio of occurrence of a selected function GOs in its own Neighborhood, b) the Log Odds Ratio of occurrence of a dissimilar function GOm in the functional Neighborhood of a function GOs, c) the statistical significance of the enrichment, computed by Fishers exact test [Fisher], d) Resnik Semantic Similarity [Resnik] and e) the Jaccard index [Jaccard] of occurrence of these GO terms in OGs present in the dataset.

**Table S2.** Selected subset of highly enriched pairs of GO functions with corresponding Log Odds Ratio along with the corresponding confidence interval, statistical significance of association, semantic distance and the corresponding Jaccard index computed on function co-occurrence in different COGs from our prokaryotic data set.

| GOs | Description | LOR GOs-GOs | GOm | Description | LOR GOs-GOm | p | Res. Sem. Simil. | J | Jrand |
|---|---|---|---|---|---|---|---|---|---|
| GO0005975 | carbohydrate metabolic process | 2.53∓ 0.01 | GO0008643 | carbohydrate transport | 2.12 ∓ 0.02 | 0.0 | 0.0 | 0.0 | 0.004 |
| GO0006260 | DNA replication | 2.99∓ 0.02 | GO0032506 | cytokinetic process | 1.41 ∓ 0.06 | 0.0 | 0.787 | 0.0 | 0.0 |
| GO0006974 | cellular response to DNA damage stimulus | 2.39∓ 0.02 | GO0006265 | DNA topological change | 1.42 ∓ 0.06 | 0.0 | 0.787 | 0.0 | 0.0 |
| GO0006974 | cellular response to DNA damage stimulus | 2.39∓ 0.02 | GO0033866 | nucleoside bisphosphate biosynthetic process | 1.41 ∓ 0.05 | 0.0 | 0.787 | 0.0 | 0.0 |
| GO0016051 | carbohydrate biosynthetic process | 4.16∓ 0.02 | GO0043163 | cell envelope organization | 2.1∓ 0.07 | 0.0 | 0.0 | 0.0 | 0.0 |
| GO0006457 | protein folding | 5.41∓ 0.02 | GO0016226 | iron-sulfur cluster assembly | 1.91 ∓ 0.08 | 0.0 | 0.521 | 0.0 | 0.0 |
| GO0046700 | heterocycle catabolic process | 1.15∓ 0.01 | GO0051180 | vitamin transport | 1.65 ∓ 0.08 | 0.0 | 0.01 | 0.01 | 0.0 |
| GO0006310 | DNA recombination | 3.09∓ 0.02 | GO0006952 | defense response | 1.02 ∓ 0.13 | 0.0 | 0.0 | 0.0 | 0.0 |
| GO0046903 | secretion | 6.56∓ 0.03 | GO0006935 | chemotaxis | 3.6∓ 0.05 | 0.0 | 0.0 | 0.0 | 0.0 |
| GO0008610 | lipid biosynthetic process | 3.21∓ 0.02 | GO0051668 | localization within membrane | 2.43∓ 0.05 | 0.0 | 0.0 | 0.0 | 0.0 |
| GO0006865 | amino acid | 2.32∓ | GO0009310 | amine | 2.17∓ | 0.0 | 0.0 | 0.0 | 0.0 |

| | | | | | | | | | |
|---|---|---|---|---|---|---|---|---|---|
| | transport | 0.04 | | catabolic process | 0.13 | | | | |
| GO0006508 | proteolysis | 1.80 ∓ 0.02 | GO0019682 | glyceraldehyde-3-phosphate metabolic process | 1.48 ∓ 0.05 | 0.0 | 0.95 | 0.0 | 0.0 |

For each selected function GOs, we plot a histogram, showing the frequencies of Log Odds Ratios of the distant, medium-distant functions as well as all immediate parents of a selected GO term (denoted CLPar). Histograms can be seen in Figures S6.

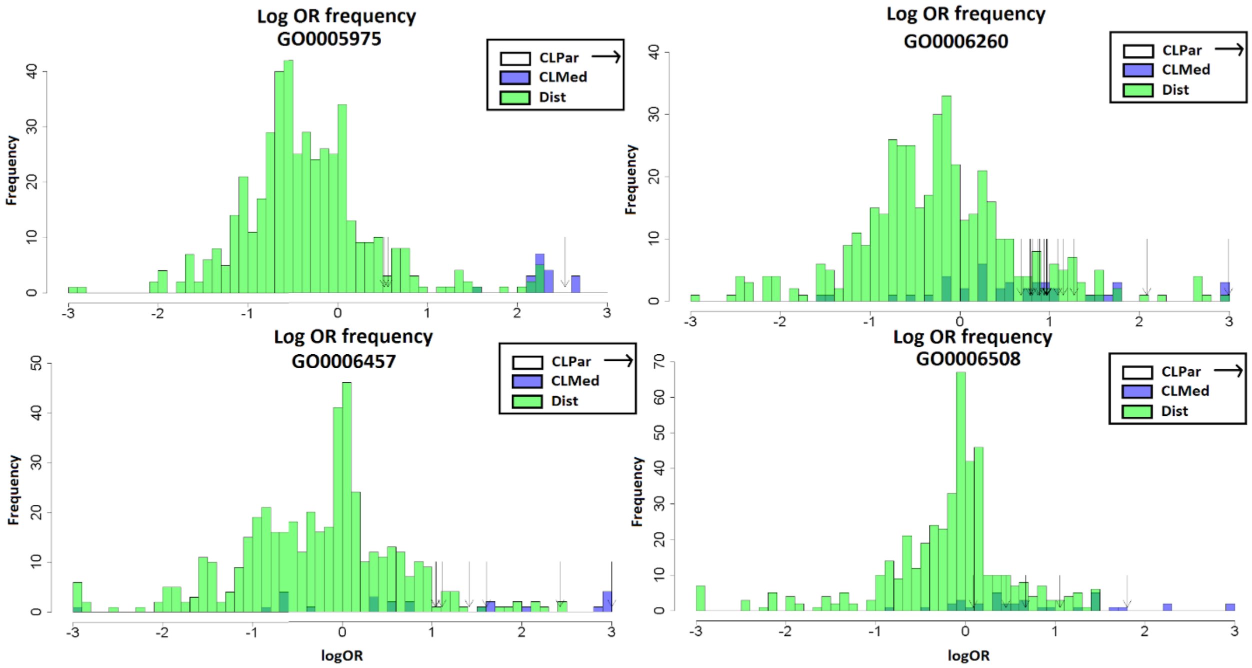

**Figure S6.** Histograms showing Log Odds Ratio frequency of GO functions and a selected GO function divided by semantic similarity to CLPar (selected function + all parent functions in GO Ontology), CLMed (functions with Resnik semantic similarity > 2 with the selected function) and Dist (functions with Resnik semantic similarity ≤2 with the selected function).

We also plot a histogram showing frequencies of Log Odds Ratios of distant and close functions as well as statistical significance of Log Odds Ratio of occurrence of function GOy in the functional Neighborhood of function GOx to be greater than zero, and the significance of this Log Odds Ratio of being larger than the Log Odds Ratio of GOx occurrence in its own functional Neighborhood. Histograms can be seen in Figure S7.

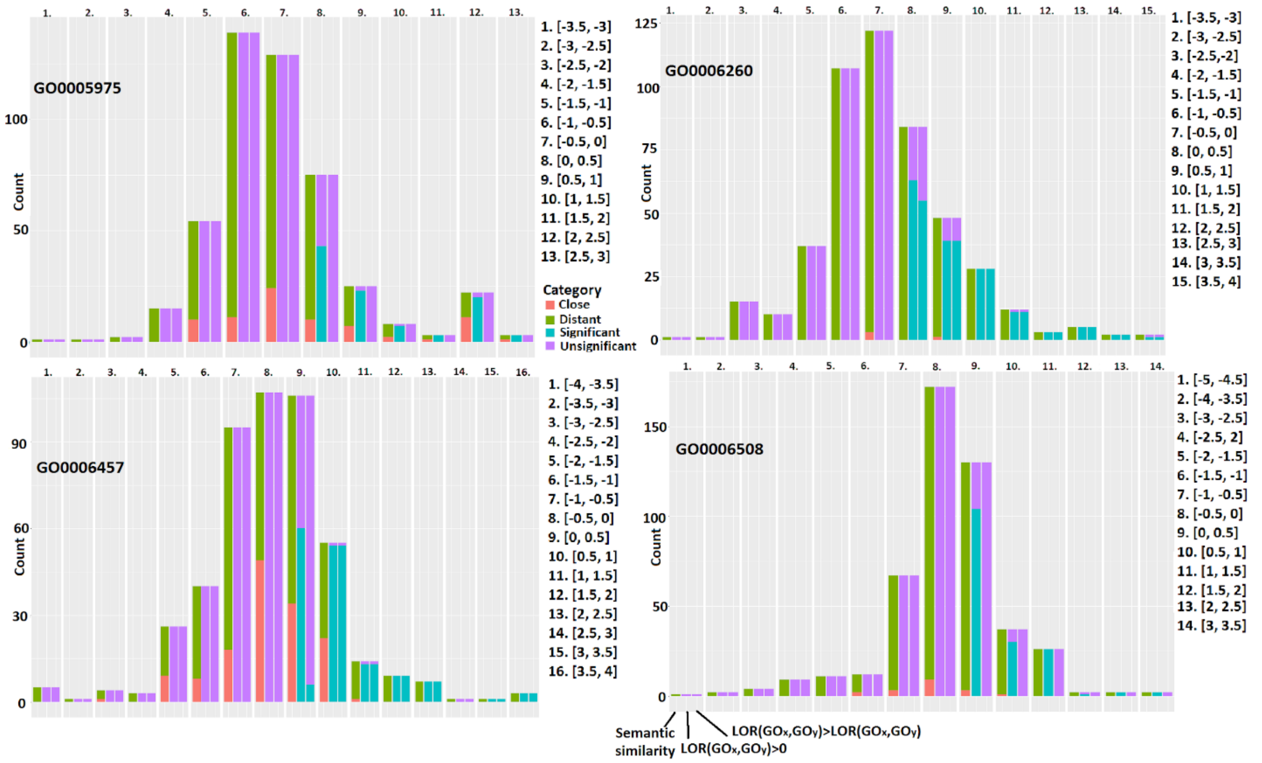

**Figure S7**. Red color denotes the frequency of semantically close functions to the selected GO category, green color denotes semantically distant functions, blue color denotes frequencies of functions with significant logOR>0 (second column) and logOR(GOy, GOx) > logOR (GOx, GOx), in the third column. Purple color denotes the frequency of insignificant categories. In this consideration, we only test the significance of enrichments, thus all pairs with depletions are deemed insignificant. Intervals not present in the bar plot have 0 frequency.

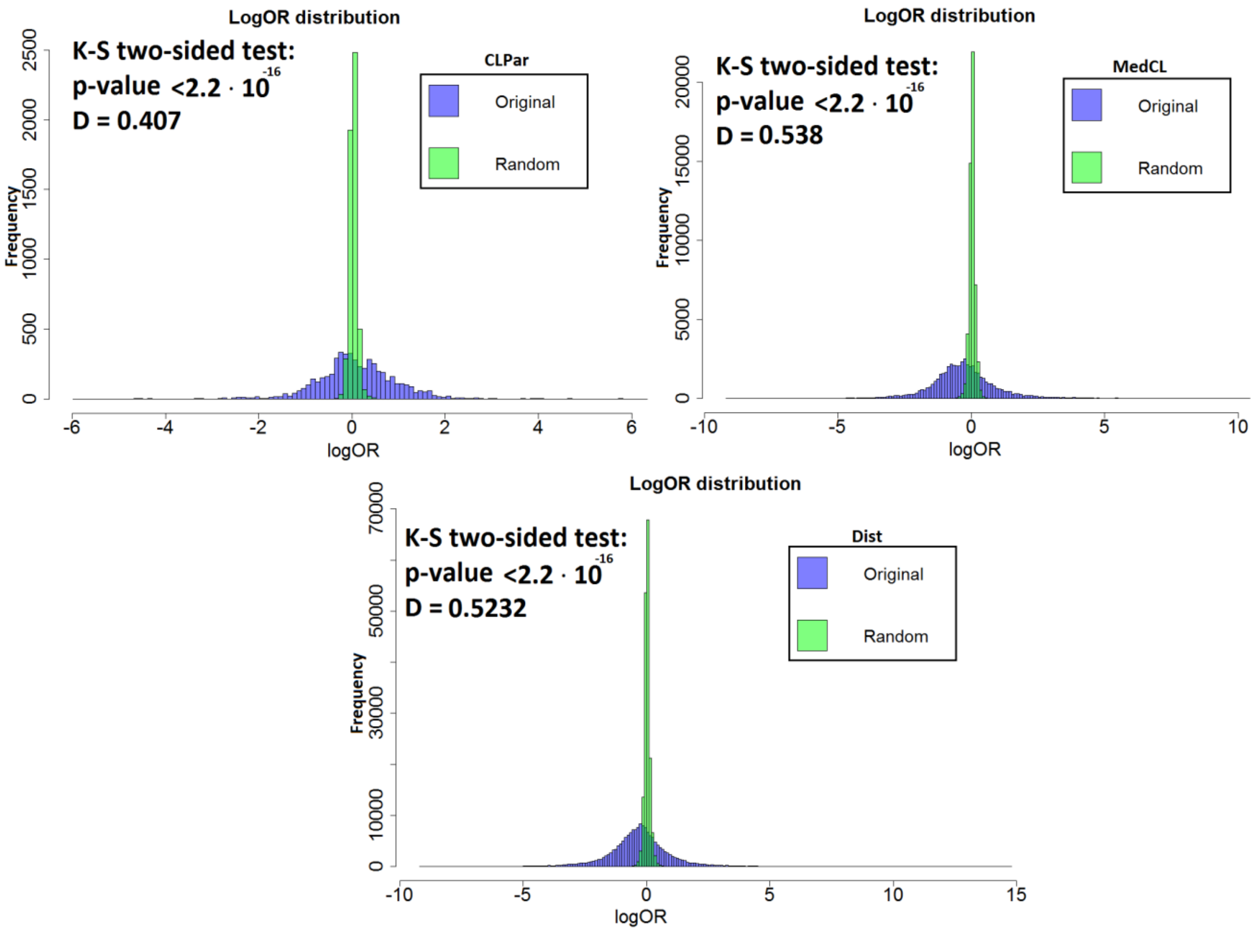

**Figure S8.**. Comparative histograms showing Log Odds Ratios computed on the original prokaryotic genomes (blue) and in randomized versions of these genomes (green).

Finally, we plot the comparative histograms showing distributions of Log Odds Ratios of all pairs of functions, contained in our dataset from the Biological Process Ontology and contained in the Prokaryotic subset, in the original and randomized versions of bacterial genomes. In the randomized version, we randomly permuted gene locations, keeping gene to OG mapping intact. Histograms presented in Figure S8. demonstrate that there is a significant difference (as computed by the two-sided Mann-Whitney U-test) in Log Odds Ratio distribution computed from the original bacterial genomes compared to those computed on the randomized version of the genome.

In the continuation, we plot a Scatter plot for two selected GO functions and its enriched GO pair (presented in Table S2) to analyze the percentage of Neighborhoods, of different COGs, containing each of these functions. We compare this with percentages of Neighborhoods containing each of the selected functions on a randomly permuted genome as a baseline comparison for the same set of COGs (denoted in red).

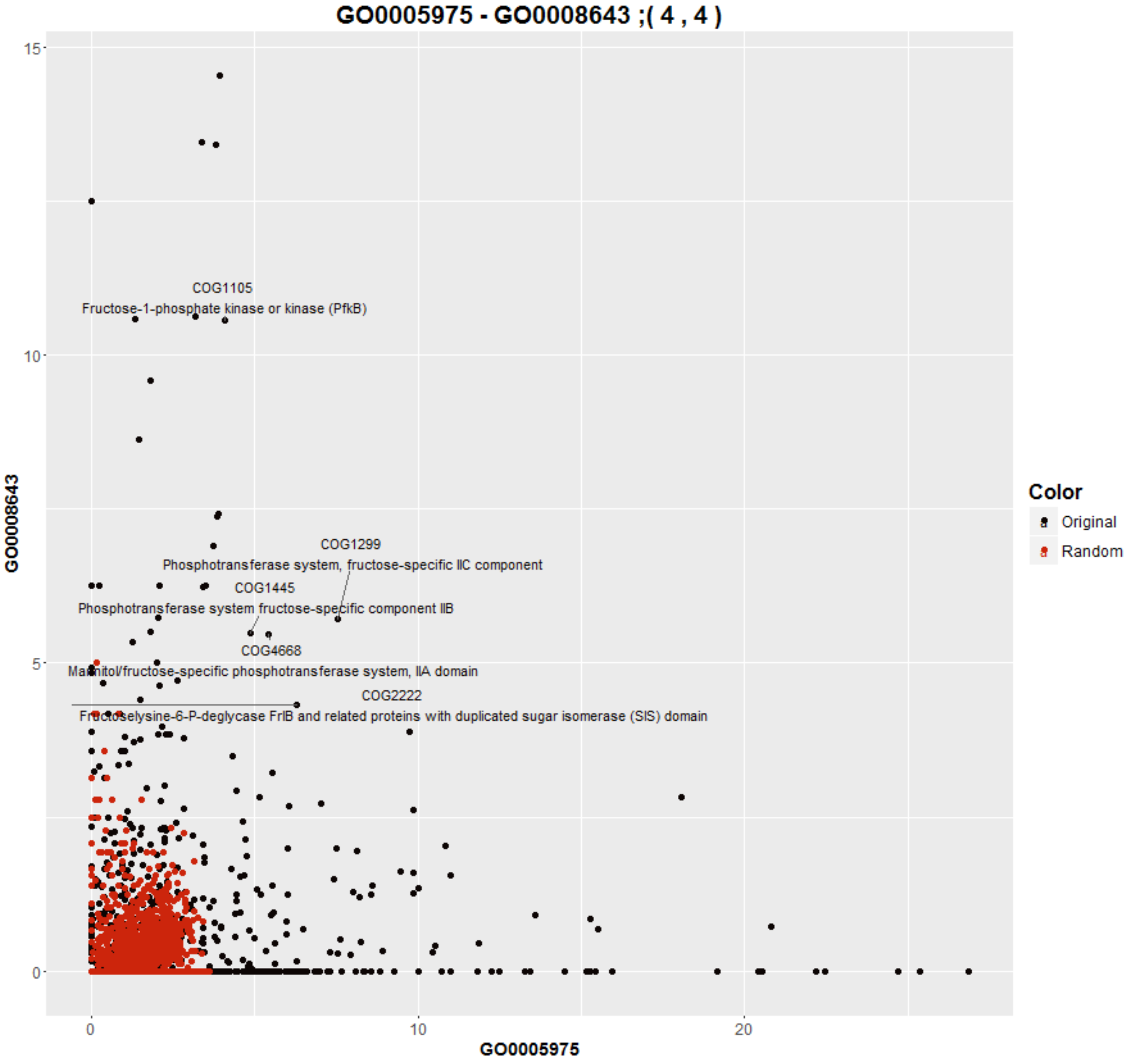

**Figure S9.** Scatter plot showing the average percentage of gene functional Neighborhood of different COGs that contain GO function GO0005975 (x-axis) and GO0008643 (y-axis).

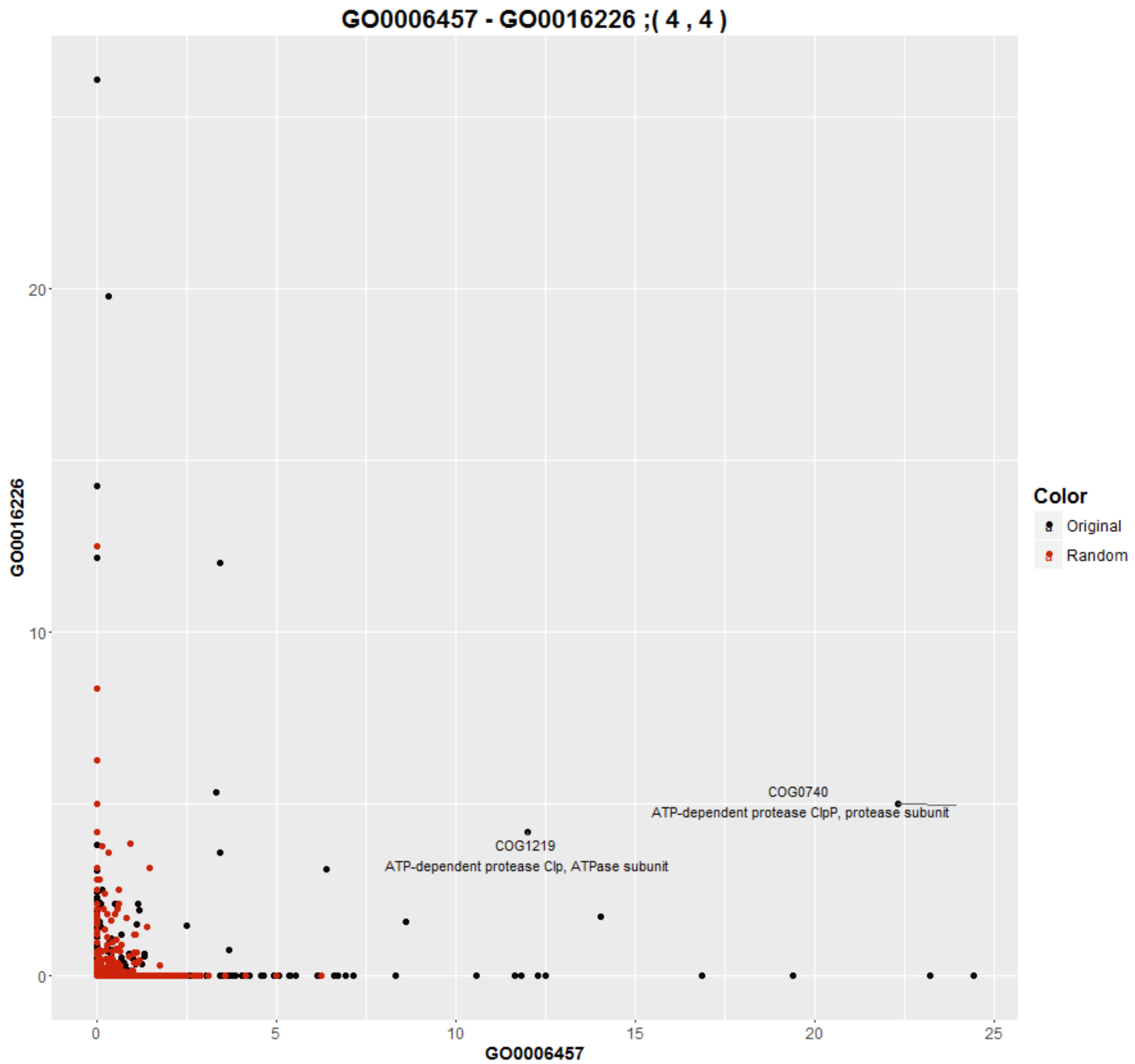

**Figure S10.** Scatter plot showing the average percentage of gene functional Neighborhood of different COGs that contain GO function GO0006457 (x-axis) and GO00016226 (y-axis).

The scatter plots in Figures S9 and S10 arrange COGs by the average percentage of their Neighborhood containing functions GO0005975, GO00008643 (Figure S9) and GO0006457, GO00016226 (Figure S10). COGs whose Neighborhoods contain ≥4% of both GO functions (in average) are pointed out along with their description.

#### Enrichments in Eukaryotes

Similarly as in prokaryotes, the distribution of enrichments on original dataset are significantly different from the enrichment distribution obtained on the randomized dataset.

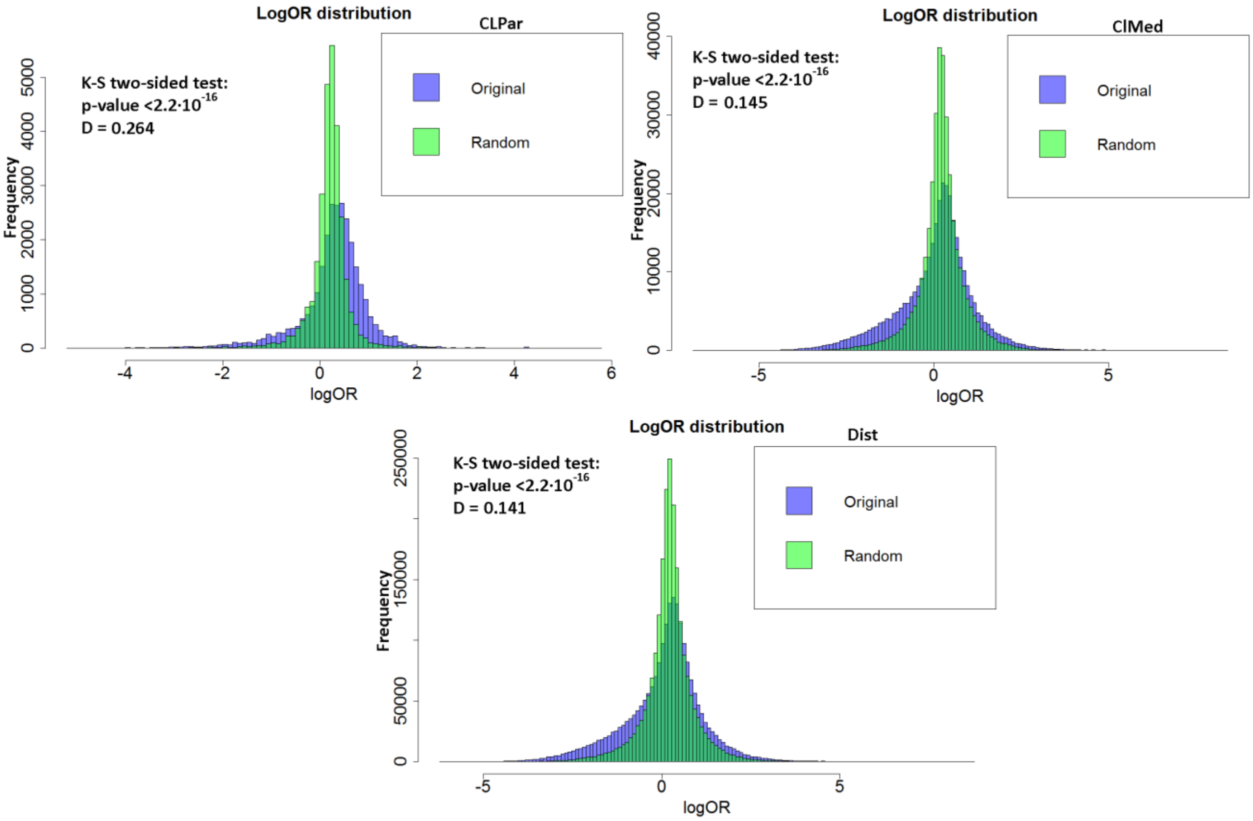

**Figure S11.** Histograms display significant difference in distribution of Log Odds Ratios on original and randomized dataset on Fungi organisms for all groups of functions.

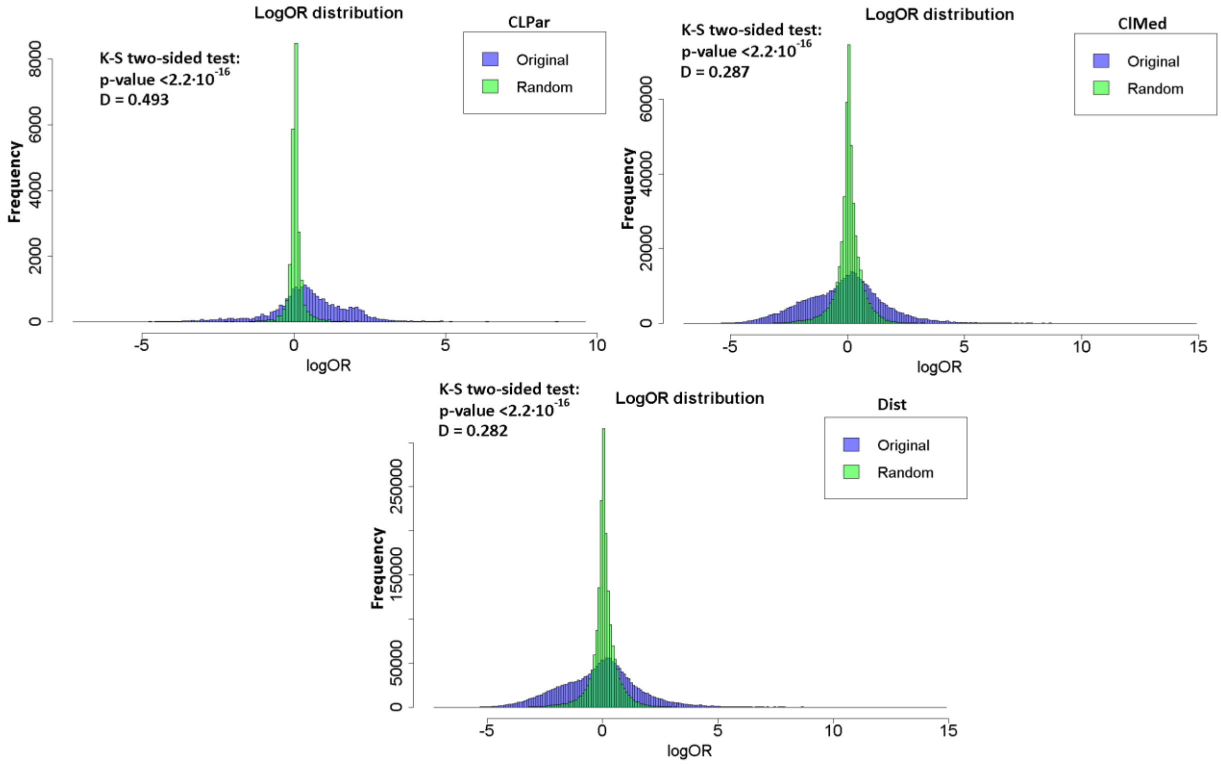

**Figure S12.** Histograms display significant difference in distribution of Log Odds Ratios on original and randomized dataset, on Metazoa organisms, for all groups of functions.

We can see the increased left tail of the original log odds ratio distribution. We show that this is the result of insignificant contingency tables (these having 0 GO co-occurrences or high imbalance of occurence between a pair of GO functions).  This is demonstrated by showing comparative plots of the log odds ratio distribution of all pairs of GO functions and a distribution of pairs of functions having significant contingency tables (as computed by the Fisher's exact test). These can be seen in Figure S13. for Fungi organisms and Figure S14. for Metazoa organisms.

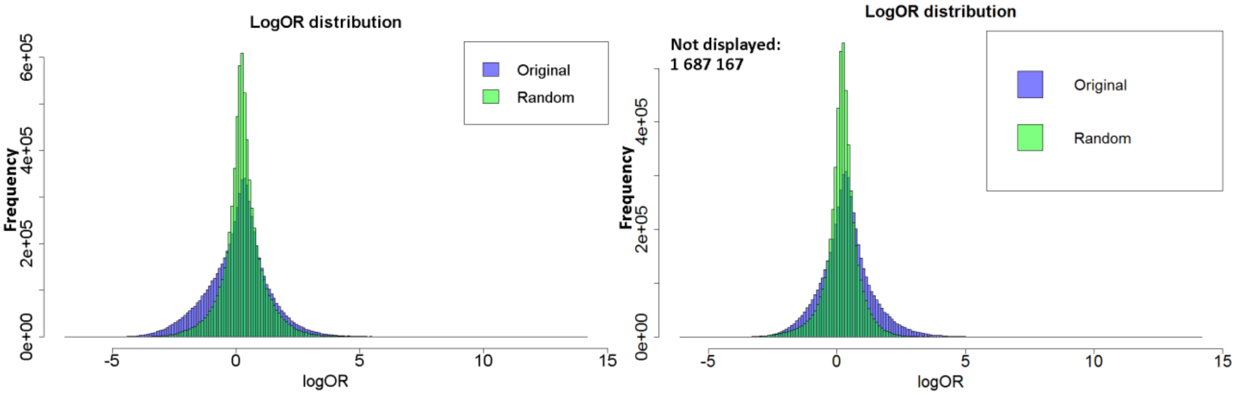

**Figure S13.** Distribution plots for all pairs of GO function and all pairs having significant contingency tables on Fungi organisms.

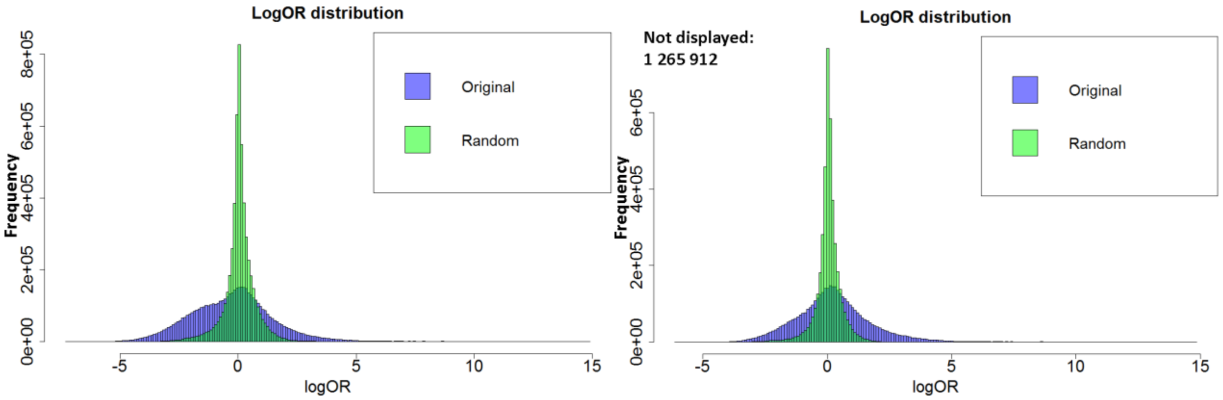

**Figure S14.** Distribution plots for all pairs of GO function and all pairs having significant contingency tables on Metazoa organisms.

#### Empirical assessment of association strength of enrichments

To empirically assess if the strength of association is significantly higher in original dataset than in randomized data (gene locations are permuted in a genome), we compute the log odds ratio on randomized data for each significantly enriched pair on the original data and assess the number of pairs with higher $log_2(OR)$ than the corresponding pair computed on randomized genome and the number of pairs with two-fold increase in $log_2(OR)$ compared to the corresponding pair computed on randomized genome.

This criterion is used, since it is not computationally feasible to calculate all log odds ratios on different randomized versions of a genome the number of times required to assess empirical p-values.

The number of pairs with higher or significantly higher $log_2(OR)$ than obtained by the corresponding pair on randomized genome for different scenarios and FDR thresholds are presented in Table S3.

Table S3. Percentage of significant pairs with a given criteria with larger or significantly larger log odds ratio than corresponding pair computed on randomized data.

| **Criteria** | #GOx-GOy pairs | %pairs $log_2(OR_{orig}) > log_2(OR_{rand})$ | %pairs $log_2(OR_{orig}) > 2 \cdot log_2(OR_{rand})$ | Organisms |
|---|---|---|---|---|
| GOx-GOx, $OR_x$>1 at FDR<20%; Z-test for significance of log odds ratio | 854/1048 | 100% | 99.2% | Prokaryot |
| | 918/2617 | 97.9% | 80% | Fungi |
| | 2316/2316 | 100% | 99.8% | Metazoa |
| GOx-GOy, $OR_x$>1 at FDR<20%; Z-test for significance of log odds ratio | $2.9x10^5$/ $1.1x10^6$ | 98.7% | 94.3% | Prokaryot |
| | $1.0x10^6$/ $6.8x10^6$ | 94.2% | 62.2% | Fungi |
| | $1.6x10^6$/ $5.4x10^6$ | 99.8% | 95.7% | Metazoa |
| GOx-GOy, $OR_x$>1 at FDR<10%; Z-test for significance of log odds ratio and Resnik Similarity < 1 | $4.6x10^4$/ $1.8x10^5$ | 98.7% | 94.1% | Prokaryot |
| GOx-GOy, $OR_x$>1 at FDR<10%; Z-test for significance of log odds ratio and Resnik Similarity < 2 | $3.1x10^5$/ $1.8x10^6$ | 93.1% | 58.7% | Fungi |
| | $3.8x10^5$/ $1.3x10^6$ | 99.8% | 95.4% | Metazoa |
| GOx-GOy, $OR_x$>2 at FDR<10%; Z-test for significance of log odds ratio and Resnik Similarity < 1 | 16356/177843 | 100% | 99.85% | Prokaryot |

| | | | | |
|---|---|---|---|---|
| GOx-GOy, $OR_x$>2 at FDR<10%; Z-test for significance of log odds ratio and Resnik Similarity < 2 | 69466/331277 | 99.7% | 85.6% | Fungi |
| | 226876/467656 | 99.96% | 96.8% | Metazoa |
| GOx-GOx or GOx-GOy with Resnik Similarity ≥6. $OR_x$>1 and J(GOx,GOy)≥0.6 at FDR<1%; Z-test for significance of log odds ratio | 6615 | 99.8% | 97.8% | Prokaryot |
| | 8853 | 97.3% | 74.4% | Fungi |
| | 36122 | 99.9% | 97.6% | Metazoa |
| GOx-GOy with Resnik Similarity <1 and J(GOx,GOy)≤ 0.05. $OR_x$>1 and FDR<1%; Z-test for significance of log odds ratio | 204202 | 98.5% | 94.1% | Prokaryot |
| GOx-GOy with Resnik Similarity <2 and J(GOx,GOy)≤ 0.05. $OR_x$>1 and FDR<1%; Z-test for significance of log odds ratio | 878850 | 94% | 61.4% | Fungi |
| | 1256151 | 98.8% | 95.5% | Metazoa |

#### Correlation between Jaccard index and Log odds ratio of pairs of functions

In this section, we demonstrate that there is only a partial correlation between the Jaccard index and the Log Odds Ratio for significantly enriched GO pairs of functions (FDR<0.1).
92% of functions presented in the plot have Jaccard index smaller than 0.1, but still have significant enrichment. This indicates that the effect of semantically distant, enriched functions is relevant and frequently occurring.

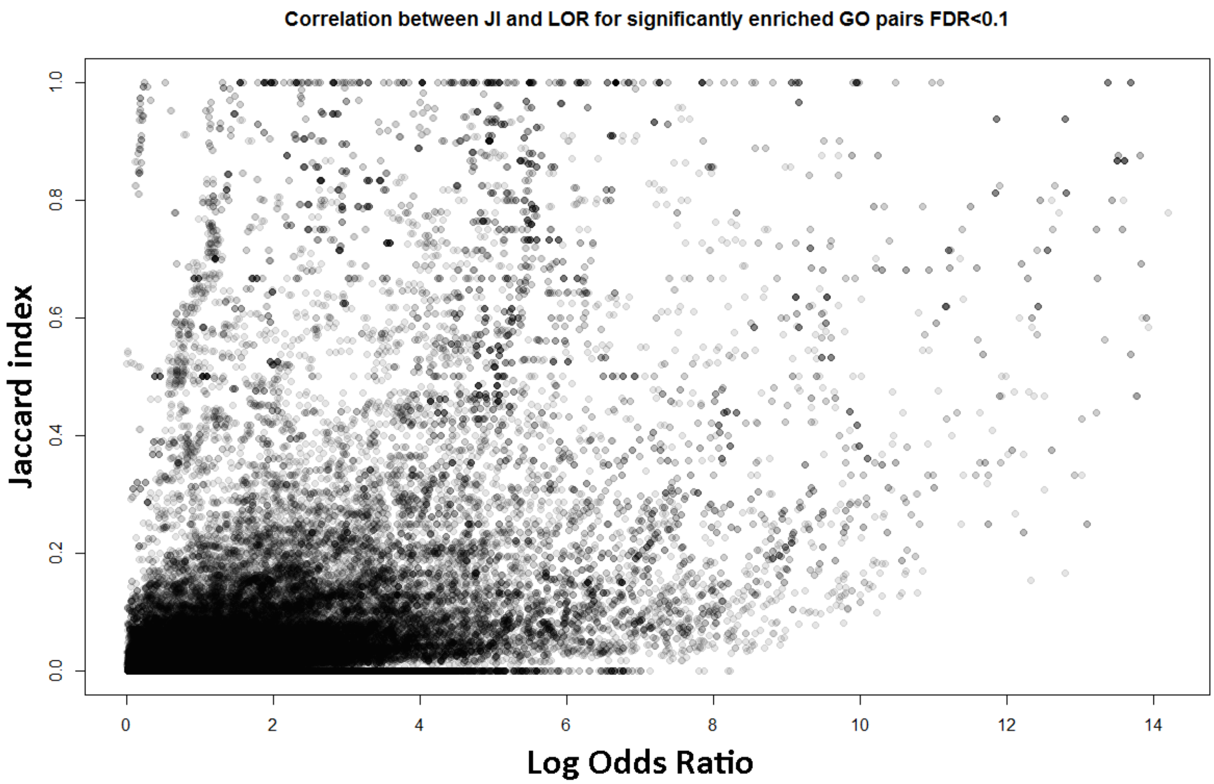

**Figure S15.** Correlation between Log Odds Ratio and Jaccard index for all significantly enriched GO pairs of functions with FDR<0.1.

#### S3.2 Evaluation of prediction accuracy of Neighborhood function profiles on prokaryotic genomes

In the continuation we present the comparative histograms showing the number of GO functions with a AUC or AUPRC (Figure S16) in the predefined interval for the 10-NN (red) and Neighborhood function profiles (green) methods. The frequency distribution is significantly shifted to the right for the NFP approach in all figures for AUC and AUPRC measures demonstrating significant improvement in accuracy of gene function prediction regardless generality of GO function. GO functions were divided to specific functions (information content ≥ 4) and general (information content <4).

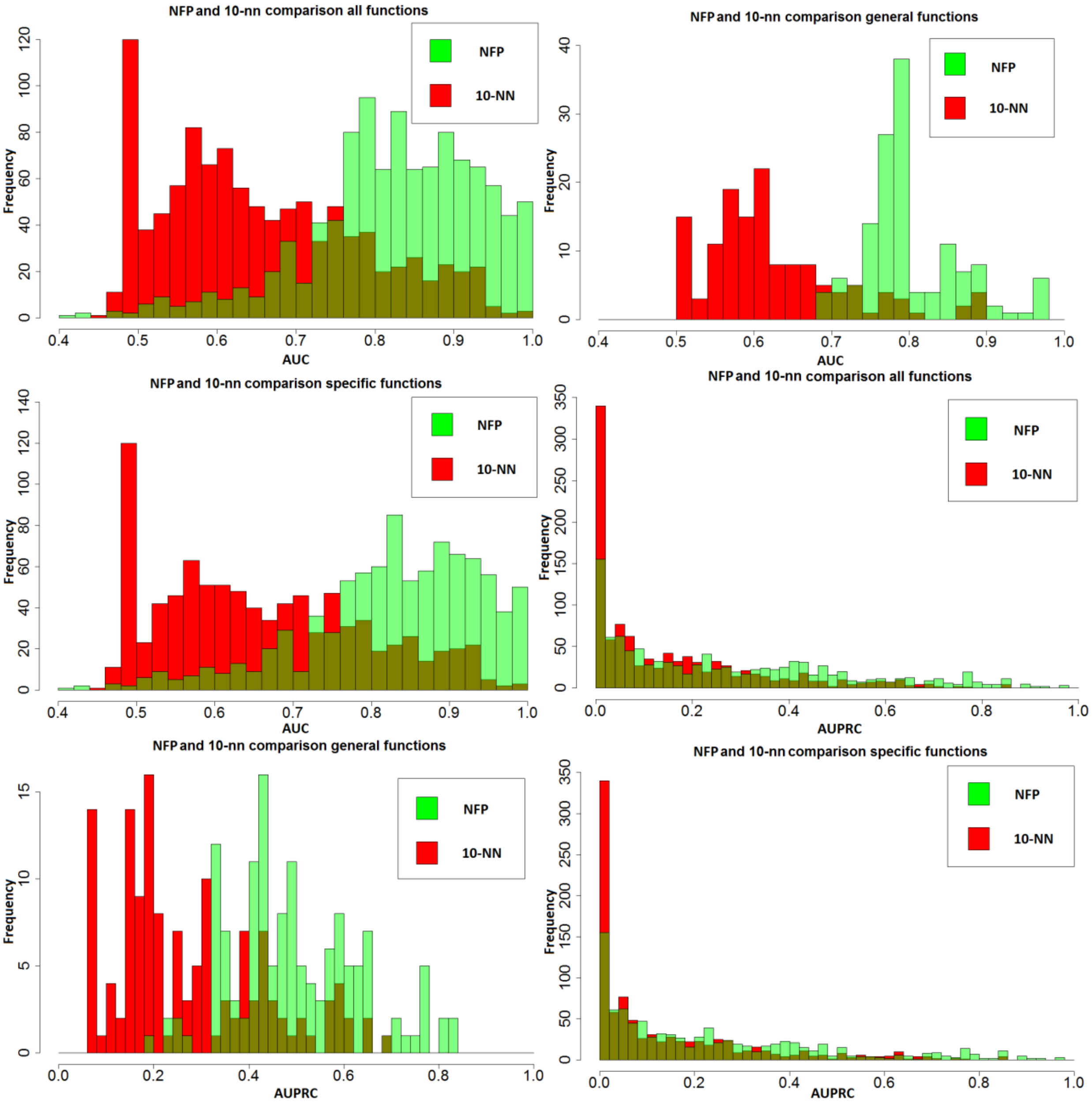

**Figure S16.** Comparative histograms showing number of GO functions with a given AUC (first three subfigures) and the number of GO functions with a given AUPRC (second three subfigures) for Neighborhood function profiles (green) and 10-NN (red) approach.

The comparative results for 1-NN, 3-NN, 10-NN, Gaussian Field Propagation [Mostafavi] and Neighborhood function profiles approach can be seen in Figure S17.

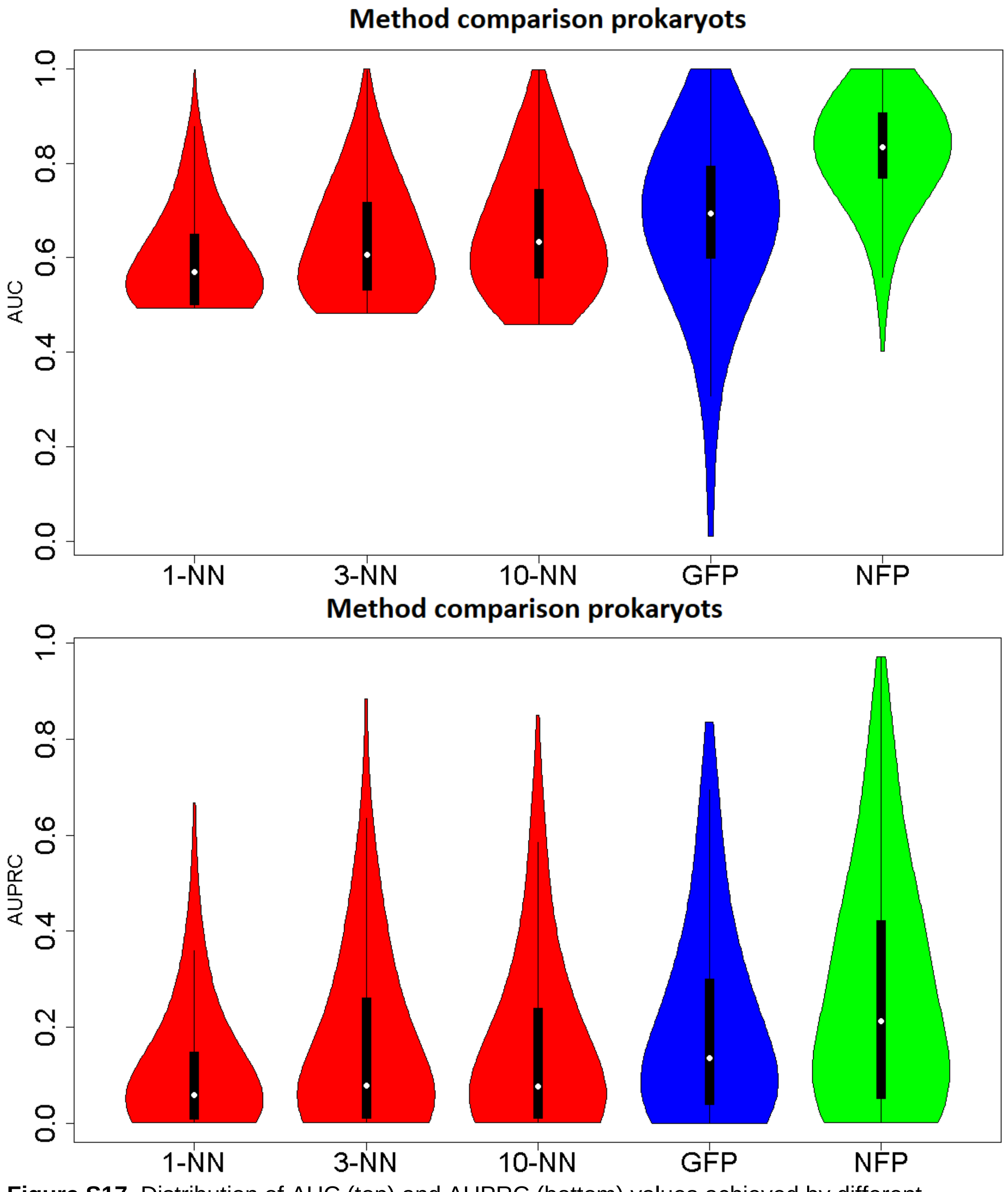

**Figure S17.** Distribution of AUC (top) and AUPRC (bottom) values achieved by different approaches on prokaryotic dataset.

We compare the AUC and AUPRC for the same set of functions using Bubble plots [Vidulin] for 1-NN, 3-NN, 10-NN, Gaussian Field Propagation [Mostafavi] and Neighborhood function profiles methods. Each bubble represents one GO term and the size of a bubble denotes GO generality. More general terms (with higher frequency) are denoted with larger bubbles whereas more specific terms with smaller bubbles. GO terms with frequency larger than 0.3*maxGenerality (very general terms) are not shown to make figures more visible. Boxplots represent

AUC/AUPRC distribution for general terms (frequency >0.2), medium general terms (frequency in [0.1,0.2>) and specific terms (frequency <0.1). GO terms are further divided by different ontologies to Biological process, Cellular component and Molecular function.

Figure S18 demonstrates that Neighborhood function profiles methodology significantly outperforms other methods with respect to AUC score on all three ontology namespaces and on GO terms contained in all three generality levels. Neighborhood function profiles methodology also outperforms all methods with respect to AUPRC score on Biological process namespace on all generality levels, whereas it has slightly worse accuracy then Gaussian Field Propagation method on the specific functions on the Cellular component namespace (however, this ontology contains a very small number of functions making results highly unstable). The Gene Functional Neighborhood methodology outperforms other methods on medium general and general categories.

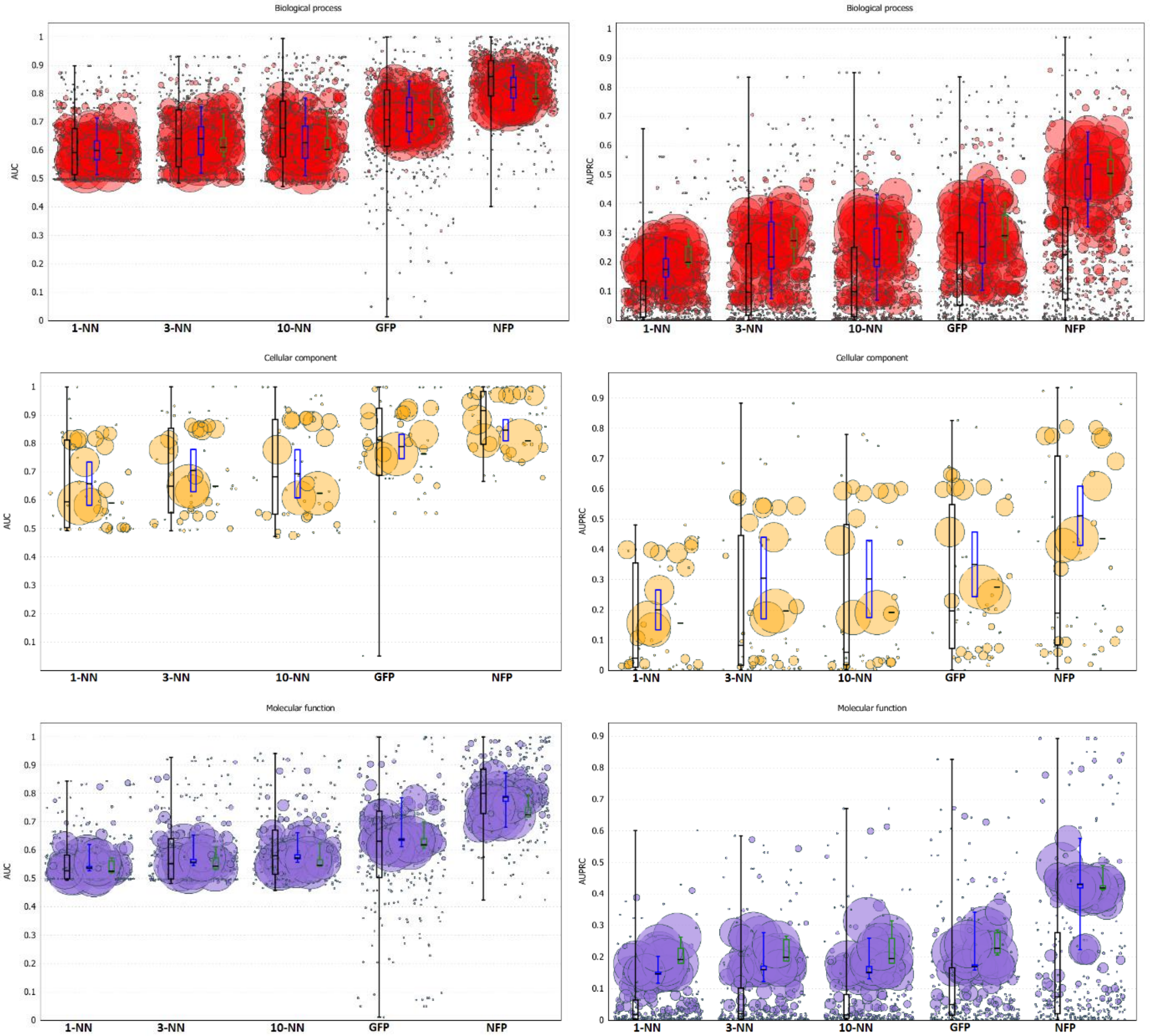

**Figure S18.** Accuracy comparison of 1-NN, 3-NN, 10-NN, Gaussian Field Propagation and Neighborhood function profiles methodologies for gene function prediction. Crossvalidation AUC (left column) and AUPRC (right column) scores are shown for functions of different generality (bubble size) and GO sub-ontologies (rows).

#### S3.3 Evaluation of prediction accuracy of Neighborhood function profiles on fungal genomes

Comparison with baseline methods (Gaussian Field Propagation and k-NN approach) was performed on 49 Fungi genomes obtained from the Eggnog database [Cepas]. Histogram comparing number of GO functions with AUC/AUPRC in a predefined interval achieved by Neighborhood function profiles and 10-NN method are available in Figure S19.

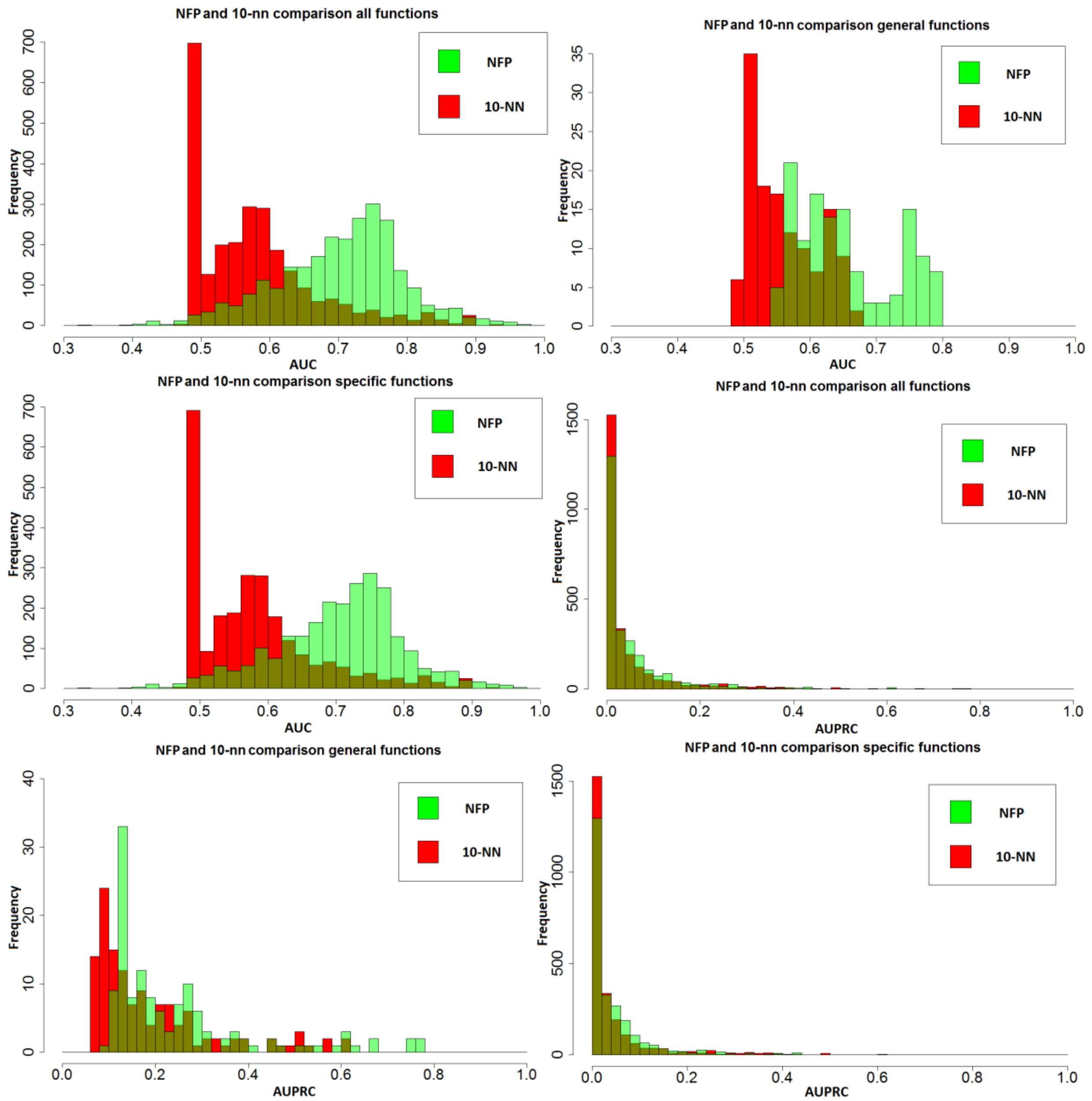

**Figure S19.** Comparative histograms showing number of GO functions with a given AUC (first three subfigures) and the number of GO functions with a given AUPRC (second three subfigures) for Neighborhood function profiles (green) and 10-NN (red) approach on the Fungi genomes.

Figure S19 demonstrates that Neighborhood function profiles methodology has significantly shifted (higher) values of AUC score on all groups of tested function, strongly shifter values of AUPRC score on general function and visibly shifted score on all and specific functions. The distribution of AUC/AUPRC scores for different approaches on fungi dataset can be seen in Figure S20.

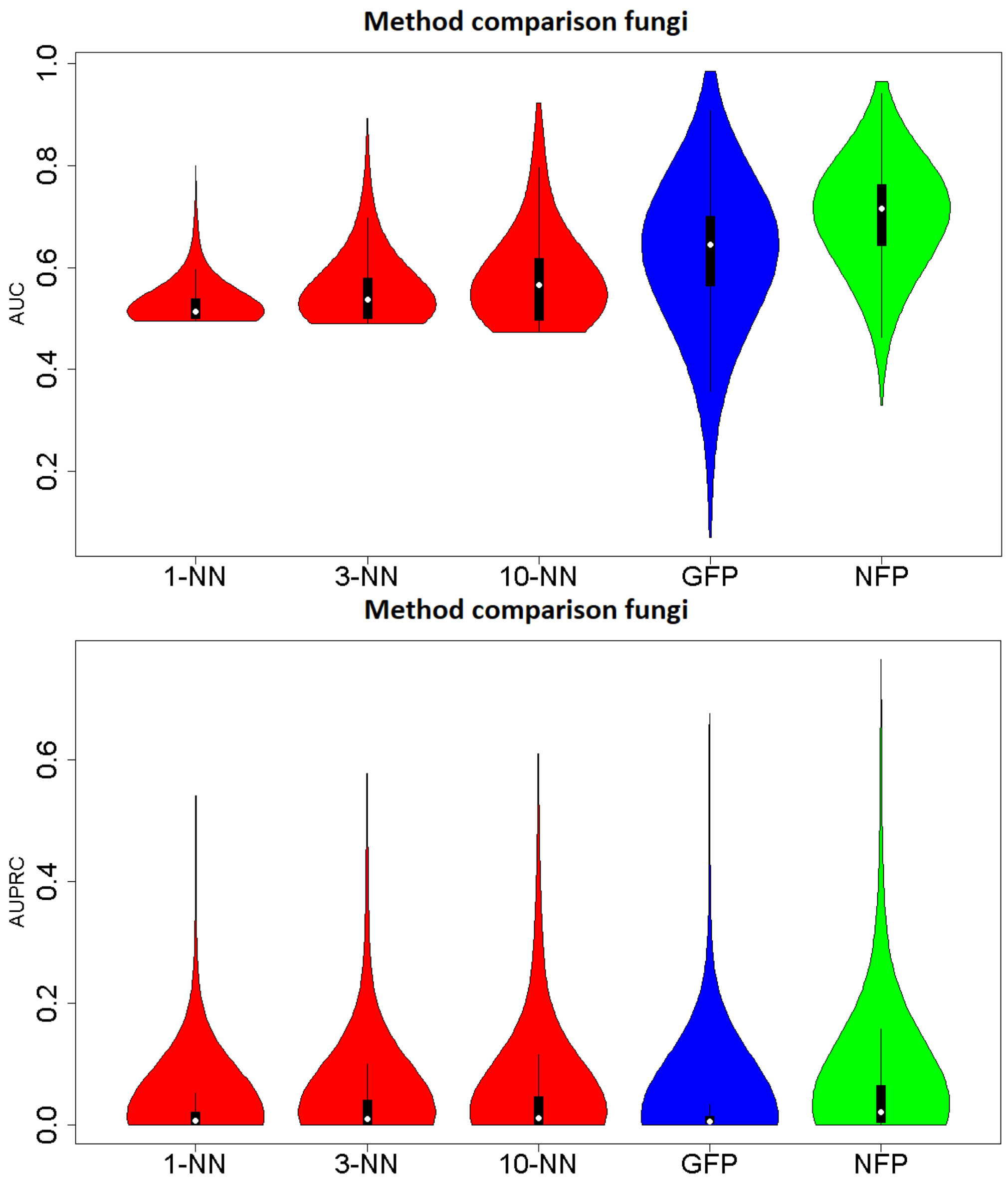

**Figure S20.** Distribution of AUC (top) and AUPRC (bottom) scores achieved by different approaches on the Fungi dataset.

It can be seen in Figure S20 that neighborhood function profiles method outperforms baseline methods on all ontologies with respect to AUC and AUPRC measures for categories contained in all three defined generality classes.  Figure S21 shows that Neighborhood function profiles outperform other approaches on Fungi organisms on both the AUC and AUPRC measures for all namespaces of GO ontologies and GO generality levels.

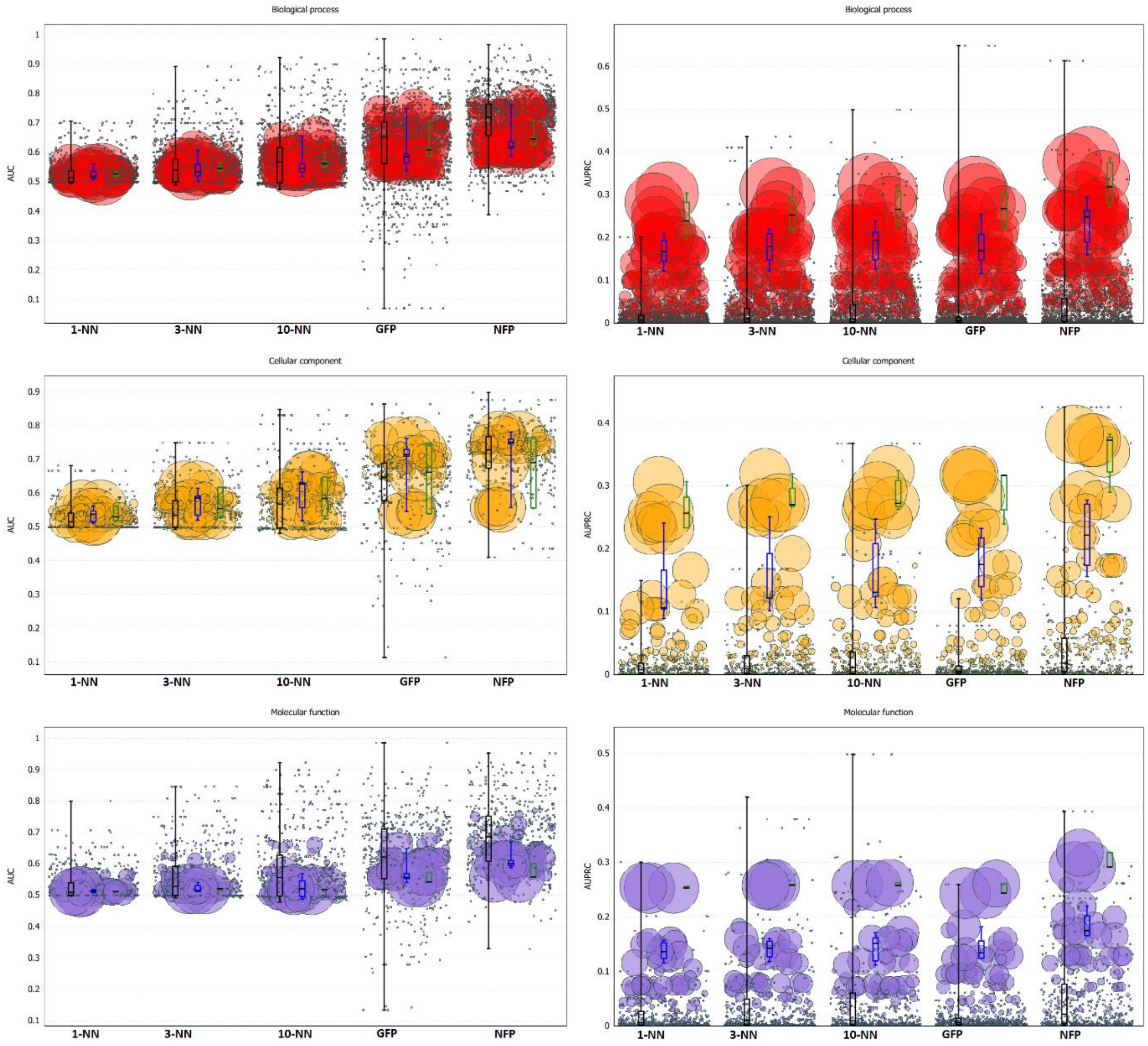

**Figure S21.** AUC (left) and AUPRC (right) distribution for GO categories of different generality (bubble size) belonging to different GO sub-ontologies (rows) for Neighborhood function profiles and Gaussian Field Propagation *versus* baseline methods on Fungi genomes.

#### S3.4 Evaluation of prediction accuracy of Neighborhood function profiles on metazoan genomes

We compare the Neighborhood function profiles with Gaussian Field Propagation and baseline k-NN methods on 80 Metazoa genomes.
Histogram comparing number of GO functions with AUC/AUPRC in a predefined interval achieved by Neighborhood function and 10-NN method are available in Figure S22.

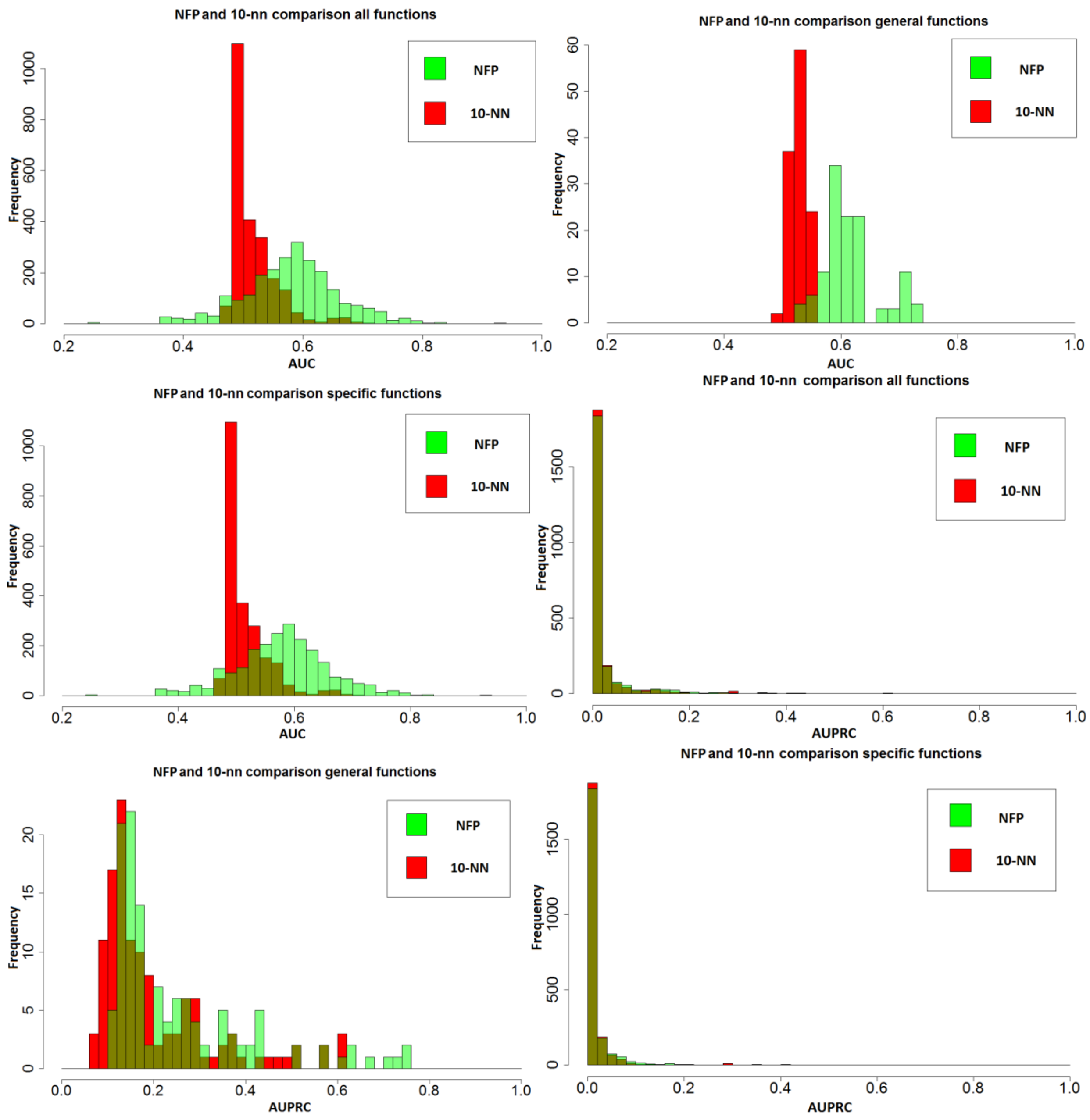

**Figure S22.** Comparative histograms showing number of GO functions with a given AUC (first three subfigures) and the number of GO functions with a given AUPRC (second three subfigures) for Neighborhood function profiles (green) and 10-NN (red) approach on the Metazoa genomes.

Comparison of 1-NN, 3-NN, 10-NN, GFP and NFP approaches on all functions used for gene function prediction on metazoa organisms can be seen in Figure S23.

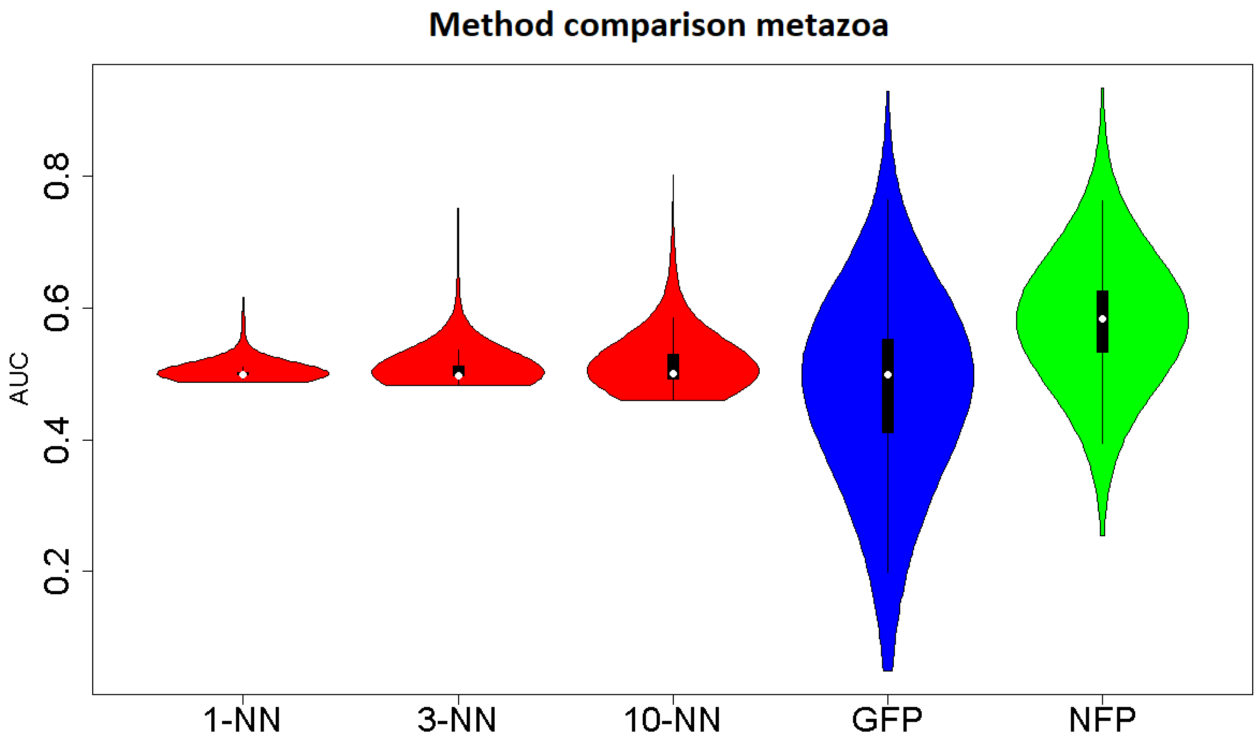

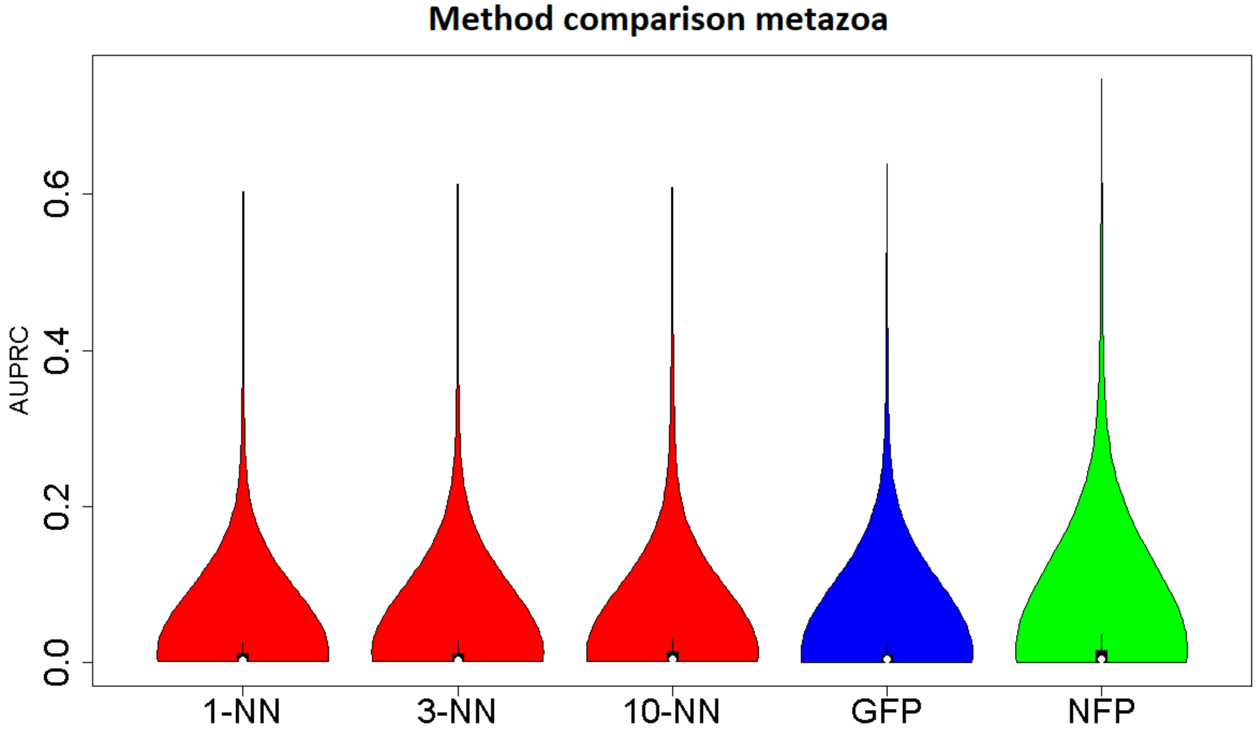

**Figure S23.** Method comparison based on AUC score(top) and AUPRC (bottom).

Method comparison, based on AUC/AUPRC score, divided in different namespaces of GO ontology is presented in Figure S24.

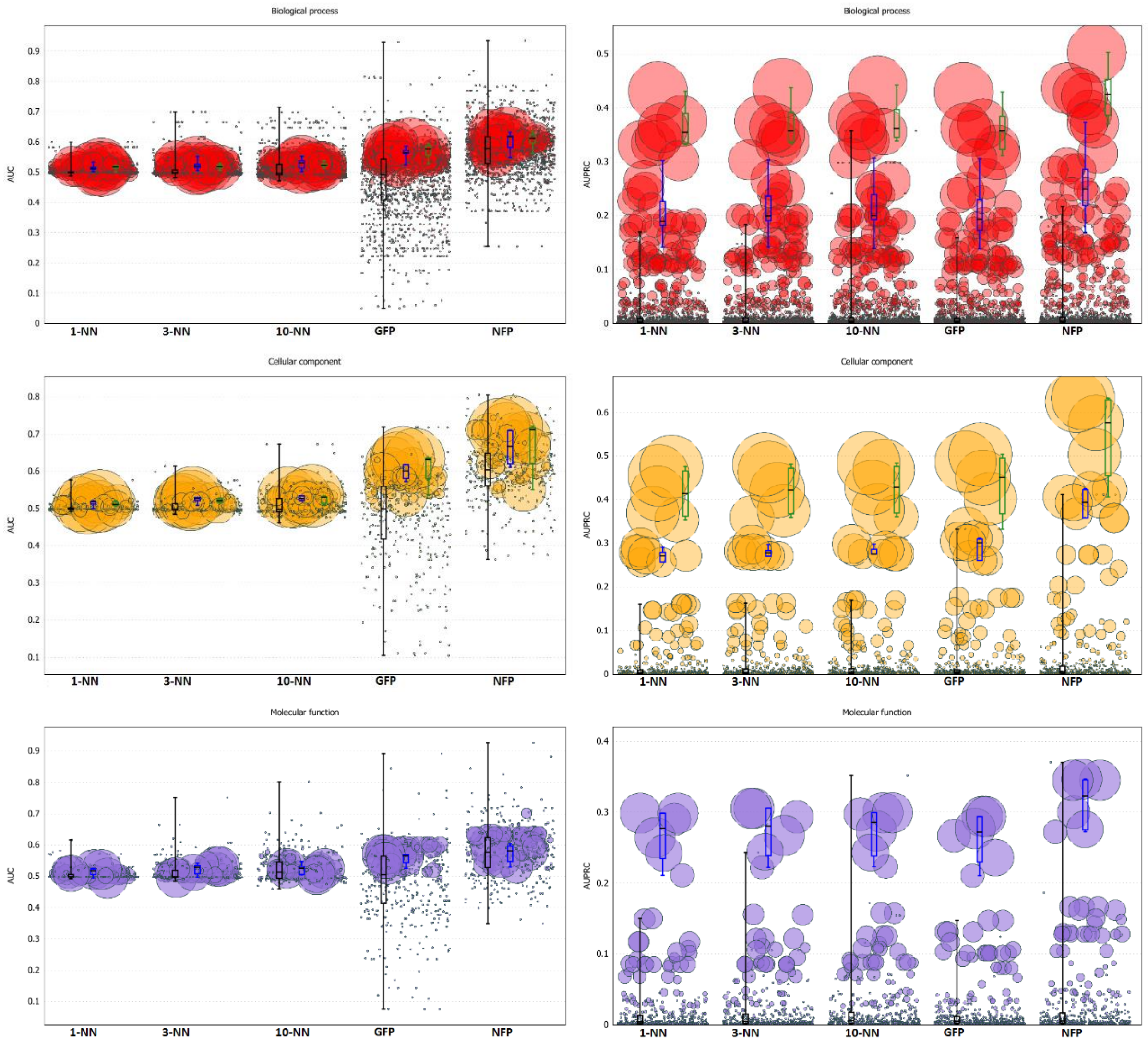

**Figure S24.** AUC (left column) and AUPRC (right column) distribution for accuracy of predicting GO terms with different generality (bubble size) and belonging to different GO sub-ontologies (rows) for the Neighborhod Function Profiles (NFP) *versus* baseline methods on Metazoa genomes.

It can be seen in Figure S24 that Neighborhood function profiles outperforms baseline methods on all ontologies with respect to AUC and AUPRC measures for categories contained in all three defined generality classes.

#### S3.5 Ru-Mi curves on prokaryotic dataset

Ru-Mi curves measure remaining uncertainty and an amount of misinformation obtained from predictions produced by a classification algorithm [Clark et al, 2013]. The *remaining uncertainty* corresponds to the information about the protein function, that is not provided by the prediction, i.e. graph *P,* in relation to the true subgraph of functions *T*. The remaining uncertainty (*ru*) is defined as:

$$ru(T,P) = \sum_{n \in T-P} ia(n)$$

This is simply the total information content of the nodes in *T* of the GO, but not in the *P.* The *misinformation* introduced by some prediction corresponds to the total information content of the nodes of GO in prediction subgraph *P*, which are not matching those in a true subgraph *T*.

$$mi(T,P) = \sum_{n \in P-T} ia(n)$$

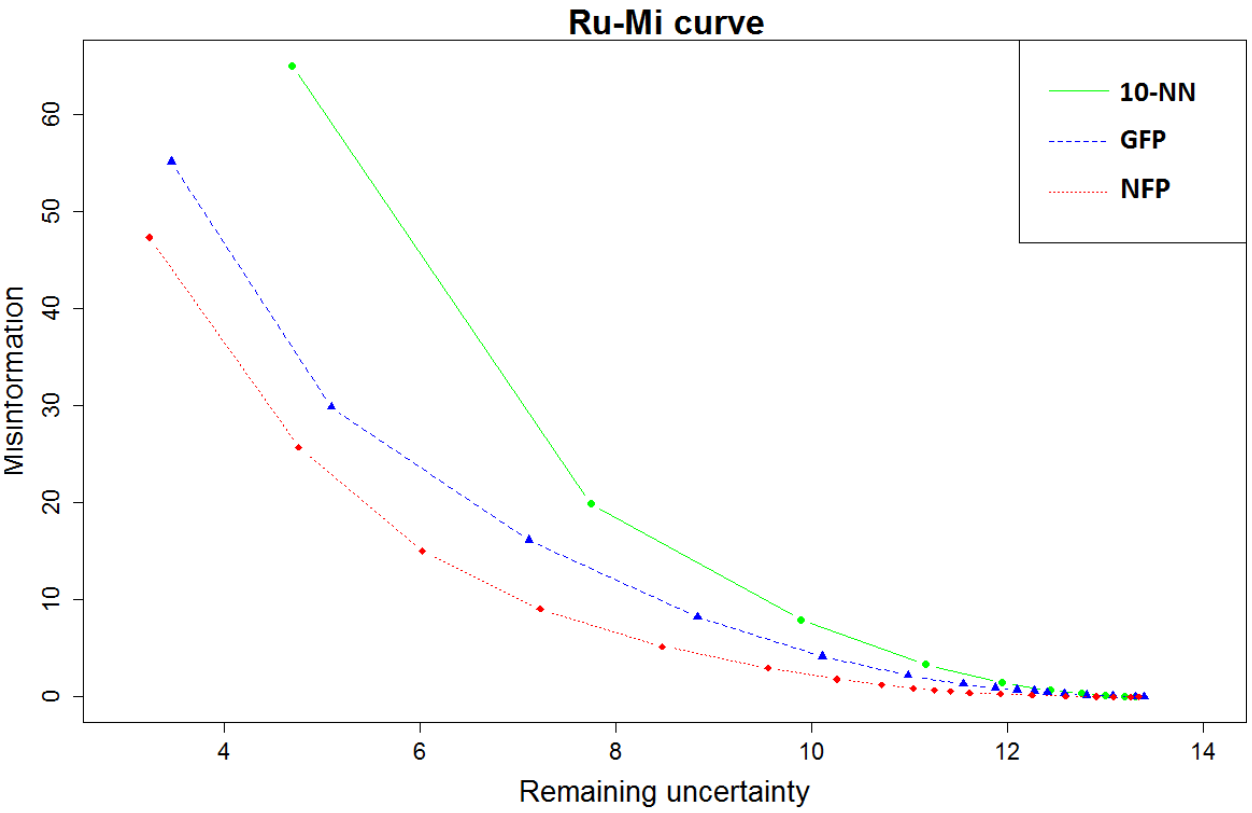

**Figure S25.** Ru-Mi curves for 10-NN, Gaussian Field Propagation (GFP) and Neighborhood Function Profiles (NFP) approach.

It can be seen in Figure S25 that the NFP approach exhibits the smallest amount of misinformation given a remaining uncertainty threshold among the tested approaches.

#### Ru-Mi curves on Eukaryotic datasets

We show Ru-Mi curves obtained in Fungi and Metazoa dataset in Figure S26. We can see that NFP approach outperforms other approaches on both datasets with respect to amount of misinformation given some uncertainty threshold.

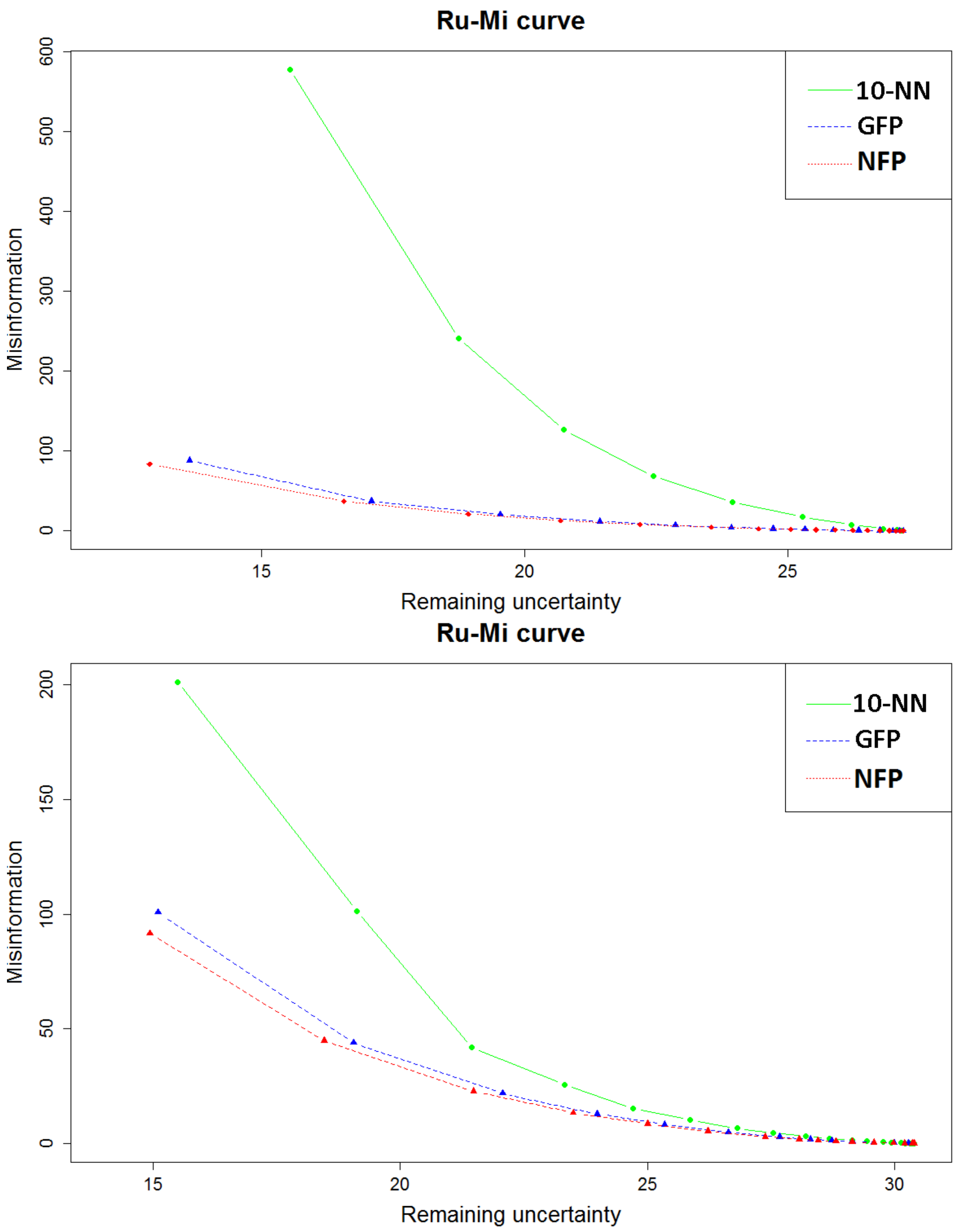

**Figure S26.** Ru-Mi curves for 10-NN, Gaussian Field Propagation and Neighborhood function profiles approach on Fungi dataset (top) and Metazoa dataset (bottom).

#### Classifier performance for different numbers of neighbours

a) prokaryotes

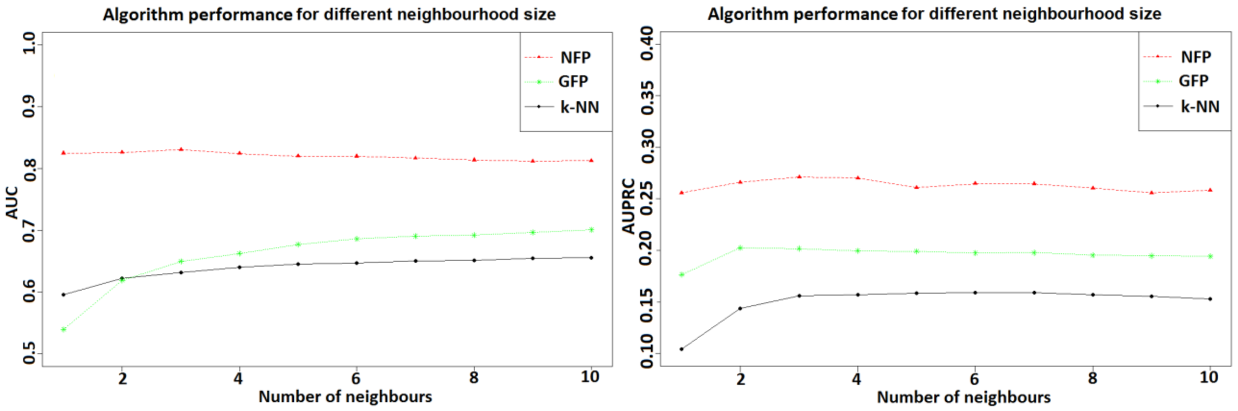

b) fungi

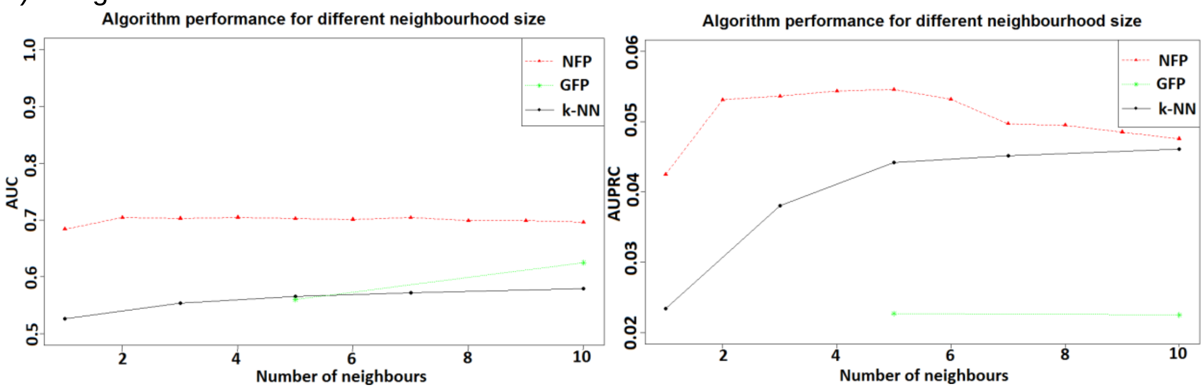

c) metazoa

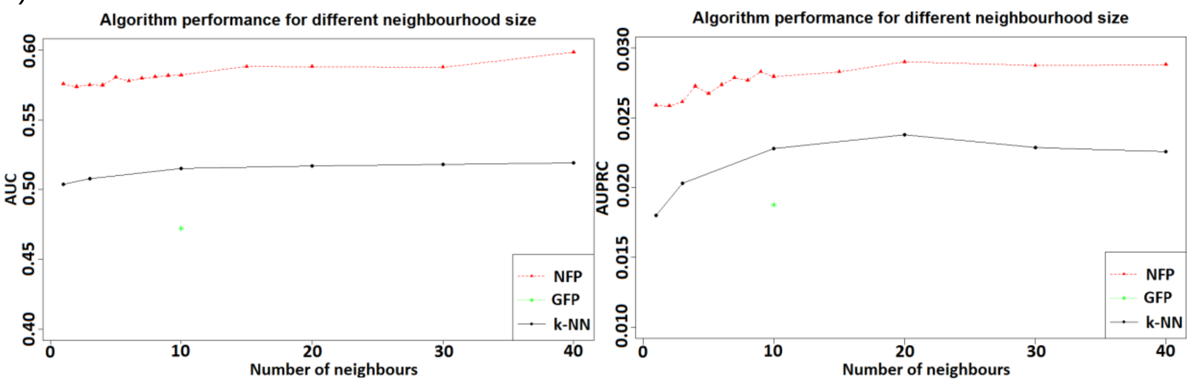

**Figure S27.** Average AUC (left) and AUPRC (right) on the dataset obtained from Prokaryotic organisms a) for Neighborhood function profiles (NFP), Gaussian Field Propagation (GFP) and k-NN approach.

Figure S27 a) demonstrates that NFP approach reaches maximal average AUPRC for k=3 (using 3 neighbouring genes to compute functional Neighborhoods). k-NN approach average AUPRC rises until k=7 and then slowly declines. AUC generally shows similar behaviour for NFP, however it increases constantly for the k-NN approach.

Since k-NN approach searches for OGs that are the nearest neighbours across all genomes (globally), increasing parameter k, leads to more stable predictions since many similar neighbours reinforce the prediction of conserved function. Neighborhood function profiles approach works on a different principle, looking at functional Neighborhoods in local (physical) environment of a gene. Increasing the parameter k (up to k=3) increases performance, however further increase slowly decreases AUPRC score. This is the consequence of increased noise when adding genes that are far apart in the genome (more than 3 genes apart) from a central gene. Such genes may contain increasing number of functions unrelated to the central gene.

The GFP approach has decreasing AUPRC score from k=2 onwards, however it has increasing AUC score.  Given the presented results, we present all detailed result analyses for k=5 (since it shows good trade-off in performance for both AUC and AUPRC score).

Genomes of eukaryotic organisms are bigger and different in structure (linear) compared to circular prokaryotic genomes. Here we use k=10 for the k-NN and the GFP algorithms[7] and k=5 for the NFP algorithm.  In Fungi organisms, the chosen parameters yield the best performance. The k-NN approach reaches the peak performance at k=20 for Metazoa organisms, although the difference compared to k=10 is very small. NFP approach has slightly increasing performance with the increase of parameter k. It has a peek at k=5 with respect to AUC score (see Figure S27 b) and c)).

#### Classifier performance for selected functions having high enrichment with at least one dissimilar function

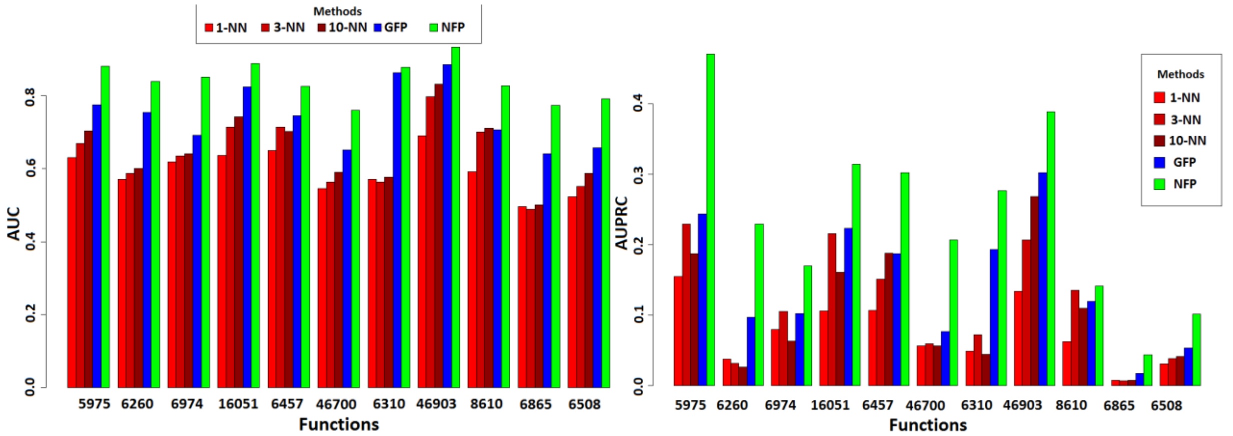

**Figure S28.** AUC (left) and AUPRC (right) for 11 selected functions predicted by 1-NN, 3-NN and 10-NN algorithms (red), Gaussian Field Propagation (blue) and Neighborhood function profiles (green) method.

[7] GFP approach has a very long execution time on eukaryotic datasets, making Greedy search approach for parameter testing infeasible. It has been determined that k=10 has better performance than k=5. For k-NN approach k=10 yields favorable performance (k=20 yields only slightly better performance on Metazoa organisms).

Figure S28 shows that Neighborhood function profiles and Gaussian Field Propagation methods significantly outperform baseline methods (k-NN) on selected functions (functions having highly enriched semantically distant functions). Moreover, NFP significantly outperform Gaussian Field Propagation method in both AUC and AUPRC measures. This figure demonstrates that NFP can use enrichments even though they are present between semantically dissimilar functions and uses this information to improve gene function prediction.

#### S3.6 Predicting gene function from genomic neighborhoods can be greatly improved by taking semantically dissimilar functions into account

In this section, we study the effects of using semantically dissimilar functions in gene function prediction, using Gene Functional Neighborhood approach. To do this, we perform 200 runs of cross-validation for each GO category using single class prediction with Fast Random Forest algorithm[8]. In these experiments, we use only GO categories from the Biological Process namespace. We test classifier performance using different subsets of features: a) All features from Biological Process namespace, b) All features corresponding to the occurence of semantically close functions to the target GO in the Neighborhood of a target GO, c) All features corresponding to the occurence of semantically close functions to the target GO in the Neighborhood of a target GO but not containing target GO and its parents, d) All features corresponding to the occurence of semantically close functions to the target GO in the Neighborhood of a target GO but not containing target GO, its parents and highly enriched GO terms to the target GO, e) All features corresponding to the occurence of medium distant functions to the target GO in the Neighborhood of a target GO, f) All features corresponding to the occurence of medium distant functions to the target GO in the Neighborhood of a target GO not containing parents of the target GO, g) All features corresponding to the occurence of medium distant functions to the target GO in the Neighborhood of a target GO not containing parents of the target GO or any enriched GO function occurring in the Neighborhood of target GO, h) All features corresponding to the occurrence of semantically distant functions to the target GO in the Neighborhood of a target GO, i)  All features corresponding to the occurrence of semantically distant functions to the target GO in the Neighborhood of a target GO  not containing parents of the target GO, j) All features corresponding to the occurrence of semantically distant functions to the target GO in the Neighborhood of a target GO  not containing parents of the target GO or any enriched GO function in the Neighborhood of a target GO, k) All features corresponding to the occurrence of a target GO function and its parents in the Neighborhood of a target GO.

[8] https://github.com/GenomeDataScience/FastRandomForest

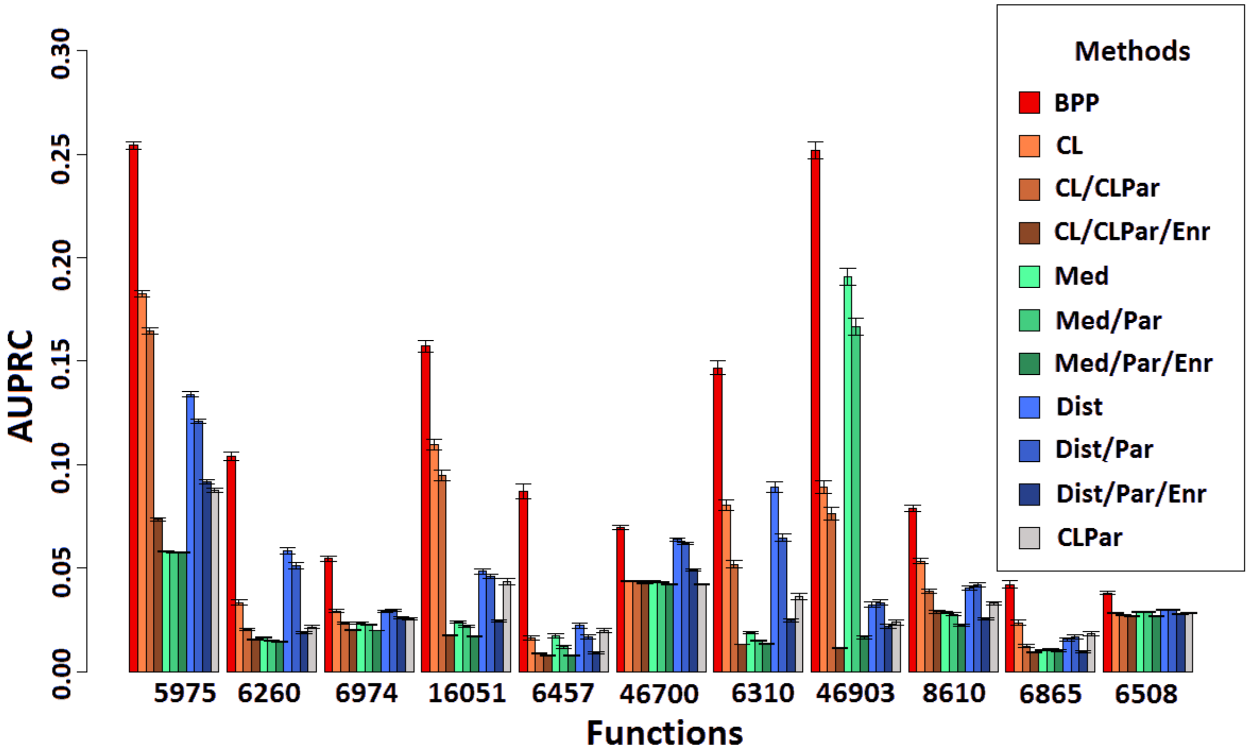

**Figure S29.** The effects of removing enriched functions from feature set for a selected subsets of features and comparison to CLPar set of features (containing the selected function and its parents, which closely mimics information obtainable by the guilt-by-association approaches). Barplots show the AUPRC value obtained by the Fast Random Forest method trained on a selected subset of features, whereas error bars show standard deviation among 200 runs of cross-validation.

We can see from Figure S29. that for the selected set of functions (these containing highly enriched dissimilar functions in the Neighborhood) occurence of highly enriched functions in the Neighborhood play important role in predictive accuracy of Neighborhood function profiles method. Moreover, for all but one function, the approach manages to achieve better performance learning from features representing occurrence of highly enriched semantically distant functions than from Neighborhoods representing occurrence of the target function and its parents.

#### The amount of new information obtained with proposed methodology

In this section, we measure the amount of information obtained from predictions of Neighborhood function profiles method (first diagram) by computing the information accretion [Clark]. We compare the amount of information obtained with other state-of-the art methods (Gaussian Field Propagation - second diagram and k-NN - third diagram).

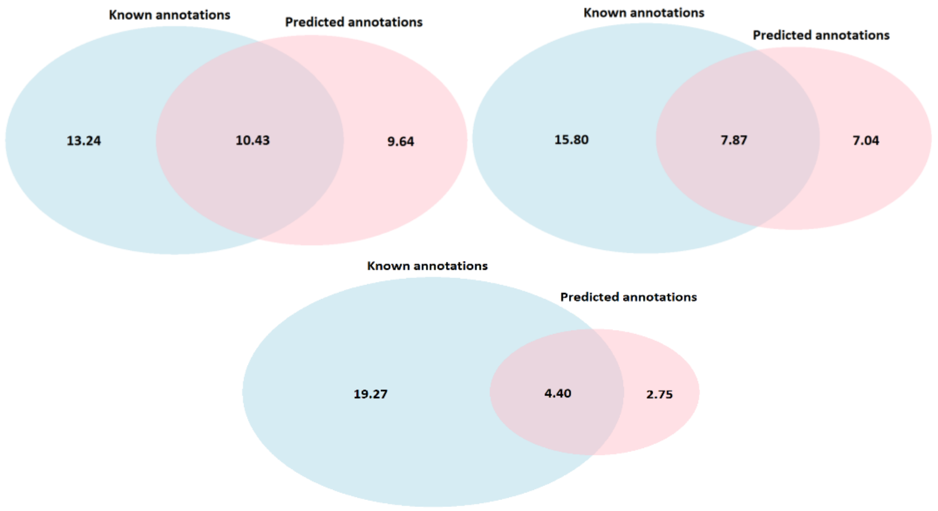

**Figure S30.** Average information accretion (bits per gene) obtained with Neighborhood function profiles approach (top-left), Gaussian Field Propagation (top-right) and 10-NN (bottom-center) on prokaryotes dataset. The diagram shows the average number of bits per gene contained in known annotations that were not obtained by the classifier with precision >=0.5, average number of bits per gene contained in known annotations obtained with the used methodology and the average number of bits per gene of newly obtained information (previously unknown).

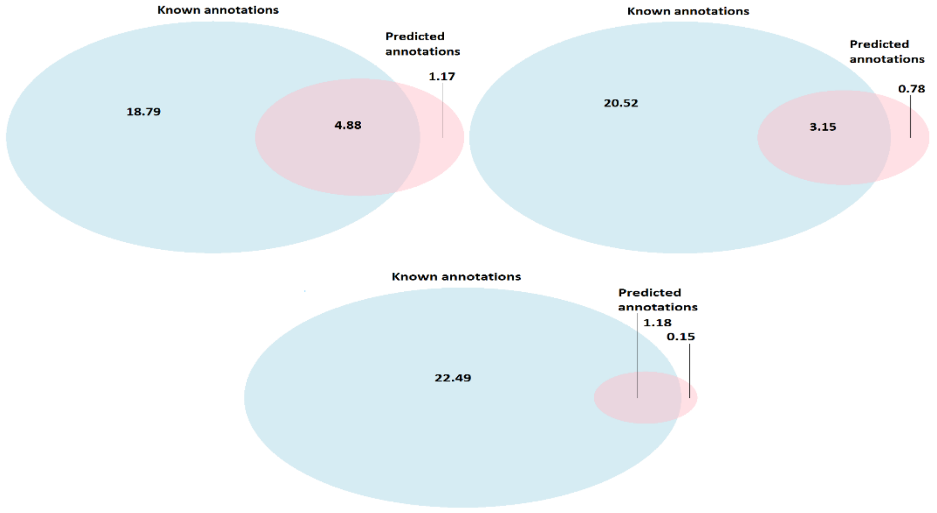

**Figure S31.** Average information accretion (bits per gene) obtained with Neighborhood function profiles approach (top-left), Gaussian Field Propagation (top-right) and 10-NN (bottom-center) on prokaryotes dataset. The diagram shows the average number of bits per gene contained in known annotations that were not obtained by the classifier with precision >=0.8, average number of bits per gene contained in known annotations obtained with the used methodology and the average number of bits per gene of newly obtained information (previously unknown).

On eukaryotic datasets all methodologies produce significantly smaller number of bits of information per gene, however it is evident that NFP approach significantly outperforms other tested approaches.

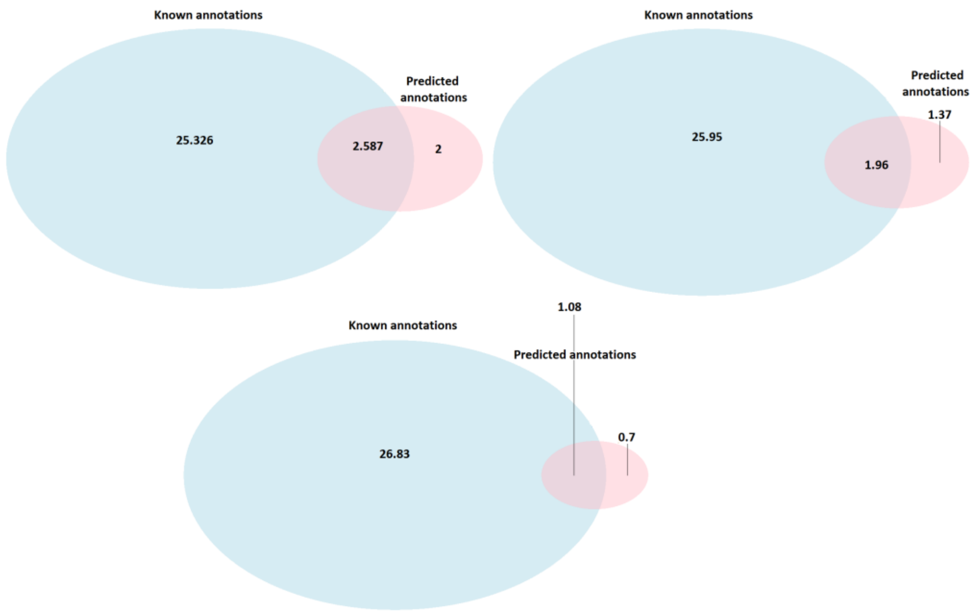

**Figure S32.** Average information accretion (bits per gene) obtained with Neighborhoods function profile approach (top-left), Gaussian Field Propagation (top-right) and 10-NN (bottom-center) on fungi dataset. The diagram shows the average number of bits per gene contained in known annotations that were not obtained by the classifier with precision >=0.5, average number of bits per gene contained in known annotations obtained with the used methodology and the average number of bits per gene of newly obtained information (previously unknown).

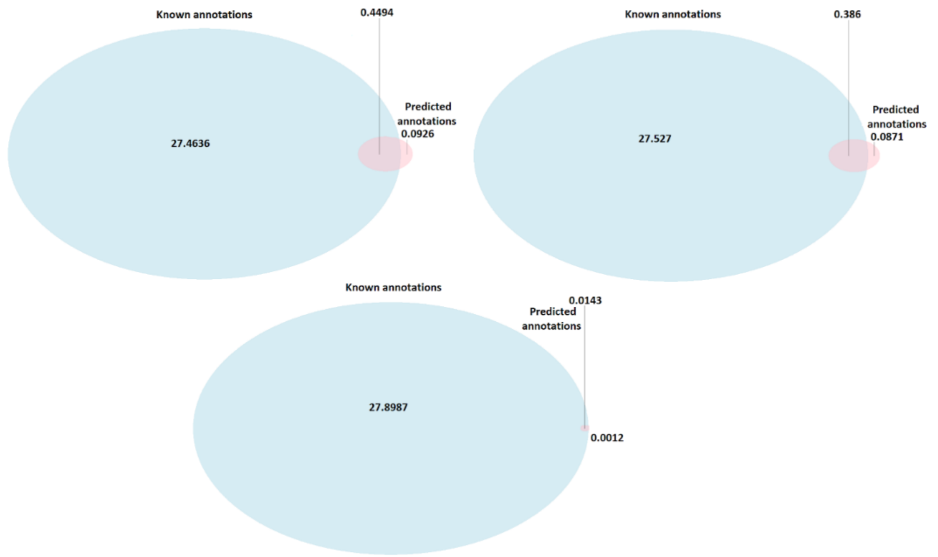

**Figure S33.** Average information accretion (bits per gene) obtained with Neighborhood function profiles approach (top-left), Gaussian Field Propagation (top-right) and 10-NN (bottom-center) on fungi dataset. The diagram shows the average number of bits per gene contained in known annotations that were not obtained by the classifier with precision >=0.8, average number of bits per gene contained in known annotations obtained with the used methodology and the average number of bits per gene of newly obtained information (previously unknown).

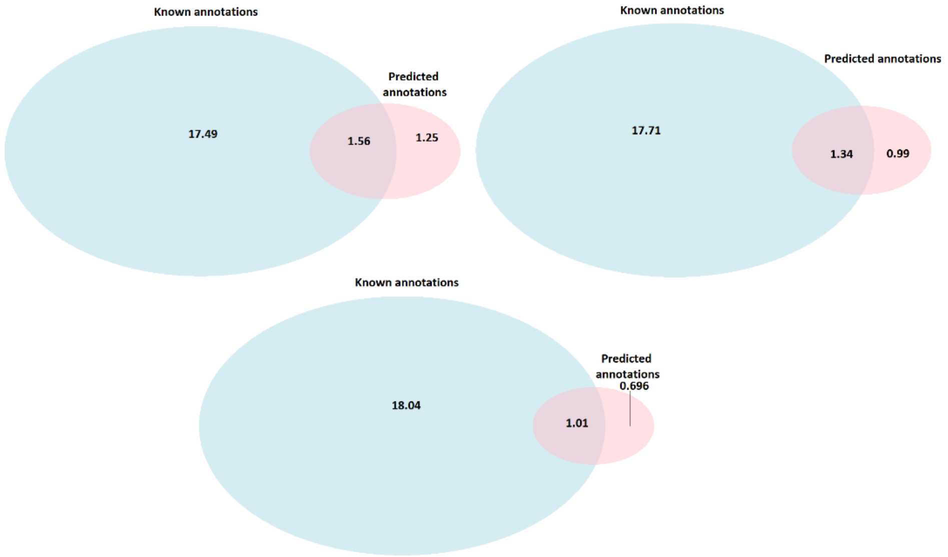

**Figure S34.** Average information accretion (bits per gene) obtained with Neighborhood function profiles approach (top-left), Gaussian Field Propagation (top-right) and 10-NN (bottom-center) on metazoa dataset. The diagram shows the average number of bits per gene contained in known annotations that were not obtained by the classifier with precision >=0.5, average number of bits per gene contained in known annotations obtained with the used methodology and the average number of bits per gene of newly obtained information (previously unknown).

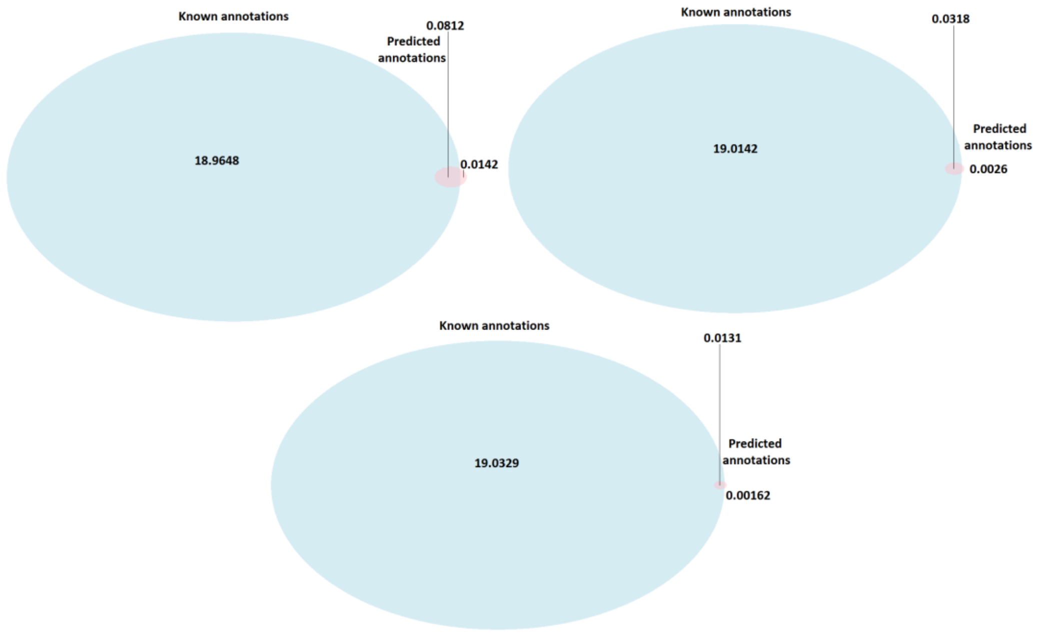

**Figure S35.** Average information accretion (bits per gene) obtained with Neighborhood function profiles approach (top-left), Gaussian Field Propagation (top-right) and 10-NN (bottom-center) on metazoa dataset. The diagram shows the average number of bits per gene contained in known annotations that were not obtained by the classifier with precision >=0.8, average number of bits  per gene contained in known annotations obtained with the used methodology and the average number of bits per gene of newly obtained information (previously unknown).

**Table S4. Information accretion (bits per gene) obrtained with Neighbourhood function profile (NFP), Gaussian Field Propagation (GFP) and 10-NN approach on prokaryotic, fungi and metazoa dataset. The results are divided by different ontology namespaces (Biological profile – BP, Molecular function – MF and Cellular component – CC).**

| Organisms | Precision | Ontology namespace | Method | Known | Known and predicted | Newly predicted |
|---|---|---|---|---|---|---|
| Prokaryots | 0.5 | BP | 10-NN | 11.77 | 2.81 | 1.74 |
| | | | GFP | | 4.90 | 4.31 |
| | | | NFP | | **6.11** | **5.53** |
| | | MF | 10-NN | 9.62 | 1.16 | 0.65 |
| | | | GFP | | 2.00 | 1.67 |
| | | | NFP | | **3.12** | **2.99** |
| | | CC | 10-NN | 2.28 | 0.43 | 0.36 |
| | | | GFP | | 0.96 | 1.05 |
| | | | NFP | | **1.20** | **1.12** |
| | 0.8 | BP | 10-NN | 11.77 | 0.69 | 0.07 |
| | | | GFP | | 1.97 | 0.51 |
| | | | NFP | | **3.13** | **0.75** |
| | | MF | 10-NN | 9.62 | 0.24 | 0.03 |
| | | | GFP | | 0.77 | 0.16 |
| | | | NFP | | **1.21** | **0.28** |
| | | CC | 10-NN | 2.28 | 0.25 | 0.05 |
| | | | GFP | | 0.41 | 0.11 |
| | | | NFP | | **0.54** | **0.13** |

| | | | | | | |
|---|---|---|---|---|---|---|
| Fungi | 0.5 | BP | 10-NN | 13.63 | 0.40 | 0.32 |
| | | | GFP | | 0.49 | 0.37 |
| | | | NFP | | **0.78** | **0.68** |
| | | MF | 10-NN | 7.24 | 0.08 | 0.08 |
| | | | GFP | | 0.19 | 0.18 |
| | | | NFP | | **0.44** | **0.43** |
| | | CC | 10-NN | 7.04 | 0.60 | 0.31 |
| | | | GFP | | 1.29 | 0.82 |
| | | | NFP | | **1.36** | **0.90** |
| | 0.8 | BP | 10-NN | 13.63 | 0.01 | 0.000 |
| | | | GFP | | 0.00 | 0.000 |
| | | | NFP | | **0.02** | **0.003** |
| | | MF | 10-NN | 7.24 | 0.006 | 0.000 |
| | | | GFP | | 0.000 | 0.000 |
| | | | NFP | | **0.009** | **0.001** |
| | | CC | 10-NN | 7.04 | 0.01 | 0.000 |
| | | | GFP | | 0.39 | 0.087 |
| | | | NFP | | **0.42** | **0.088** |
| Metazoa | 0.5 | BP | 10-NN | 8.28 | 0.13 | 0.12 |
| | | | GFP | | 0.23 | 0.21 |
| | | | NFP | | **0.33** | **0.32** |
| | | MF | 10-NN | 5.65 | 0.21 | 0.15 |
| | | | GFP | | **0.25** | 0.17 |
| | | | NFP | | **0.25** | **0.19** |
| | | CC | 10-NN | 5.12 | 0.66 | 0.42 |
| | | | GFP | | 0.85 | 0.61 |
| | | | NFP | | **0.97** | **0.74** |
| | 0.8 | BP | 10-NN | 8.28 | 0.007 | 0.001 |
| | | | GFP | | 0.001 | 0.000 |
| | | | NFP | | **0.012** | **0.002** |
| | | MF | 10-NN | 5.65 | 0.005 | 0.000 |
| | | | GFP | | **0.011** | 0.000 |
| | | | NFP | | **0.011** | **0.002** |
| | | CC | 10-NN | 5.12 | 0.000 | 0.000 |
| | | | GFP | | 0.019 | 0.002 |
| | | | NFP | | **0.058** | **0.011** |

#### S3.7 Diversity of predictions

In this section, we provide the results on the number of OG families that obtained at least one prediction at precision levels 0.5 and 0.8 for GO functions of different generality and for 10-NN, GFP and NFP classifier.

**Table S5. Number of OGs receiving at least one prediction on prokaryotic dataset.**

| Method | Number of OGs with at least one predicted GO function | Number of predictions of functions with $IC \in <2,4]$ | Number of OGs with at least one predicted GO function with IC>4 |
|---|---|---|---|

| | Precision: 0.5 | Precision: 0.8 | Precision: 0.5 | Precision: 0.8 | Precision: 0.5 | Precision: 0.8 |
|---|---|---|---|---|---|---|
| 10-NN | **3475/3475** | 461/3475 | 924/3475 | 187/3475 | 705/3475 | 243/3475 |
| GFP | **3475/3475** | 2007/3475 | 2176/3475 | 606/3475 | **1538/3475** | 581/3475 |
| NFP | **3475/3475** | **2536/3475** | **2390/3475** | **849/3475** | 1443/3475 | **622/3475** |

**Table S6. Number of OGs receiving at least one prediction on fungi dataset**

| Method | Number of OGs with at least one predicted GO function | | Number of predictions of functions with $IC \in <2,4]$ | | Number of OGs with at least one predicted GO function with IC>4 | |
|---|---|---|---|---|---|---|
| | Precision: 0.5 | Precision: 0.8 | Precision: 0.5 | Precision: 0.8 | Precision: 0.5 | Precision: 0.8 |
| 10-NN | 15404/15741 | 95/15741 | 17/15741 | 0/15741 | 262/15741 | 95/15741 |
| GFP | 15716/15741 | **5227/15741** | 3/15741 | 2/15741 | 0/15741 | 0/15741 |
| NFP | **15741/15741** | 4652/15741 | **1159/15741** | **129/15741** | **908/15741** | **172/15741** |

**Table S7. Number of OGs receiving at least one prediction on metazoa dataset**

| Method | Number of OGs with at least one predicted GO function | | Number of predictions of functions with $IC \in <2,4]$ | | Number of OGs with at least one predicted GO function with IC>4 | |
|---|---|---|---|---|---|---|
| | Precision: 0.5 | Precision: 0.8 | Precision: 0.5 | Precision: 0.8 | Precision: 0.5 | Precision: 0.8 |
| 10-NN | **9185/9185** | 61/9185 | 417/9185 | 23/9185 | **279/9185** | 50/9185 |
| GFP | **9185/9185** | 217/9185 | 1589/9185 | 8/9185 | 132/9185 | 19/9185 |
| NFP | **9185/9185** | **963/9185** | **4835/9185** | **119/9185** | 247/9185 | **64/9185** |

The results provided show that the NFP approach mostly provides more versatile predictions then baseline classifiers.

#### S3.8 Evaluation on model organisms

In this section, we present the evaluation results of k-NN, Gaussian Field Propagation and Neighborhoods function profiles approach on several selected prokaryotic, fungi and metazoa model organisms. We show the number of newly predicted genes with a given precision threshold and Information Content for each method.

**Table S8**. Number of predicted annotations (gene-function pairs - #OA), new annotations (#NA), and previously non-annotated genes that got at least one annotation (#NPG) on different model Prokaryotic, Fungi and Metazoa organisms for k-NN, Gaussian Field Propagation and Neighbourhood function profiles methods. The number of predicted annotations is computed for precision thresholds (Pr. tr.) 0.5 and 0.8. The #OA denotes the number of annotations present in the dataset. Counted GO functions have IC>4.

**Prokaryotes**

| Organism | Method | Pr. tr. | #OA | #PA | #NA | #NPG |
|---|---|---|---|---|---|---|
| *Pseudomonas aeruginosa* | 1-NN | 0.5 | 31913 | 2050 | 1017 | 301 |
| | 3-NN | | | 5416 | 1939 | 380 |
| | 10-NN | | | 6188 | 2646 | 495 |
| | GFP | | | 15505 | 8674 | 1398 |
| | NFP | | | 21227 | 12165 | 1351 |
| | 1-NN | 0.8 | 31913 | 11 | 2 | 2 |
| | 3-NN | | | 1536 | 220 | 58 |
| | 10-NN | | | 1954 | 232 | 50 |
| | GFP | | | 5172 | 1160 | 277 |
| | NFP | | | 6504 | 1537 | 235 |
| *Escherichia coli* | 1-NN | 0.5 | 29632 | 1725 | 811 | 254 |
| | 3-NN | | | 4833 | 1750 | 363 |
| | 10-NN | | | 5629 | 2614 | 487 |
| | GFP | | | 13750 | 7572 | 1236 |
| | NFP | | | 18912 | 10559 | 1277 |
| | 1-NN | 0.8 | 29632 | 11 | 2 | 2 |
| | 3-NN | | | 1261 | 171 | 48 |
| | 10-NN | | | 1687 | 207 | 45 |
| | GFP | | | 4351 | 905 | 218 |

| | | | | | | |
|---|---|---|---|---|---|---|
| | NFP | | | 5491 | 1211 | 209 |
| *Bacillus subtilis* | 1-NN | 0.5 | 23428 | 1629 | 765 | 212 |
| | 3-NN | | | 4253 | 1437 | 278 |
| | 10-NN | | | 4697 | 1988 | 361 |
| | GFP | | | 11600 | 6176 | 975 |
| | NFP | | | 15263 | 8079 | 968 |
| | 1-NN | 0.8 | 23428 | 10 | 0 | 0 |
| | 3-NN | | | 1139 | 102 | 31 |
| | 10-NN | | | 1492 | 152 | 37 |
| | GFP | | | 3932 | 772 | 167 |
| | NFP | | | 4916 | 1061 | 158 |
| *Streptomyces coelicolor* | 1-NN | 0.5 | 31200 | 2059 | 1066 | 282 |
| | 3-NN | | | 4847 | 1534 | 311 |
| | 10-NN | | | 5668 | 2388 | 487 |
| | GFP | | | 13542 | 7898 | 1298 |
| | NFP | | | 20052 | 10470 | 1339 |
| | 1-NN | 0.8 | 31200 | 14 | 0 | 0 |
| | 3-NN | | | 1614 | 138 | 40 |
| | 10-NN | | | 2234 | 247 | 48 |
| | GFP | | | 4386 | 1012 | 226 |
| | NFP | | | 5620 | 947 | 175 |
| *S. aureus* | 1-NN | 0.5 | 18517 | 1252 | 528 | 136 |
| | 3-NN | | | 3237 | 952 | 199 |
| | 10-NN | | | 3600 | 1374 | 247 |
| | GFP | | | 8656 | 4314 | 683 |
| | NFP | | | 11027 | 5386 | 712 |
| | 1-NN | | | 8 | 0 | 0 |

| | | | | | | |
|---|---|---|---|---|---|---|
| | 3-NN | 0.8 | 18517 | 918 | 84 | 21 |
| | 10-NN | | | 1373 | 114 | 27 |
| | GFP | | | 3143 | 603 | 135 |
| | NFP | | | 3535 | 586 | 119 |

**Fungi**

| Organism | Method | Pr. tr. | #OA | #PA | #NA | #NPG |
|---|---|---|---|---|---|---|
| *S. pombe* | 1-NN | 0.5 | 78377 | 1 | 1 | 1 |
| | 3-NN | | | 101 | 44 | 22 |
| | 10-NN | | | 142 | 87 | 29 |
| | GFP | | | 0 | 0 | 0 |
| | NFP | | | 464 | 282 | 67 |
| | 1-NN | 0.8 | 78377 | 0 | 0 | 0 |
| | 3-NN | | | 45 | 0 | 0 |
| | 10-NN | | | 12 | 0 | 0 |
| | GFP | | | 0 | 0 | 0 |
| | NFP | | | 25 | 0 | 0 |
| *S. cerevisiae* | 1-NN | 0.5 | 107013 | 4 | 3 | 3 |
| | 3-NN | | | 106 | 44 | 25 |
| | 10-NN | | | 139 | 89 | 39 |
| | GFP | | | 0 | 0 | 0 |
| | NFP | | | 757 | 552 | 148 |
| | 1-NN | 0.8 | 107013 | 0 | 0 | 0 |
| | 3-NN | | | 0 | 0 | 0 |
| | 10-NN | | | 45 | 0 | 0 |
| | GFP | | | 0 | 0 | 0 |
| | NFP | | | 71 | 3 | 2 |

| | | | | | | |
|---|---|---|---|---|---|---|
| *Aspergillus nidulans* | 1-NN | 0.5 | 242728 | 41 | 12 | 10 |
| | 3-NN | | | 2158 | 505 | 136 |
| | 10-NN | | | 5273 | 1817 | 452 |
| | GFP | | | 0 | 0 | 0 |
| | NFP | | | 12870 | 5683 | 805 |
| | 1-NN | 0.8 | 242728 | 0 | 0 | 0 |
| | 3-NN | | | 807 | 21 | 11 |
| | 10-NN | | | 933 | 46 | 20 |
| | GFP | | | 0 | 0 | 0 |
| | NFP | | | 2846 | 342 | 53 |
| *Neurospora crassa* | 1-NN | 0.5 | 91926 | 4 | 3 | 3 |
| | 3-NN | | | 274 | 151 | 52 |
| | 10-NN | | | 511 | 302 | 98 |
| | GFP | | | 0 | 0 | 0 |
| | NFP | | | 1113 | 786 | 139 |
| | 1-NN | 0.8 | 91926 | 0 | 0 | 0 |
| | 3-NN | | | 52 | 1 | 1 |
| | 10-NN | | | 25 | 3 | 2 |
| | GFP | | | 0 | 0 | 0 |
| | NFP | | | 143 | 28 | 4 |
| *Cryptococcus neoformans* | 1-NN | 0.5 | 75076 | 1 | 0 | 0 |
| | 3-NN | | | 58 | 45 | 24 |
| | 10-NN | | | 200 | 160 | 43 |
| | GFP | | | 0 | 0 | 0 |
| | NFP | | | 305 | 224 | 61 |
| | 1-NN | | | 0 | 0 | 0 |
| | 3-NN | | | 4 | 1 | 1 |

| | | | | | | |
|---|---|---|---|---|---|---|
| | 10-NN | 0.8 | 75076 | 12 | 0 | 0 |
| | GFP | | | 0 | 0 | 0 |
| | NFP | | | 14 | 0 | 0 |

Metazoa

| Organism | Method | Pr. tr. | #OA | #PA | #NA | #NPG |
|---|---|---|---|---|---|---|
| *Mus musculus* | 1-NN | 0.5 | 461620 | 0 | 0 | 0 |
| | 3-NN | | | 1279 | 900 | 348 |
| | 10-NN | | | 1875 | 1391 | 531 |
| | GFP | | | 1233 | 447 | 244 |
| | NFP | | | 2101 | 1060 | 484 |
| | 1-NN | 0.8 | 461620 | 0 | 0 | 0 |
| | 3-NN | | | 249 | 25 | 14 |
| | 10-NN | | | 263 | 44 | 27 |
| | GFP | | | 193 | 18 | 10 |
| | NFP | | | 453 | 14 | 9 |
| *D. melanogaster* | 1-NN | 0.5 | 313360 | 0 | 0 | 0 |
| | 3-NN | | | 1234 | 561 | 200 |
| | 10-NN | | | 1786 | 1018 | 388 |
| | GFP | | | 1514 | 345 | 165 |
| | NFP | | | 3639 | 1534 | 582 |
| | 1-NN | 0.8 | 313360 | 0 | 0 | 0 |
| | 3-NN | | | 461 | 8 | 5 |
| | 10-NN | | | 356 | 33 | 16 |
| | GFP | | | 475 | 10 | 5 |
| | NFP | | | 994 | 65 | 35 |
| | 1-NN | | | 0 | 0 | 0 |

| | | | | | | |
|---|---|---|---|---|---|---|
| *Homo sapiens* | 3-NN | 0.5 | 936678 | 2428 | 1601 | 686 |
| | 10-NN | | | 3771 | 2655 | 1088 |
| | GFP | | | 2349 | 921 | 484 |
| | NFP | | | 3633 | 1902 | 866 |
| | 1-NN | 0.8 | 936678 | 0 | 0 | 0 |
| | 3-NN | | | 646 | 57 | 33 |
| | 10-NN | | | 632 | 55 | 32 |
| | GFP | | | 269 | 40 | 24 |
| | NFP | | | 765 | 37 | 25 |
| *C. elegans* | 1-NN | 0.5 | 16614 | 0 | 0 | 0 |
| | 3-NN | | | 23 | 20 | 9 |
| | 10-NN | | | 45 | 43 | 15 |
| | GFP | | | 42 | 20 | 11 |
| | NFP | | | 66 | 34 | 8 |
| | 1-NN | 0.8 | 16614 | 0 | 0 | 0 |
| | 3-NN | | | 0 | 0 | 0 |
| | 10-NN | | | 1 | 1 | 1 |
| | GFP | | | 19 | 3 | 1 |
| | NFP | | | 24 | 4 | 2 |

**Table S9.** Number of predicted annotations (gene-function pairs - #OA), new annotations (#NA), and previously non-annotated genes that got at least one annotation (#NPG) on different model Prokaryotic, Fungi and Metazoa organisms for k-NN, Gaussian Field Propagation and Neighbourhood function profiles methods. The number of predicted annotations is computed for precision thresholds (Pr. tr.) 0.5 and 0.8. The #OA denotes the number of annotations present in the dataset. Counted GO functions have 2<IC≤4.

| **Prokaryotes** |
|---|

| Organism | Method | Pr. tr. | #OA | #PA | #NA | #NPG |
|---|---|---|---|---|---|---|
| *Pseudomonas aeruginosa* | 1-NN | 0.5 | 41583 | 0 | 0 | 0 |
| | 3-NN | | | 3332 | 1324 | 453 |
| | 10-NN | | | 4190 | 1736 | 684 |
| | GFP | | | 14907 | 7957 | 1926 |
| | NFP | | | 38596 | 21007 | 2294 |
| | 1-NN | 0.8 | 41583 | 0 | 0 | 0 |
| | 3-NN | | | 284 | 35 | 24 |
| | 10-NN | | | 1128 | 149 | 57 |
| | GFP | | | 3802 | 929 | 289 |
| | NFP | | | 9042 | 2323 | 419 |
| *Escherichia coli* | 1-NN | 0.5 | 35852 | 0 | 0 | 0 |
| | 3-NN | | | 2941 | 1185 | 411 |
| | 10-NN | | | 3876 | 1748 | 622 |
| | GFP | | | 12944 | 6970 | 1692 |
| | NFP | | | 32809 | 18165 | 2048 |
| | 1-NN | 0.8 | 35852 | 0 | 0 | 0 |
| | 3-NN | | | 254 | 29 | 23 |
| | 10-NN | | | 969 | 141 | 53 |
| | GFP | | | 3031 | 694 | 232 |
| | NFP | | | 7491 | 1942 | 367 |
| *Bacillus subtilis* | 1-NN | 0.5 | 29449 | 0 | 0 | 0 |
| | 3-NN | | | 2363 | 1007 | 324 |
| | 10-NN | | | 3046 | 1288 | 488 |
| | GFP | | | 10285 | 5378 | 1276 |
| | NFP | | | 26911 | 13999 | 1573 |
| | 1-NN | | | 0 | 0 | 0 |

| | | | | | | |
|---|---|---|---|---|---|---|
| | 3-NN | 0.8 | 29449 | 167 | 20 | 14 |
| | 10-NN | | | 826 | 108 | 46 |
| | GFP | | | 2738 | 585 | 179 |
| | NFP | | | 6464 | 1824 | 287 |
| *Streptomyces coelicolor* | 1-NN | 0.5 | 44119 | 0 | 0 | 0 |
| | 3-NN | | | 3280 | 1392 | 481 |
| | 10-NN | | | 4590 | 2015 | 703 |
| | GFP | | | 14914 | 8363 | 1827 |
| | NFP | | | 41723 | 20624 | 2226 |
| | 1-NN | 0.8 | 44119 | 0 | 0 | 0 |
| | 3-NN | | | 339 | 25 | 18 |
| | 10-NN | | | 1322 | 187 | 66 |
| | GFP | | | 3577 | 954 | 263 |
| | NFP | | | 9106 | 1558 | 382 |
| *S. aureus* | 1-NN | 0.5 | 21553 | 0 | 0 | 0 |
| | 3-NN | | | 1861 | 639 | 223 |
| | 10-NN | | | 2497 | 941 | 363 |
| | GFP | | | 8221 | 4029 | 904 |
| | NFP | | | 19105 | 9431 | 1125 |
| | 1-NN | 0.8 | 21553 | 0 | 0 | 0 |
| | 3-NN | | | 154 | 18 | 12 |
| | 10-NN | | | 755 | 89 | 34 |
| | GFP | | | 2319 | 494 | 151 |
| | NFP | | | 4413 | 881 | 184 |

**Fungi**

| Organism | Method | Pr. tr. | #OA | #PA | #NA | #NPG |
|---|---|---|---|---|---|---|

| | | | | | | |
|---|---|---|---|---|---|---|
| *S. pombe* | 1-NN | 0.5 | 69483 | 0 | 0 | 0 |
| | 3-NN | | | 4 | 2 | 2 |
| | 10-NN | | | 27 | 11 | 6 |
| | GFP | | | 0 | 0 | 0 |
| | NFP | | | 550 | 265 | 130 |
| | 1-NN | 0.8 | 69483 | 0 | 0 | 0 |
| | 3-NN | | | 0 | 0 | 0 |
| | 10-NN | | | 5 | 1 | 1 |
| | GFP | | | 0 | 0 | 0 |
| | NFP | | | 9 | 6 | 2 |
| *S. cerevisiae* | 1-NN | 0.5 | 92501 | 0 | 0 | 0 |
| | 3-NN | | | 9 | 4 | 4 |
| | 10-NN | | | 35 | 20 | 15 |
| | GFP | | | 0 | 0 | 0 |
| | NFP | | | 1307 | 637 | 216 |
| | 1-NN | 0.8 | 92501 | 0 | 0 | 0 |
| | 3-NN | | | 0 | 0 | 0 |
| | 10-NN | | | 3 | 1 | 1 |
| | GFP | | | 0 | 0 | 0 |
| | NFP | | | 14 | 4 | 2 |
| *Aspergillus nidulans* | 1-NN | 0.5 | 223748 | 0 | 0 | 0 |
| | 3-NN | | | 14 | 4 | 4 |
| | 10-NN | | | 163 | 45 | 25 |
| | GFP | | | 5 | 1 | 1 |
| | NFP | | | 6422 | 3474 | 841 |
| | 1-NN | | | 0 | 0 | 0 |
| | 3-NN | | | 0 | 0 | 0 |

| | | | | | | |
|---|---|---|---|---|---|---|
| | 10-NN | 0.8 | 223748 | 27 | 2 | 2 |
| | GFP | | | 4 | 0 | 0 |
| | NFP | | | 587 | 132 | 78 |
| *Neurospora crassa* | 1-NN | 0.5 | 89349 | 0 | 0 | 0 |
| | 3-NN | | | 6 | 3 | 3 |
| | 10-NN | | | 34 | 17 | 10 |
| | GFP | | | 0 | 0 | 0 |
| | NFP | | | 711 | 375 | 164 |
| | 1-NN | 0.8 | 89349 | 0 | 0 | 0 |
| | 3-NN | | | 0 | 0 | 0 |
| | 10-NN | | | 3 | 1 | 1 |
| | GFP | | | 0 | 0 | 0 |
| | NFP | | | 10 | 3 | 1 |
| *Cryptococcus neoformans* | 1-NN | 0.5 | 70685 | 0 | 0 | 0 |
| | 3-NN | | | 6 | 4 | 4 |
| | 10-NN | | | 24 | 11 | 6 |
| | GFP | | | 0 | 0 | 0 |
| | NFP | | | 595 | 307 | 140 |
| | 1-NN | 0.8 | 70685 | 0 | 0 | 0 |
| | 3-NN | | | 0 | 0 | 0 |
| | 10-NN | | | 5 | 1 | 1 |
| | GFP | | | 0 | 0 | 0 |
| | NFP | | | 11 | 6 | 2 |

**Metazoa**

| Organism | Method | Pr. tr. | #OA | #PA | #NA | #NPG |
|---|---|---|---|---|---|---|
| *Mus musculus* | 1-NN | 0.5 | 543059 | 0 | 0 | 0 |
| | 3-NN | | | 978 | 493 | 60 |
| | 10-NN | | | 2545 | 1450 | 493 |
| | GFP | | | 9873 | 5141 | 3797 |
| | NFP | | | 30558 | 16158 | 9860 |
| | 1-NN | 0.8 | 543059 | 0 | 0 | 0 |
| | 3-NN | | | 0 | 0 | 0 |
| | 10-NN | | | 258 | 49 | 16 |
| | GFP | | | 19 | 0 | 0 |
| | NFP | | | 1089 | 135 | 42 |
| *D. melanogaster* | 1-NN | 0.5 | 334632 | 0 | 0 | 0 |
| | 3-NN | | | 640 | 236 | 35 |
| | 10-NN | | | 3686 | 2655 | 463 |
| | GFP | | | 6260 | 2982 | 2205 |
| | NFP | | | 21579 | 10834 | 6022 |
| | 1-NN | 0.8 | 334632 | 0 | 0 | 0 |
| | 3-NN | | | 0 | 0 | 0 |
| | 10-NN | | | 174 | 23 | 8 |
| | GFP | | | 29 | 0 | 0 |
| | NFP | | | 2129 | 332 | 40 |
| *Homo sapiens* | 1-NN | 0.5 | 1081545 | 0 | 0 | 0 |
| | 3-NN | | | 1688 | 611 | 108 |
| | 10-NN | | | 4662 | 2555 | 994 |
| | GFP | | | 18854 | 9767 | 7045 |
| | NFP | | | 58041 | 30383 | 19159 |
| | 1-NN | | | 0 | 0 | 0 |

| | | | | | | |
|---|---|---|---|---|---|---|
| | 3-NN | 0.8 | 1081545 | 0 | 0 | 0 |
| | 10-NN | | | 443 | 44 | 17 |
| | GFP | | | 44 | 0 | 0 |
| | NFP | | | 1959 | 311 | 78 |
| *C. elegans* | 1-NN | 0.5 | 18579 | 0 | 0 | 0 |
| | 3-NN | | | 32 | 31 | 3 |
| | 10-NN | | | 67 | 51 | 24 |
| | GFP | | | 422 | 186 | 125 |
| | NFP | | | 1069 | 479 | 324 |
| | 1-NN | 0.8 | 18579 | 0 | 0 | 0 |
| | 3-NN | | | 0 | 0 | 0 |
| | 10-NN | | | 0 | 0 | 0 |
| | GFP | | | 1 | 0 | 0 |
| | NFP | | | 5 | 0 | 0 |

#### S3.9 Evaluation on CAFA 2 challenge data

We have evaluated and compared the performance of Neighborhood function profiles approach with baseline (k-NN) method on a Cafa 2 challenge data (https://biofunctionprediction.org/cafa/). Since the Ontology version we used to perform our experiments (from January 2014.) does not contain annotations being tested in Cafa 2, we use out of bag predictions (obtained by Random Forest trained on gene functional Neighborhood features) and leave one out predictions from the k-NN approach obtained on our GO function set directly to evaluate performance on Cafa 2 gene evaluation set. Ontology version used (from December 2016.) to obtain predictions on Fungi and Metazoa data contained the annotations from Cafa 2, so we first removed all OGs associated to genes contained in the Cafa 2 challenge, then created gene functional Neighborhood and location Neighborhood features and trained Random forest and k-NN models.  All removed OGs were placed in a test set on which obtained models were evaluated. Since genes can be associated to multiple OGs, and our models predict functions for a set of selected OGs, we use average score obtained on all associated OGs as classifier score for a given gene. AUC and AUPRC measures have been computed on the obtained scores to evaluate these models on Cafa 2 challenge data.

The obtained results (see Figures S36.) show that Neighborhood function profiles approach (average AUC : 0.78, AUPRC: 0.207 - prokaryotes, AUC: 0.67, AUPRC: 0.065 - fungi, AUC: 0.72, AUPRC: 0.025 - metazoa) significantly outperforms the baseline method (average AUC : 0.62, AUPRC: 0.13 - prokaryotes, AUC: 0.52, AUPRC: 0.055 - fungi, AUC: 0.53, AUPRC: 0.0195 - metazoa). The corresponding p-values, as computed by Mann-Whitney U test are ($p_{AUC} < 2.2 \cdot 10^{-16}, p_{AUPRC} < 2.2 \cdot 10^{-16}$ - prokaryotes, $p_{AUC} < 2.2 \cdot 10^{-16}, p_{AUPRC} = 5.28 \cdot 10^{-16}$ - fungi, $p_{AUC} < 2.2 \cdot 10^{-16}, p_{AUPRC} < 2.2 \cdot 10^{-16}$ - metazoa). It can also be shown that the AUC scores obtained on Cafa validation data have significant Pearson and Spearman [PearsonCorr,Spearman] correlation to obtained out of bag and leave one out validation scores obtained on the initial datasets ( $r = 0.38, p_{Pearson,GFN} < 2.2 \cdot 10^{-16}, \rho = 0.37, p_{Spearman,GFN} < 2.2 \cdot 10^{-16}$, $r = 0.39, p_{Pearson,Baseline} < 2.2 \cdot 10^{-16}, \rho = 0.38, p_{Spearman,Baseline} < 2.2 \cdot 10^{-16}$, - prokaryotes, $r = 0.09, p_{Pearson,GFN} = 0.002, \rho = 0.095, p_{Spearman,GFN} = 0.001$, $r = 0.15, p_{Pearson,Baseline} = 2.3 \cdot 10^{-7}, \rho = 0.02, p_{Spearman,Baseline} = 0.43$, - fungi, $r = 0.093, p_{Pearson,GFN} = 4.5 \cdot 10^{-5}, \rho = 0.12, p_{Spearman,GFN} = 2.76 \cdot 10^{-7}$, $r = 0.196, p_{Pearson,Baseline} < 2.2 \cdot 10^{-16}, \rho = 0.25, p_{Spearman,Baseline} < 2.2 \cdot 10^{-16}$, - metazoa).

See also Figure S37.

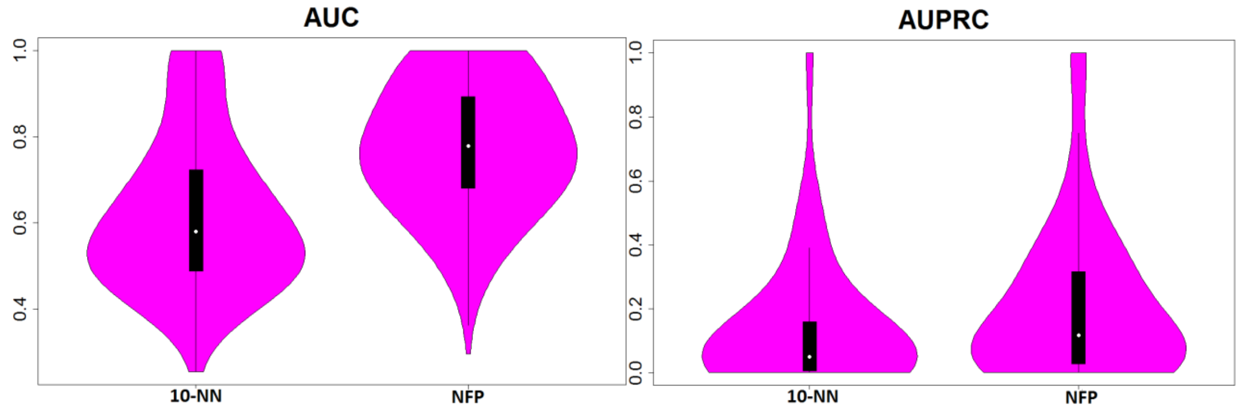

a) Prokaryotic organisms

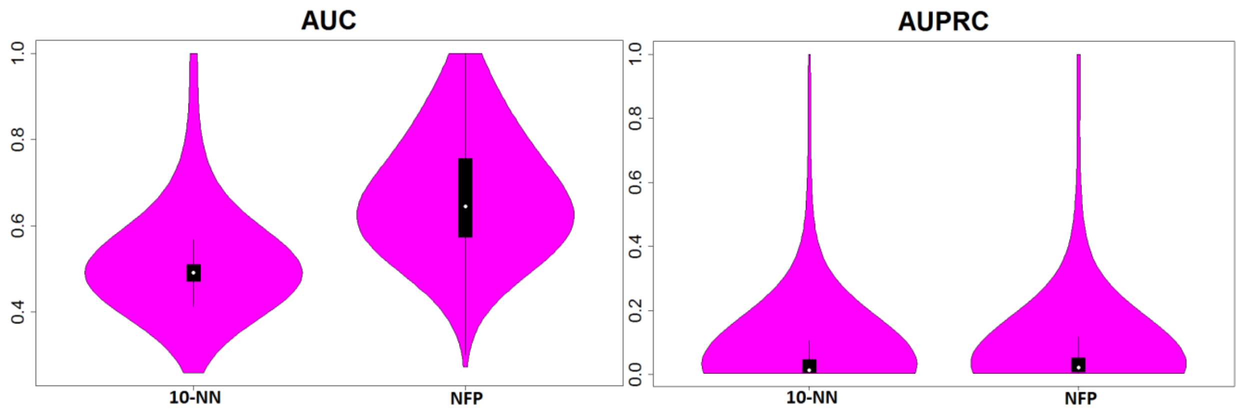

b) Fungi dataset

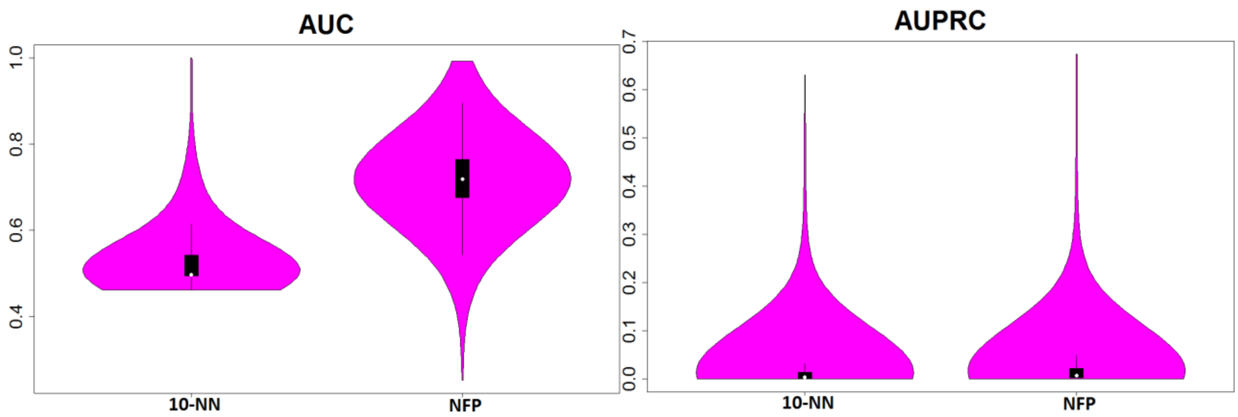

c) Metazoa dataset

**Figure S36.** Comparative performance results of 10-NN and Neighborhood function profiles on the CAFA2 gene validation sets for Prokaryotic a), Fungi b) and Metazoa c) organisms.

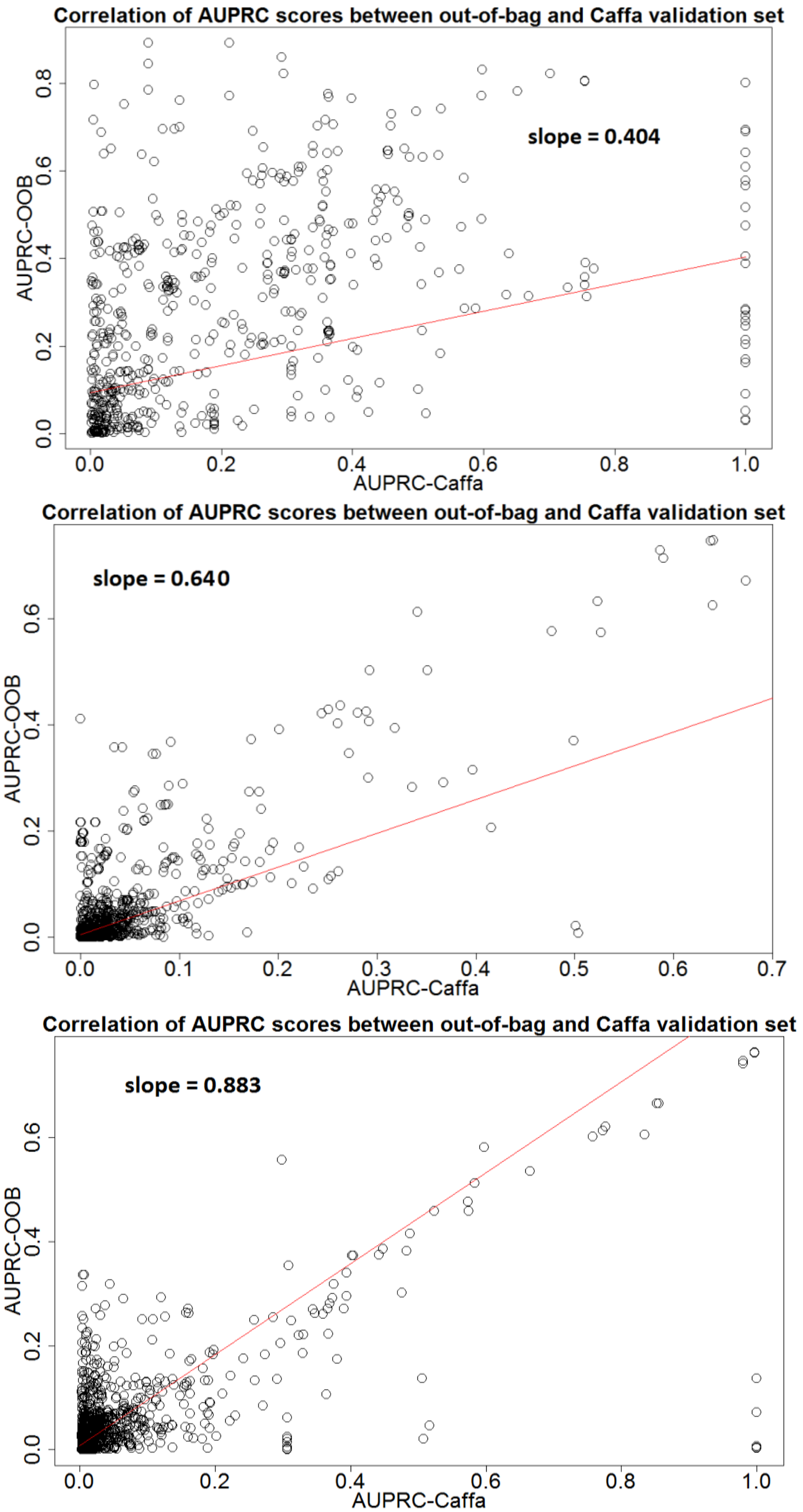

**Figure S37.** Correlation between out-of-bag and Caffa set for the NFP approach computed on Prokaryotic dataset (top), Fungi dataset (middle) and Metazoa dataset (bottom).

#### S3.10 Gene functional annotations from three different GO namespaces complement each other

Throughout our work, we use a set of 1048 GO functions from all three namespaces (Biological process - BP, Molecular function - MF and Cellular component - CC) to train our model for gene function prediction on prokaryotic organisms. In this section, we explore the complementarity of functional annotations contained in these namespaces. The main goal is to explore if adding (non-circular) annotations from two namespaces to the annotations of one selected namespace increases prediction performance of Neighborhood function profiles approach. To fully understand the influence of (non-circular) annotations from different namespaces to gene function prediction performance, we compare the obtained results with the result of NFP approach on a dataset containing annotations of the selected namespace extended with randomly generated features. We generate equal number of random features as the number of non-circular annotations from remaining two namespaces used in the original experiment. This experimental setup demonstrate in what amount (non-circular) annotations from other namespaces complement the information contained in the target namespace and what is the benefit of using this information for gene function prediction in the NFP approach. The level of circularity of an annotation is measured with Jaccard index [Jaccard]. We test the performance gain with different Jaccard index thresholds: 0.0, 0.1, 0.6 and 1.0. The Jaccard index 0.0 disallows adding any GO annotation from complementing namespaces such that there exists any annotation from selected namespace sharing at least one OG. On the other hand, Jaccard index 1.0 allows all GO annotations from complementing namespaces to be used in model training. For practical purposes, using Jaccard index 1.0 allows adding all functions from the complementing namespaces to be used for model training. We compute the Mann - Whitney U test of statistical significance of difference of the mean between two sets of predictions (containing annotations from selected namespace and complementary annotations vs containing annotations from selected namespace and randomly generated annotations in the same range) and show comparative violin plots.

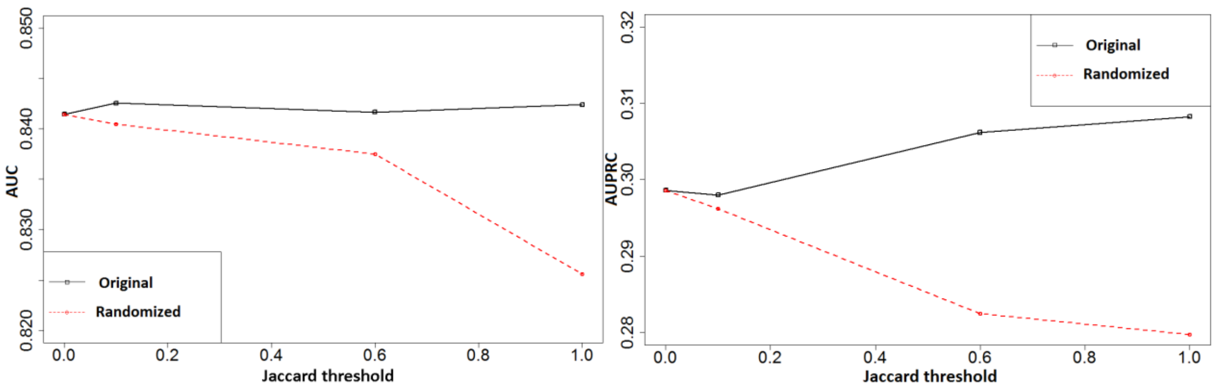

a) Biological process

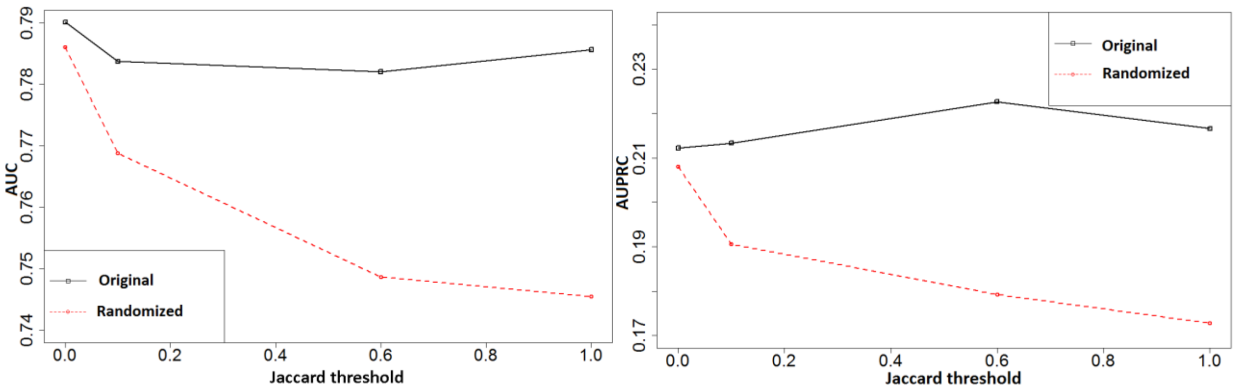

b) Molecular function

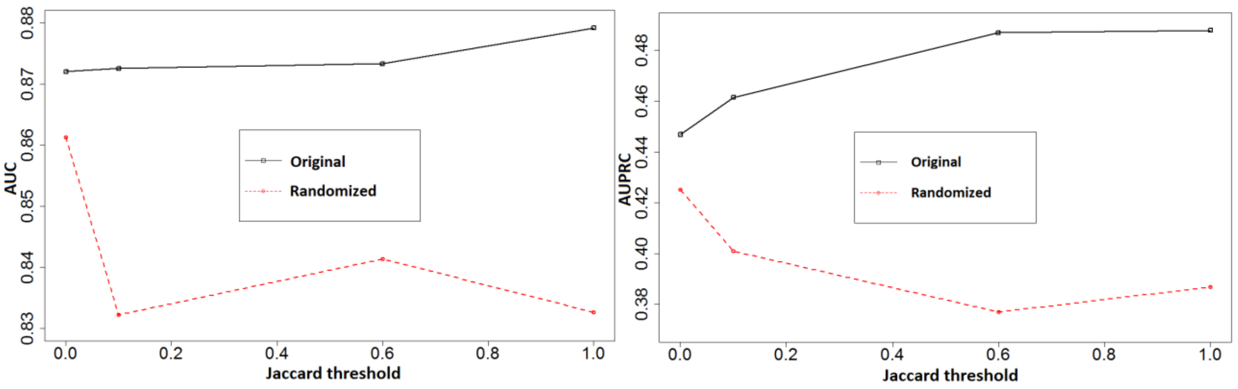

c) Cellular component

**Figure S38.** Neighborhood function profiles performance measured with AUC and AUPRC measures for different values of Jaccard index threshold.

Results presented in Figure S38. show that classifier performance mostly increases with the increase of the Jaccard index threshold. This is to be expected since more permissive threshold allows adding features derived from more complementing functions. At the same time, replacing these features with randomly generated features degrades the classifier performance (red line). This demonstrates the usefulness of using complementing namespaces in gene function prediction with Neighborhood function profiles method.

The difference in the mean of AUC and AUPRC across functions when predicted using Biological process features and the complementing features compared to Biological process features and randomized features (equal number to the complementing features) is not significant (according to one-sided Mann-Whitney U test) for very strict Jaccard index level (0.0 and 0.1), however mean of AUPRC scores is statistically significantly higher ($p = 0.023$) when using complementing features for the level 0.6 and statistically significantly higher for both the AUC ($p = 0.009$) and AUPRC ($p = 0.0102$) for Jaccard threshold 1.0.

The difference in the mean of AUC and AUPRC across functions when predicted using Molecular function features and the complementing features compared to Molecular function features and randomized features (equal number to the complementing features) is not significant (according to one-sided Mann-Whitney U test) for very strict Jaccard index level (0.0), however mean of AUC scores ($p = 0.024$ ) and the mean of AUPRC scores ($p = 0.029$) is statistically significantly higher when using complementing features for the Jaccard threshold level 0.1, AUC (p = $5 \cdot 10^{-4}$), AUPRC (p =$2.5 \cdot 10^{-4}$) for threshold 0.6 and AUC (p = $8.2 \cdot 10^{-6}$ ), AUPRC (p = $5.5 \cdot 10^{-5}$) for threshold 1.0.

The difference in the mean of AUC and AUPRC across functions when predicted using Cellular component features and the complementing features compared to Cellular component features and randomized features (equal number to the complementing features) is not significant (according to one-sided Mann-Whitney U test) for very strict Jaccard index level (0.0), however mean of AUC scores (p = 0.023 ) and the mean of AUPRC scores (p = 0.026) is statistically significantly higher when using complementing features for the Jaccard threshold level 0.1, AUC (p = 0.035), AUPRC (p = 0.004) for threshold 0.6 and AUC (p = 0.0105 ), AUPRC (p = 0.0053) for threshold 1.0.

As can be seen, prediction of gene functions for all namespaces can be improved by introducing functions from the complementary namespaces. Higher level of allowed redundancy significantly increases the performance gain.

#### S3.11 Neighborhood function profiles improve prediction of conditional growth defects in different *E. coli* strains

The aim of this experiment was to improve the prediction of conditional growth defects for 696 different strains of E.Coli bacteria. Genomic data, disruption scores and conditional scores used to create datasets were obtained from [Galardini]. In addition to conditional scores, provided by the authors of [Galardini] and used to predict conditional growth defects, we created three additional datasets: a) functional gene Neighborhoods for each E.Coli strain. Features in this datasets are GO function - OG pairs, thus a feature vector contains frequency counts of GO function occurrence in the Neighborhood of each OG contained in the E.Coli strain. This resulted in large number of features, which were reduced by removing all pairs with 0 frequency in all strains. As a result, a dataset with 71540 features was obtained. b) We apply PCA [PearsonK] on the dataset obtained in a) and obtain 228 components, c) we combine the conditional score and PCA features in the combined dataset.

As a baseline predictor, we use conditional scores computed and used in [Galardini]. We notice that using Random Forest on datasets a) and b) does not yield better performance compared to the baseline predictor. Using Random Forest directly on the conditional scores dataset does not significantly increase the AUC score compared to baseline predictor (p=0.4538 using Mann-Whitney U test), however it significantly increases performance (measured using DeLong test [DeLong]) for 40 conditional growth defects (FDR<0.2) and has significantly worse performance for 10 conditional growth defects (FDR<0.2). Using Random Forest algorithm on dataset c) which combines conditional scores and PCA components significantly increases prediction performance (as measured by the AUC score, p=0.0109 as computed by the Mann-Whitney U test). Further analyses using DeLong test [DeLong] for ROC curve comparison revealed that 62 conditional growth defects were significantly more accurately predicted by the Random Forest classifier using dataset c) than the baseline predictor (FDR<0.2) whereas 9 conditional growth defects had significantly worse prediction using Random Forest on dataset c) than baseline predictor (FDR < 0.2).

We noticed complementarity in predictions between the baseline predictor and the Random Forest applied to dataset c), thus we created an Ensemble predictor that predicted conditional growth defects as follows: $Pr_{ensemble}(d_i) = (w_1 \cdot Pr_{baseline}(d_i) + w_2 \cdot Pr_{RF3}(d_i)), w_1 + w_2 = 1.$ $d_i$ are normalized scores obtained as follows: $d_i = c_i/(max\ c)$, where $c_i$ denotes the corresponding score of the original classifier.

It shows that using $w_1 = 0.1,\ w_2 = 0.9$ significantly improves accuracy (as measured by AUC) of both the Baseline predictor and PCABaselineRF predictor (see Figure x23). A good tradeoff for both AUC and AUPRC scores is achieved with these weights (see Figures x25, x26).

Ensemble classifier significantly improves predictions of conditional growth defects compared to the baseline predictor (p=$1.7 \cdot 10^{-15}$) and compared to the PCABaselineRF predictor (p=$1.6 \cdot 10^{-9}$), as measured by the one-sided Wilcox singed-rank test. It predicts 62 defects significantly better than baseline predictor (FDR<0.2) and 10 defects worse than baseline predictor (FDR<0.2).

Distributions of AUC scores obtained by baseline predictor, Random Forest applied to dataset b) and ensemble predictor are shown in Figures S39 and S40.

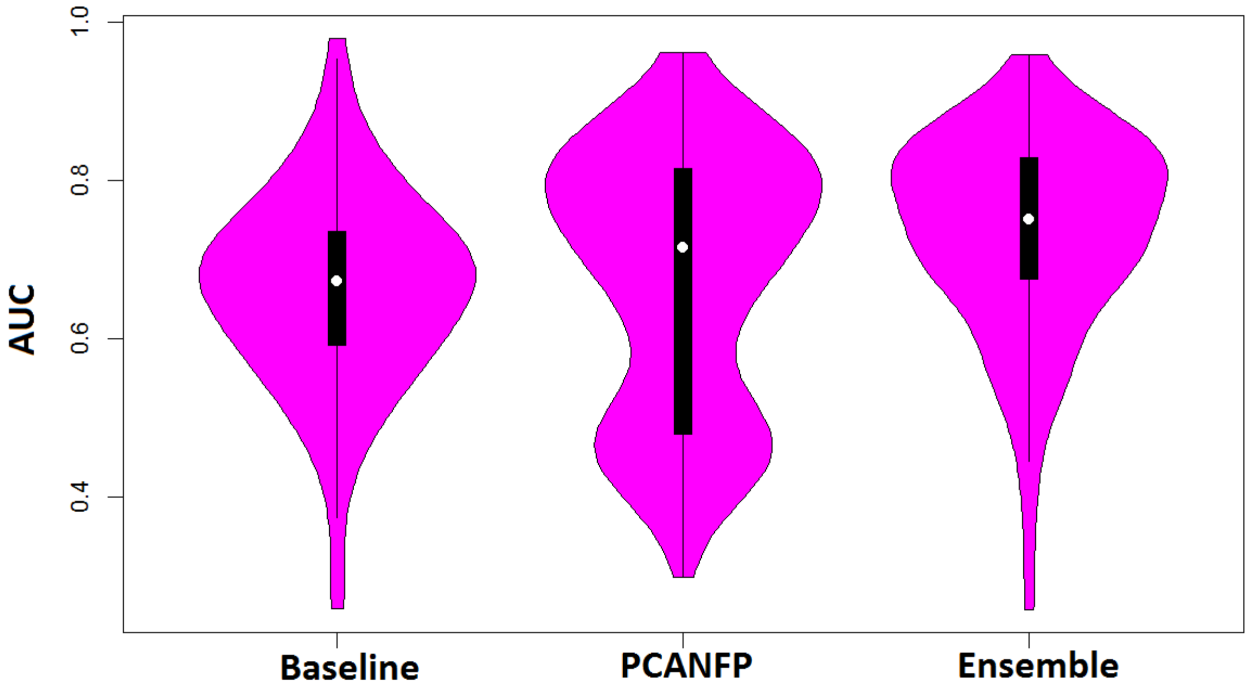

**Figure S39.** Distribution of AUC scores for the baseline classifier, Random Forest classifier applied to a dataset containing PCA components as features and the ensemble classifier.

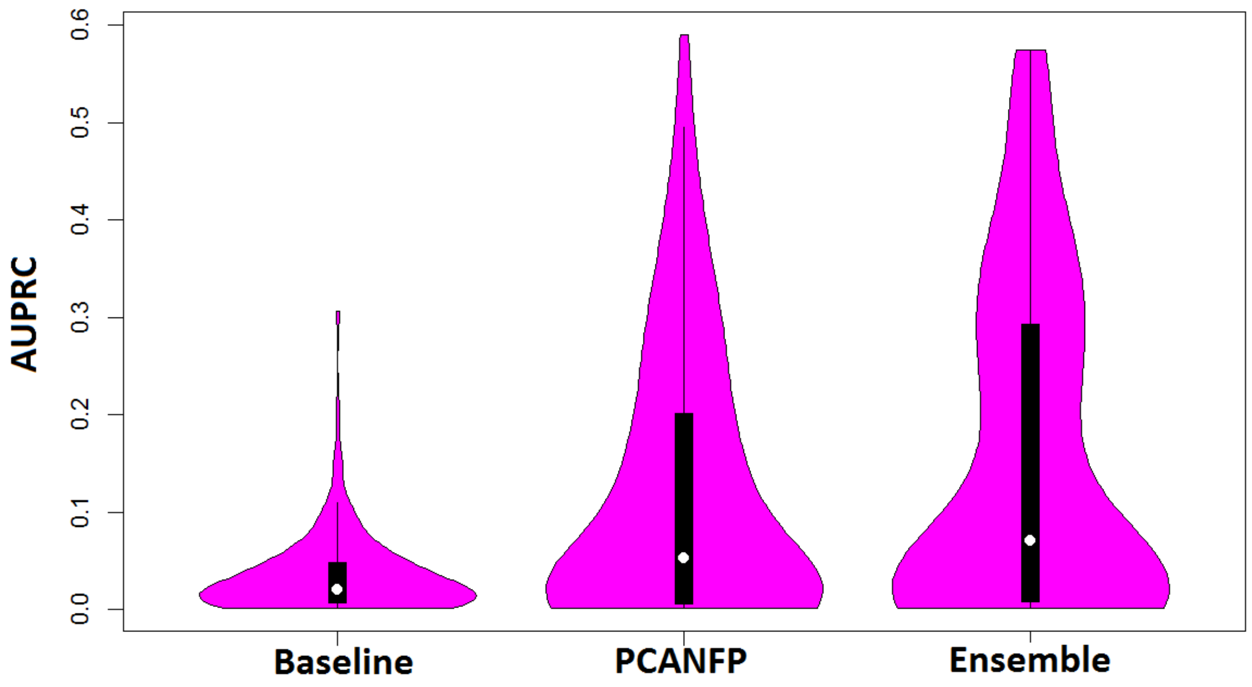

**Figure S40.** Distribution of AUPRC scores for the baseline classifier, Random Forest classifier applied to a dataset containing PCA components as features and the ensemble classifier.

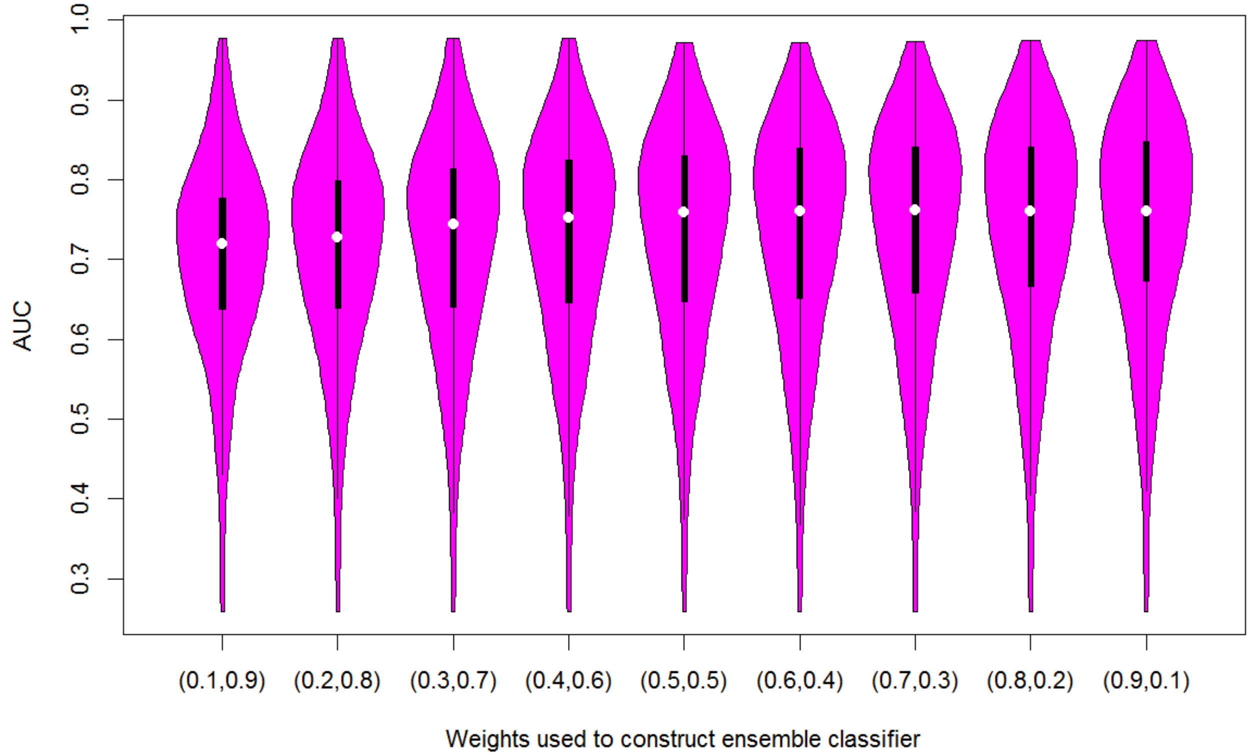

**Figure S41.** Distribution of AUC scores for different values of parameter $w_1, w_2$ in ensemble classifier.

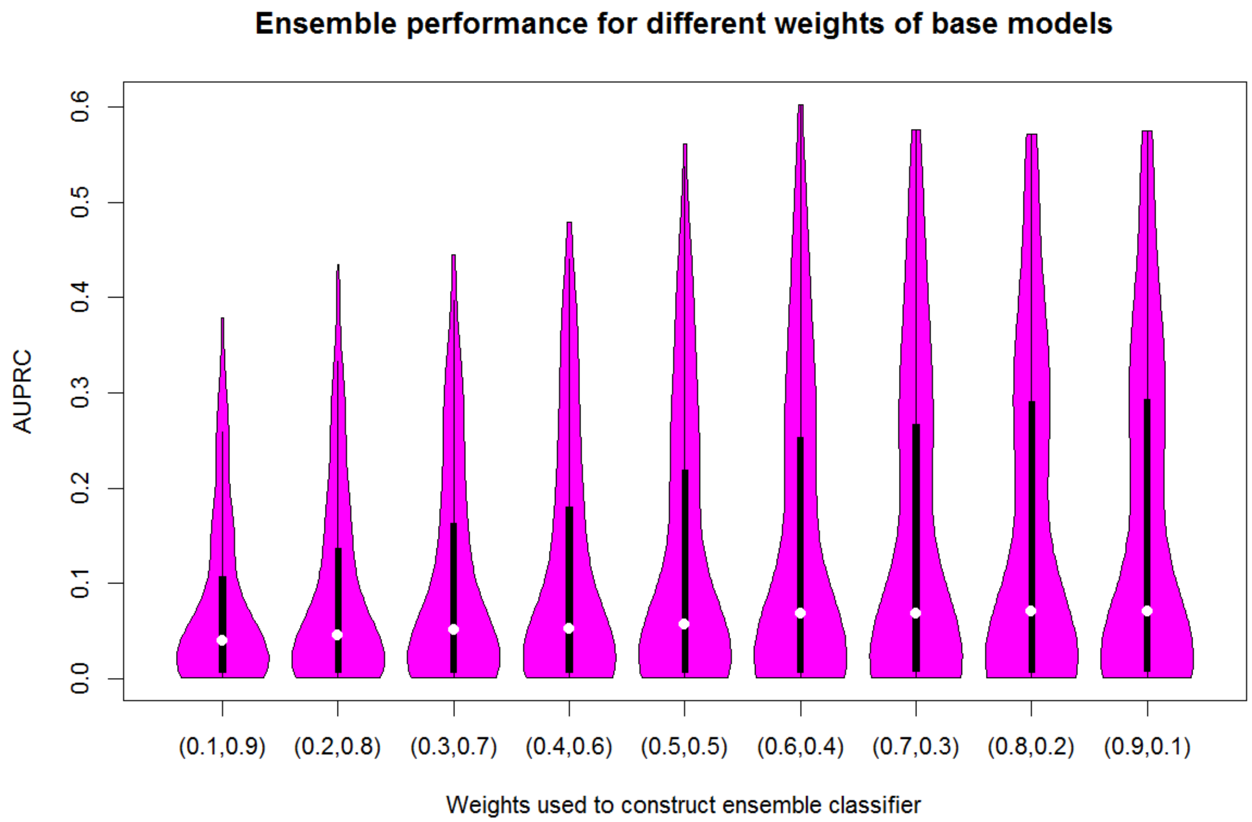

**Figure S42**. Distribution of AUC scores for different values of parameter $w_1, w_2$ in ensemble classifier.

Ensemble metaparameters $w_1, w_2$ were chosen based on the results demonstrated in Figures S41 and S42.

Area under ROC curve achieved by Baseline method and the ensemble method for the top phenotypes with the smallest corrected p-value (as measured by the De Long test and corrected for false discovery rate) of difference in AUC between approaches can be seen in Figure S43.

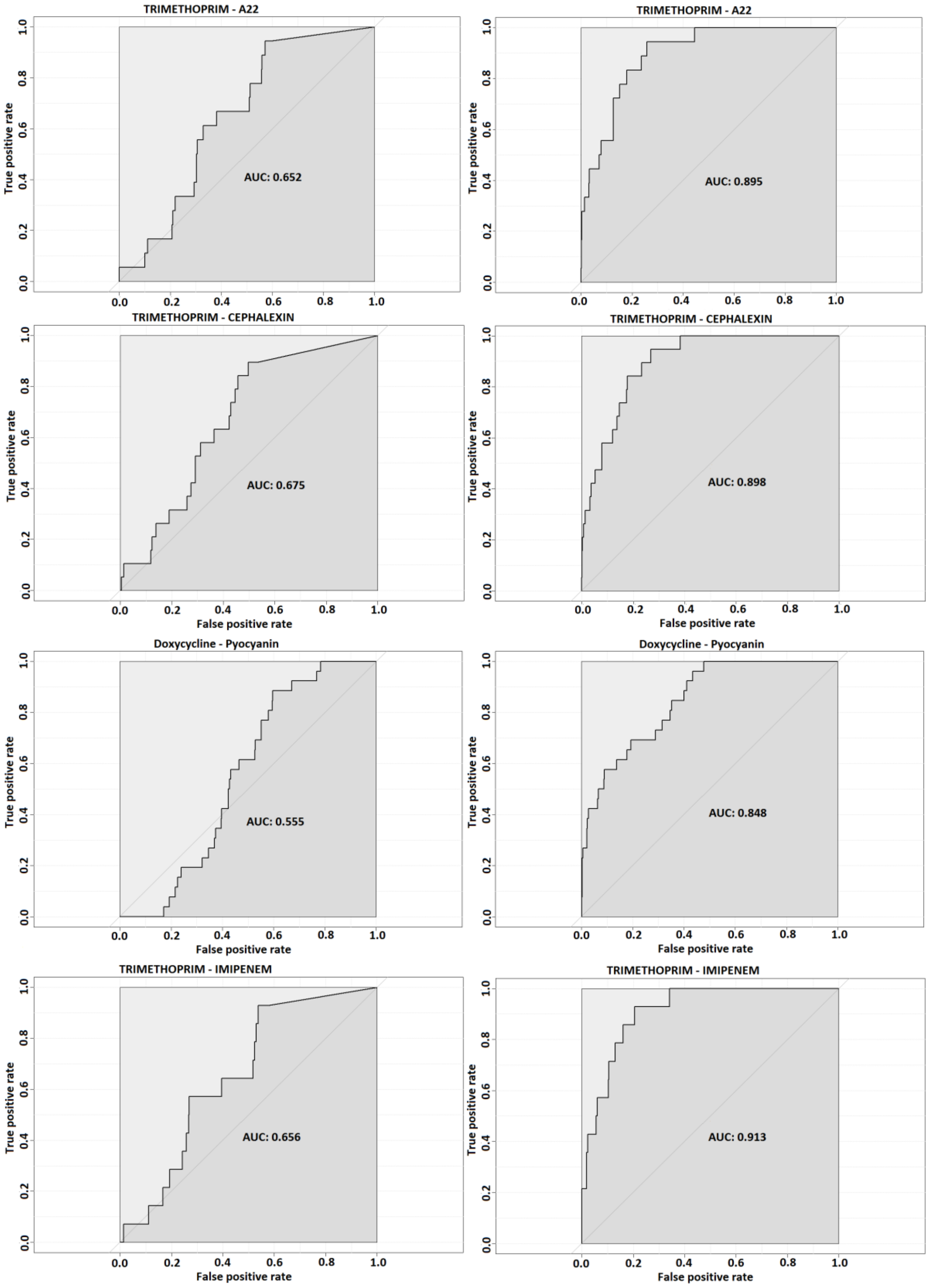

**Figure S43.** Area under ROC curve achieved by Baseline method (left) and the ensemble method (right) for the top phenotypes with the smallest corrected p-value (as measured by the De Long test and corrected for false discovery rate) of difference in AUC between approaches.

#### S3.12 Removing information about Operons and Enrichments significantly reduces NFP performance

We show that the performance of NFP model significantly decreases after removing information about operons in prokaryotes (which is expected), however it further significantly decreases when removing information about enrichments. For each OG, we replace the feature (GO function) that is enriched with at least one GO on the target side of this OG with a randomized version of this feature (values are generated randomly from uniform distribution on the interval [$minValue_{atx}$, $maxValue_{atx}$]).

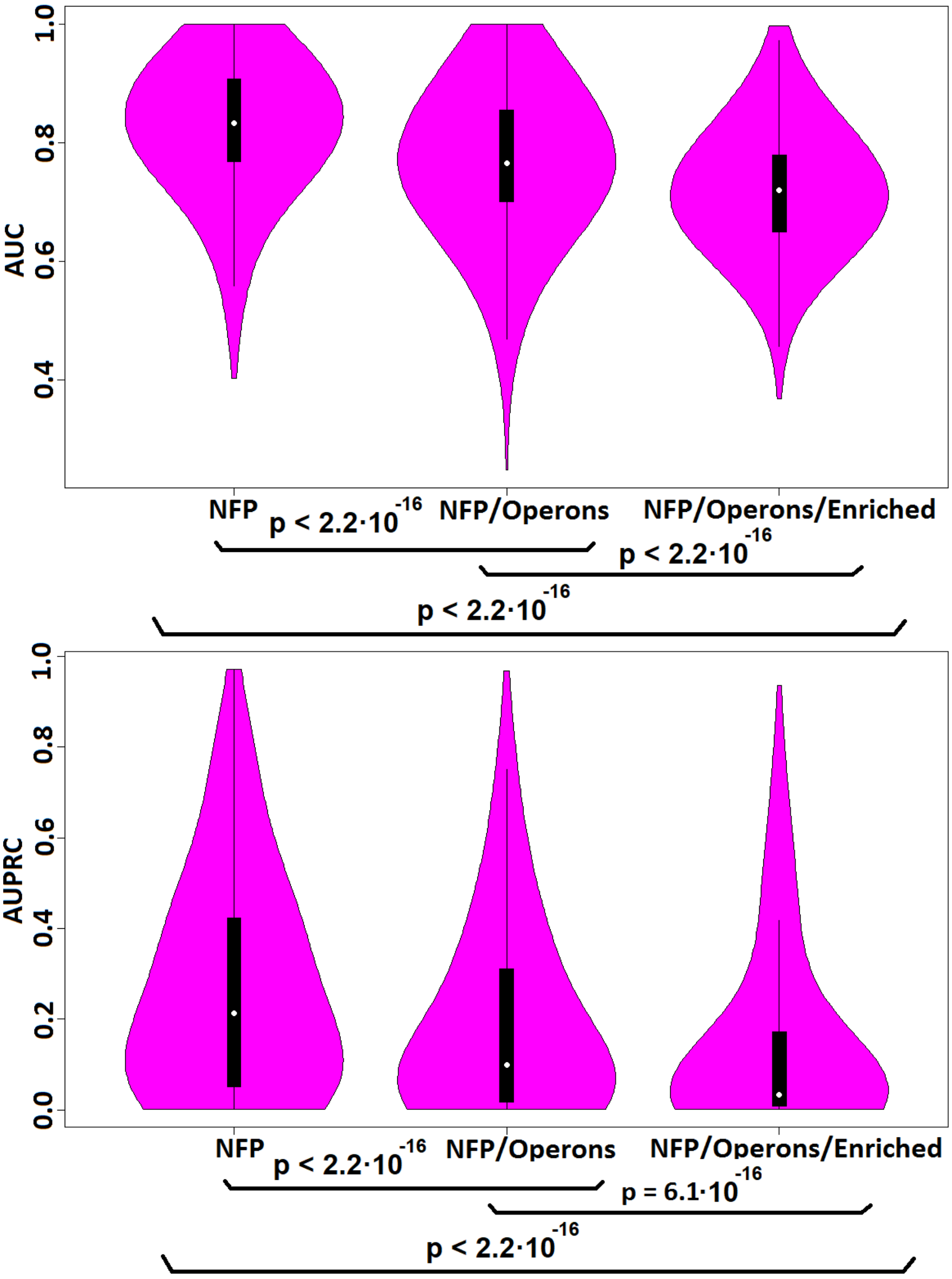

**Figure S44.** The decrease in AUC and AURPC in NFP method caused by excluding operons in feature computation and randomizing features that represent functions that are significantly enriched with at least one other GO contained in the set of target functions for some OG.
As it can be seen from Figure S44., the NFP model performance significantly decreases (as measured by one – sided Wilcoxon signed-rank test) when information about operons in prokaryotic organisms is omitted during feature construction. Further on, when features representing functions that have significant enrichment with at least one target variable are replaced with randomized attributes, further significant decrease in performance is detected.
